# Mechanism of a COVID-19 nanoparticle vaccine candidate that elicits a broadly neutralizing antibody response to SARS-CoV-2 variants

**DOI:** 10.1101/2021.03.26.437274

**Authors:** Yi-Nan Zhang, Jennifer Paynter, Cindy Sou, Tatiana Fourfouris, Ying Wang, Ciril Abraham, Timothy Ngo, Yi Zhang, Linling He, Jiang Zhu

## Abstract

Vaccines that induce potent neutralizing antibody (NAb) responses against emerging variants of severe acute respiratory syndrome coronavirus 2 (SARS-CoV-2) are essential for combating the coronavirus disease 2019 (COVID-19) pandemic. We demonstrated that mouse plasma induced by self-assembling protein nanoparticles (SApNPs) that present 20 rationally designed S2GΔHR2 spikes of the ancestral Wuhan-Hu-1 strain can neutralize the B.1.1.7, B.1.351, P.1, and B.1.617 variants with the same potency. The adjuvant effect on vaccine-induced immunity was investigated by testing 16 formulations for the multilayered I3-01v9 SApNP. Using single-cell sorting, monoclonal antibodies (mAbs) with diverse neutralization breadth and potency were isolated from mice immunized with the receptor binding domain (RBD), S2GΔHR2 spike, and SApNP vaccines. The mechanism of vaccine-induced immunity was examined in mice. Compared with the soluble spike, the I3-01v9 SApNP showed 6-fold longer retention, 4-fold greater presentation on follicular dendritic cell dendrites, and 5-fold stronger germinal center reactions in lymph node follicles.

**ONE-SENTENCE SUMMARY:** With a well-defined mechanism, spike nanoparticle vaccines can effectively counter SARS-CoV-2 variants.

## INTRODUCTION

The COVID-19 pandemic has led to more than 188 million infection cases and 4 million deaths globally. Antibody responses to SARS-CoV-2 spike antigens can be sustained for several months in most COVID-19 patients after infection (*1-4*). However, recently identified variants of concern (VOCs) exhibit higher transmissibility and resistance to prior immunity as SARS-CoV-2 continues to adapt to the human host (*5, 6*). One such variant, B.1.1.7 (WHO classification: Alpha), emerged from southeast England in October 2020 and accounted for two-thirds of new infections in London in December 2020, with a higher transmission rate (43-90%) and risk of mortality (32-104%) than previously circulating strains (*7, 8*). Other variants, such as B.1.351 (Beta) and P.1 (Gamma), also became prevalent in three provinces in South Africa and Manaus, Brazil, respectively (*6, 9, 10*). The B.1.617.2 (Delta) variant, which was initially identified in India, is becoming a dominant strain in many countries (*11, 12*) and responsible for the majority of new COVID-19 cases. This variant was found to be ∼60% more transmissible than the highly infectious B.1.1.7 variant (*12*). The rise of SARS-CoV-2 VOCs and their rapid spread worldwide result in more infection cases, hospitalizations, and potentially more deaths, further straining healthcare resources (*10*).

To date, eight COVID-19 vaccines have been approved for emergency use in humans, with more than 90 candidates assessed in various phases of clinical trials (*13*). With the exception of inactivated whole-virion vaccines, diverse platforms have been used to deliver the recombinant SARS-CoV-2 spike, such as mRNA-encapsulating liposomes (e.g., BNT162b2 and mRNA-1273), adenovirus vectors (e.g., ChAdOx1 nCoV-19 [AZD1222], CTII-nCoV, Sputnik V, and Ad26.COV2.S), and micelle-attached spikes (e.g., NVX-CoV2373). These vaccines demonstrated 65-96% efficacy in Phase 3 trials, with lower morbidity and mortality associated with COVID-19 disease (*14-19*). However, a notable loss of vaccine efficacy against new SARS-CoV-2 variants was reported, likely caused by spike mutations in the receptor-binding domain (RBD; e.g., K417N, E484K, and N501Y), N-terminal domain (NTD; e.g., L18F, D80A, D215G, and Δ242-244), and other regions that are critical to spike stability and function (e.g., D614G and P681R) (*6, 11, 20-25*). Among circulating VOCs, the B.1.351 lineage appeared to be most resistant to neutralization by convalescent plasma (9.4-fold) and vaccine sera (10.3- to 12.4-fold) (*26*), whereas a lesser degree of reduction was observed for an early variant, B.1.1.7 (*27-29*). Based on these findings, it was suggested that vaccines would need to be updated periodically to maintain protection against rapidly evolving SARS-CoV-2 (*30-32*). However, in a recent study, convalescent sera from B.1.351 or P.1-infected individuals showed a more visible reduction of B.1.617.2 neutralization than convalescent sera from individuals infected with early pandemic strains (*33*). Together, these issues raise the concern that herd immunity may be difficult to achieve, highlighting the necessity of developing vaccines that can elicit a broadly neutralizing antibody (bNAb) response to current and emerging variants (*25, 31*). As previously reported (*34-38*), the production of a bNAb response relies on long-lived germinal center (GC) reactions to activate precursor B cells, stimulate affinity maturation, and form long-term immune memory. In particular, antigen retention and presentation within lymph node follicles are key to the induction of long-lived GC reactions (*34, 36, 39*) and should be considered in the development of bNAb-producing vaccines (*40*).

We previously investigated the cause of SARS-CoV-2 spike metastability and rationally designed the S2GΔHR2 spike, which was displayed on three self-assembling protein nanoparticle (SApNP) platforms, including ferritin (FR) 24-mer and multilayered E2p and I3-01v9 60-mers, as COVID-19 vaccine candidates (*41*). In the present study, we investigated the vaccine-induced NAb response to SARS-CoV-2 VOCs and mechanism by which SApNP vaccines (e.g., I3-01v9) generate such a response. We first examined the neutralizing activity of mouse plasma from our previous study (*41*) against four representative SARS-CoV-2 variants, B.1.1.7, B.1.351, P.1, and B.1.617_Rec_, which was derived from an early analysis of the B.1.617 lineage (*11*) and shares key spike mutations with VOC B.1.617.2. Mouse plasma induced by the S2GΔHR2 spike-presenting I3-01v9 SApNP potently neutralized all four variants with comparable titers to the wildtype strain, Wuhan-Hu-1. When a different injection route was tested in mouse immunization, E2p and I3-01v9 SApNPs sustained neutralizing titers against the four variants, even at a low dosage of 3.3 μg, whereas a significant reduction of plasma neutralization was observed for the soluble spike. Next, we examined the adjuvant effect on vaccine-induced humoral and T-cell responses for the I3-01v9 SApNP. While detectable plasma neutralization was observed for the non-adjuvanted I3-01v9 group, conventional adjuvants, such as aluminum hydroxide (AH) and phosphate (AP), boosted the titers by 8.6- to 11.3-fold (or 9.6 to 12.3 times). Adjuvants that target the stimulator of interferon genes (STING) and Toll-like receptor 9 (TLR9) pathways enhanced neutralization by 21- to 35-fold, alone or combined with AP, in addition to a Th1-biased cellular response. We then performed antigen-specific single-cell sorting and isolated 20 monoclonal antibodies (mAbs) from RBD, spike, and I3-01v9 SApNP-immunized mice. These mAbs were derived from diverse B cell lineages, of which some neutralized the wildtype Wuhan-Hu-1 strain and four variants with equivalent potency. Lastly, we investigated how SApNPs behave in lymph nodes and induce GCs by characterizing vaccine delivery and immunological responses at the intraorgan, intracellular, and intercellular levels in mice. The I3-01v9 SApNP showed 6-fold longer retention, 4-fold greater presentation on follicular dendritic cell (DC) dendrites, and 5-fold higher GC reactions than the soluble spike.

Intact SApNPs in lymph node tissues could be visualized by transmission electron microscopy (TEM). Our study thus demonstrates that a spike-presenting SApNP vaccine derived from the “ancestral” SARS-CoV-2 strain may confer broad protection against emerging variants.

## RESULTS

### Spike and SApNP vaccine-induced neutralizing responses to SARS-CoV-2 variants

We previously demonstrated that the rationally designed S2GΔHR2 spike was more immunogenic than the S2P spike (*42*), and SApNPs displaying 8-20 spikes outperformed soluble spikes in NAb elicitation (*41*) **(Fig. 1A)**. Notably, the I3-01v9 SApNP that presents 20 S2GΔHR2 spikes induced a potent NAb response to both SARS-CoV-1 and SARS-CoV-2, as well as critically needed T-cell responses (*41*). Because SARS-CoV-1 shares only modest sequence similarity (∼73% in the RBD) with SARS-CoV-2, we hypothesized that our vaccines would protect against emerging variants that are much more closely related to the ancestral SARS-CoV-2 strain, Wuhan-Hu-1.

**Fig. 1.**
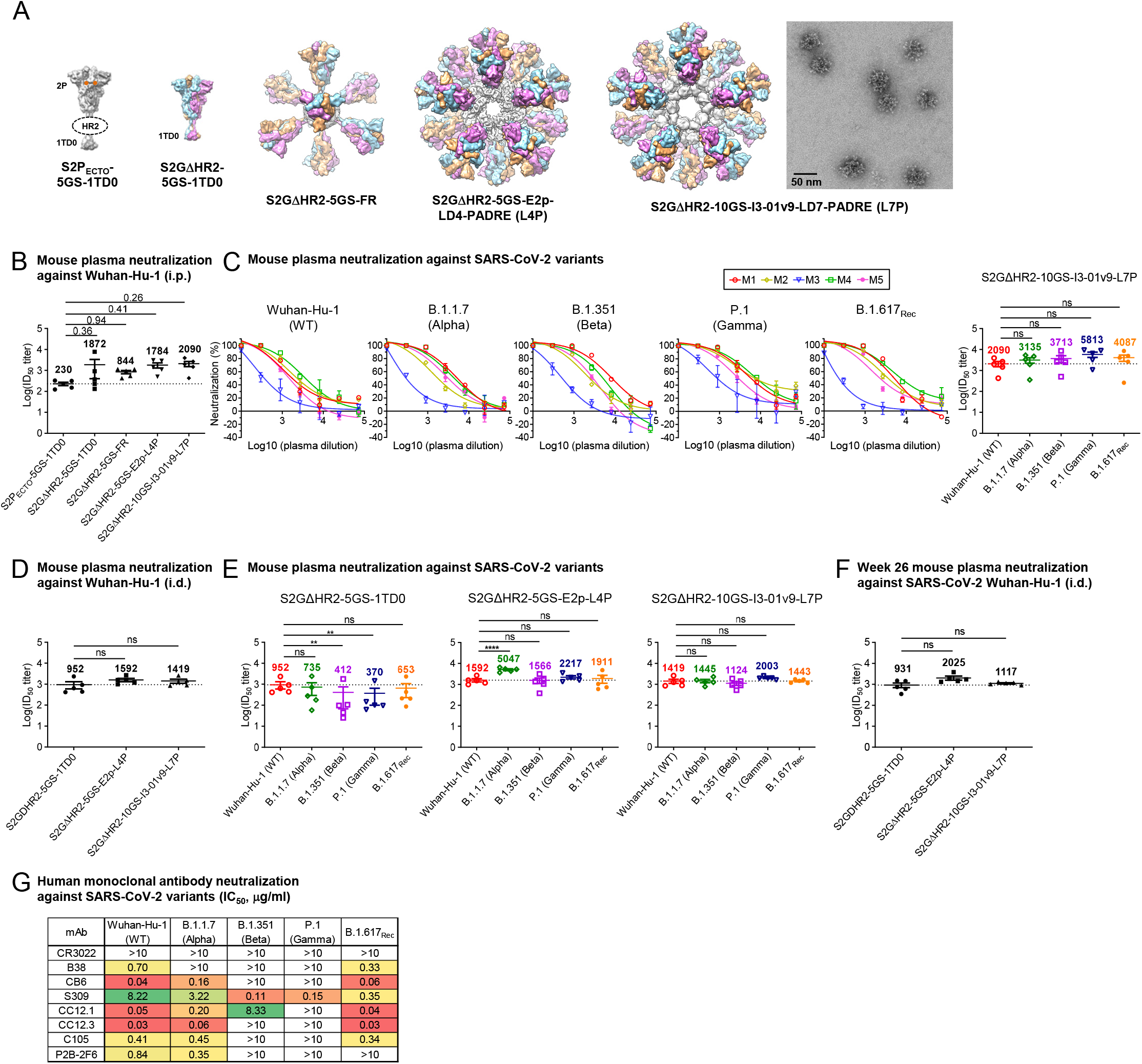
SApNP vaccines induce broadly neutralizing plasma responses to four representative SARS-CoV-2 variants. (**A**) Molecular surface representations of two spike (S2P_ECTO_-5GS-1TD0 and S2GΔHR2-5GS-1TD0) and three spike-SApNP (S2GΔHR2-5GS-ferritin [FR], S2GΔHR2-5GS-E2p-LD4-PADRE [E2p-L4P], and S2GΔHR2-10GS-I3-01v9-LD7-PADRE [I3-01v9-L7P]) vaccines. Representative EM image of S2GΔHR2-10GS-I3-01v9-L7P SApNPs is shown on the right. (**B**) Neutralization of the wildtype Wuhan-Hu-1 strain by mouse plasma induced by five different vaccines at week 5 after two intraperitoneal (i.p.) injections (*n* = 5 mice/group). ID_50_ titers derived from SARS-CoV-2-pp neutralization assays are plotted, with average ID_50_ values labeled on the plots. (**C**) Mouse plasma neutralization against Wuhan-Hu-1 and the B.1.1.7, B.1.351, P.1, and B.1.617_Rec_ variants at week 5 after two i.p. injections of the adjuvanted S2GΔHR2-10GS-I3-01v9-L7P vaccine (Left panels 1-5: percent neutralization plots; Right panel: ID_50_ plot). In (B) and (C), the plasma samples were generated in our previous study (*41*), in which mice were immunized with 50 μg of adjuvanted vaccine antigen. (**D**) Neutralization of mouse plasma against the wildtype Wuhan-Hu-1 strain induced by the S2GΔHR2 spike and two large SApNPs at week 5. Vaccines were administered via intradermal (i.d.) footpad injections (0.8 μg/injection, for a total of 3.3 μg/mouse). **(E)** Mouse plasma neutralization against Wuhan-Hu-1 strain and the B.1.1.7, B.1.351, P.1, and B.1.617_Rec_ variants at week 5 after two i.d. footpad injections. (**F**) Neutralization of mouse plasma against Wuhan-Hu-1 induced by the S2GΔHR2 spike and two large SApNPs at week 26. In (B)-(F), the ID_50_ values are plotted as mean ± SEM. The data were analyzed using one-way ANOVA for comparison between different vaccine groups or repeated measures ANOVA for comparison of ID_50_ titers from the same plasma sample against different SARS-Cov-2 strains. Dunnett’s multiple comparison *post hoc* test was performed. ns (not significant), \*\**p* < 0.01, \*\*\*\**p* < 0.0001. (**G**) Neutralization of five SARS-CoV-2 strains by eight human monoclonal antibodies. The IC_50_ values were calculated with the % neutralization range constrained within 0.0-100.0% and color-coded (white: IC_50_ > 10 μg/ml; green to red: low to high).

We first assessed the neutralizing activity of polyclonal plasma induced by various spike and SApNP vaccine formulations from our previous study (*41*) against the wildtype SARS-CoV-2 strain, Wuhan-Hu-1, as a baseline for comparison (**Fig. 1B**). Mouse plasma collected at week 5 after two intraperitoneal (i.p.) injections of adjuvanted vaccine antigens (50 μg) was analyzed in pseudoparticle (pp) neutralization assays (*43*). The soluble S2P_ECTO_ spike elicited the lowest 50% inhibitory dilution (ID_50_) titers, whereas the soluble S2GΔHR2 spike increased neutralization with a 7.1-fold higher average ID_50_ titer, which did not reach statistical significance because of within-group variation. All three spike-presenting SApNPs elicited superior neutralizing responses than the soluble S2P_ECTO_ spike (*41*). Notably, the I3-01v9 SApNP achieved the highest potency, with an average ID_50_ titer of 2090, which was 8.1-fold higher than the soluble S2P_ECTO_ spike. Despite differences in ID_50_ titers, the overall pattern remained the same as reported in our previous study (*41*). The differences might be attributable to the inherent variation of pseudovirus assays (*43, 44*). We then assessed plasma neutralization against four major SARS-CoV-2 variants (**Fig. 1C, fig. S1A, B**). The I3-01v9 SApNP induced a stronger neutralizing response against variants, with 0.5-fold (B.1.1.7), 0.8-fold (B.1.351), 1.8-fold (P.1), and 1.0-fold (B.1.617_Rec_) higher (or 1.5-2.8 times) ID_50_ titers compared with the wildtype strain (**Fig. 1C**). Altogether, these results confirmed our hypothesis and highlighted the advantages of spike-presenting SApNPs.

Next, we examined the influence of injection dosage and route on the plasma neutralizing response to various SARS-CoV-2 strains. To this end, we performed a mouse study, in which three groups of mice were immunized with 5, 15, and 45 μg of the I3-01v9 SApNP three times via i.p. injection. Remarkably, all four variants were neutralized by mouse plasma with comparable ID_50_ titers observed across dose groups (**fig. S1C, D**). To examine whether routes of injection affect the plasma neutralizing response against variants, we performed another mouse study, in which a low dose (3.3 μg) of adjuvanted antigen was intradermally administered into four footpads (i.e., 0.8 μg/footpad). At week 5, the large (∼55-60 nm) E2p and I3-01v9 SApNPs that present 20 S2GΔHR2 spikes yielded higher ID_50_ titers against the wildtype strain than the soluble S2GΔHR2 spike (**Fig. 1D, fig. S1E, F**), whereas a notable reduction of ID_50_ titers against the variants was noted for mouse plasma from the S2GΔHR2 group (**Fig. 1E, fig. S1E, F**), suggesting that multivalent display is critical for eliciting a broad neutralizing response. Overall, the E2p and I3-01v9 SApNP groups exhibited similar or slightly stronger plasma neutralization against the four variants relative to the wildtype strain, Wuhan-Hu-1 (**Fig. 1E**). Lastly, we assessed longevity of the low-dose vaccination-induced neutralizing response by testing week-26 plasma against Wuhan-Hu-1 (**Fig. 1F, fig. S1G, H**). It is noteworthy that ID_50_ titers at week 26 were at the same level as week 5, suggesting a long-lasting protective humoral immunity. In our previous study, a panel of human NAbs was used to evaluate antigenicity of the stabilized S2GΔHR2 spike and SApNPs and validate the SARS-CoV-2-pp neutralization assays (*41*). Here, this antibody panel was tested against SARS-CoV-2-pps that carry spikes of the wildtype strain and the four variants (**Fig. 1G, fig. S1I**). Lower potency against the B.1.351 and P.1 variants, measured by the 50% inhibitory concentration (IC_50_), was observed for all human NAbs, with the exception of NAb S309, which was identified from a SARS-CoV-1 patient (*45*). This finding is consistent with recent reports on convalescent patient plasma (*26-28*). Interestingly, most human NAbs remained effective against B.1.617_Rec_ showing a similar pattern to the wildtype Wuhan-Hu-1 strain and B.1.1.7 variant, consistent with the results of a recent cohort analysis of convalescent sera from individuals infected with early VOCs against a rising B.1.617 (*33*). As a negative control, mouse plasma induced by the S2GΔHR2-presenting I3-01v9 SApNP was tested against pseudoviruses carrying the murine leukemia virus (MLV) envelope glycoprotein (Env), or MLV-pps. Nonspecific MLV-pp neutralization was not detected for plasma samples produced in two independent immunization experiments (**fig. S1J, K**).

Altogether, our results demonstrate that spike-presenting SApNPs are more advantageous than soluble spikes in eliciting a strong neutralizing response to diverse SARS-CoV-2 variants. In our previous study, soluble SARS-CoV-2 spikes induced a more effective neutralizing response to SARS-CoV-1 than a scaffolded SARS-CoV-2 RBD trimer (*41*). Recently, a two-component RBD-NP vaccine showed reduced serum neutralization of variants bearing the E484K mutation (*46*). It is plausible that both the nanoparticle (NP) platform (one-component SApNP *vs*. two-component NP) and the antigen type (spike *vs*. RBD) contribute to vaccine breadth.

### Adjuvant effect on vaccine-induced neutralizing antibody and T-cell responses

Innate immunity plays an important role in regulating adaptive immunity, including humoral and cellular immune responses (*47-49*). Adjuvant-formulated vaccines have been shown to recruit and activate innate immune cells more effectively at injection sites and local lymph nodes (*50-52*). Among commonly used adjuvants, AH and AP create depots for the recruitment and activation of antigen-presenting cells (APCs) at injection sites and sentinel lymph nodes (*53, 54*), whereas oil-in-water emulsions such as MF59 promote antigen retention and APC stimulation in lymph nodes (*55*). Pattern recognition receptor (PRR) agonists (e.g., STING, TLR3, TLR4, TLR7/8, and TLR9 agonists) stimulate APCs at injection sites and nearby lymph nodes (*47, 52, 56-59*). Macrophage inhibitors (e.g., clodronate liposomes, termed CL) directly stimulate B cells or inhibit antigen sequestration by subcapsular sinus macrophages, thus resulting in more effective GC simulation in lymph nodes (*60*). Adjuvant combinations may generate a synergistic immune response by simultaneously activating multiple pathways (*52, 57*).

To examine the effect of innate signaling pathways on SApNP-induced immune responses, we tested 16 adjuvant formulations in a systematic study (**Fig. 2A**), in which mice were immunized with the adjuvanted I3-01v9 SApNP (20 μg) via intradermal injections in four footpads (i.e., 5 μg/footpad). We first tested mouse plasma neutralization against the wildtype Wuhan-Hu-1 stain. Mouse plasma at week 2 after a single dose was analyzed in SARS-CoV-2-pp assays, with most groups showing negligible or borderline ID_50_ titers (**fig. S2A, B**). We then analyzed mouse plasma at week 5 after two injections (**Fig. 2B, fig. S2C, D)**. The non-adjuvanted group showed detectable neutralization after two doses, with an average ID_50_ titer of 160, which was used as a baseline in this analysis. By comparison, conventional adjuvants, such as AH, AP, and AddaVax, increased ID_50_ titers by 10.3-, 7.6-, and 12.5-fold, respectively. The macrophage inhibitor CL boosted plasma neutralization by merely 1.6-fold relative to the non-adjuvanted group. Adjuvants that target various PRRs exhibited differential effects on plasma neutralization, increasing ID_50_ titers by 1.1- to 34.2-fold. Notably, STING and CpG (TLR9) substantially enhanced neutralizing titers, whereas TLR3, TLR4, and TLR7/8 agonists only exerted a modest effect. In most cases, adjuvants combined with AP further boosted plasma neutralizing activity. For example, when TLR4 and TLR7/8 agonists were mixed with AP, a 3.1-fold increase in ID_50_ titers was observed, suggesting a synergistic effect of stimulating multiple immune pathways. Overall, STING and CpG, either alone or combined with AP, showed plasma neutralization superior to that of any other adjuvant or adjuvant mix, increasing ID_50_ titers by 21 to 34-fold compared with the non-adjuvanted group. This is consistent with the results of the S-Trimer (SCB-2019), which, when formulated with CpG 1018 (TLR9 agonist) and alum adjuvants, induced potent NAb responses in nonhuman primates and human trials (*61, 62*). Mouse plasma at week 8 showed further increases in ID_50_ titers (1 to 3-fold) for most adjuvant groups (**Fig. 2C, fig. S2E, F**). Lastly, we examined mouse plasma at week 5 from the STING and CpG groups against the B.1.1.7, B.1.351, P.1, and B.1.617_rec_ variants (**Fig. 2D, fig. S2G, H**). Both adjuvant groups exhibited potent neutralizing responses to the four variants, with ID_50_ titers comparable to the wildtype strain.

**Fig. 2.**
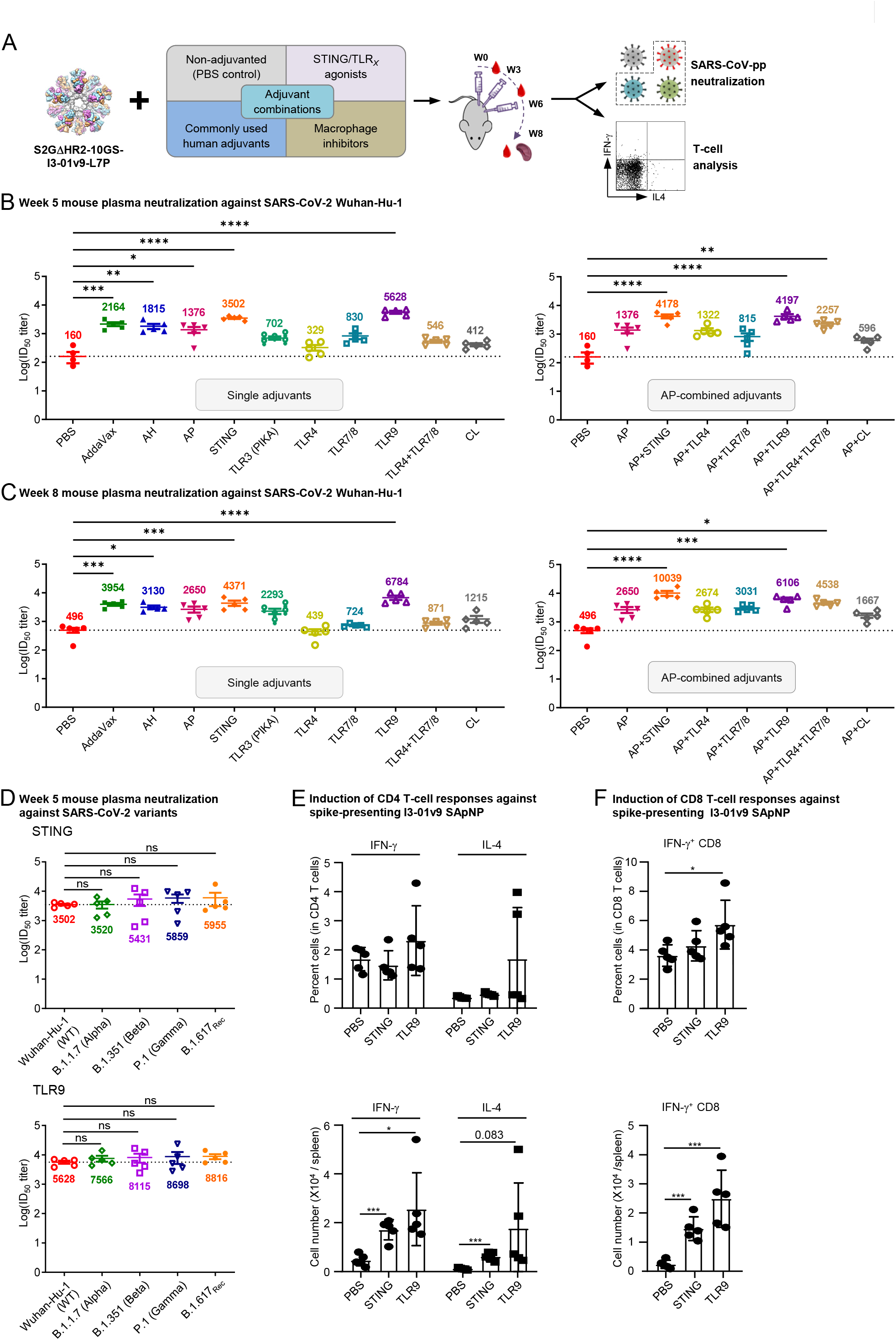
Adjuvants enhance the I3-01v9 SApNP vaccine -induced plasma neuralization of both the wildtype strain and four variants. **(A)** Schematic representation of mouse immunization with the I3-01v9 SApNP with diverse adjuvant formulations and functional assessment by SARS-CoV-2-pp neutralization assays and T-cell analysis. Conventional adjuvants, STING/TLR agonists, macrophage inhibitors, and adjuvant combinations were compared to non-adjuvanted control (PBS). (**B, C**) Mouse plasma neutralization against the wildtype SARS-CoV-2 strain, Wuhan-Hu-1, at weeks 5 and 8 after two and three footpad injections, respectively. ID_50_ titers derived from SARS-CoV-2-pp neutralization assays are plotted, with average ID_50_ values labeled on the plots. (**D**) Neutralization against four variants by mouse plasma from STING (top) and CpG (bottom)-formulated vaccine groups. ID_50_ titers derived from SARS-CoV-2-pp neutralization assays are plotted. Neutralization data were analyzed using either one-way ANOVA (B-C) or repeated measures one-way ANOVA (D) to compare ID_50_ titers. Dunnett’s multiple comparison *post hoc* test was performed. Splenic mononuclear cells derived from mice in the STING and CpG groups (*n* = 5 mice/group) at week 8 were cultured in the presence of BALB/C DCs pulsed with I3-01v9 SApNP (1 × 10^−7^ mM). Cells were harvested 16 h following re-activation. (**E**) Production of IFN-γ-producing Th1 CD4^+^ T cells and IL-4-producing Th2 CD4^+^ T cells. (**F**) IFN-γ-producing CD8^+^ effector T cells. T-cell responses were analyzed using one-way ANOVA followed by Tukey’s multiple comparison *post hoc* test. ns (not significant), \**p* < 0.05, \*\**p* < 0.01, \*\*\**p* < 0.001, \*\*\*\**p* < 0.0001.

We previously demonstrated that the AP-formulated I3-01v9 SApNP induces interferon-γ (IFN-γ)-producing CD4^+^ T helper 1 (Th1) cells and IFN-γ/interleukin-4 (IL-4) double-positive memory CD4^+^ T cells (*41*). Given the superior plasma neutralizing response observed for STING and CpG, we examined the impact of these two adjuvants on vaccine-induced T-cell responses. IFN-γ-producing CD4^+^ Th cells are important for optimal antibody responses and the induction of cellular immunity to clear viruses (*63-65*). To assess the effect of STING and CpG on vaccine-induced Th cells, we isolated splenocytes from mice 8 weeks after vaccination and cultured them in the presence of BALB/c mouse DCs pulsed with the spike-presenting I3-01v9 SApNP. Compared with the non-adjuvanted control, STING and CpG (TLR9) induced 3.7 and 5.5-fold more IFN-γ-producing CD4^+^ Th1 cells, and 5.5 and 16-fold more IL-4-producing CD4^+^ Th2 cells, respectively (**Fig. 2E, fig. S2I**). A visible but nonsignificant trend toward a higher frequency of both Th1 and Th2 cells was noted in mice immunized with the CpG-formulated I3- 01v9 SApNP than other formulations. Nonetheless, both adjuvants induced more IFN-γ-producing CD4^+^ Th1 cells than IL-4-producing CD4^+^ Th2 cells, suggesting a dominant Th1 response in these mice. This is consistent with the results for the S-Trimer (SCB-2019), which, when formulated with the AS03 adjuvant or mixed CpG 1018/alum adjuvants, induced Th1-biased cellular responses in mice (*61*). STING and CpG also enhanced CD8^+^ T-cell responses by 6- and 10-fold, respectively, compared with the PBS control. Notably, this effect was more visible for CpG in terms of both the frequency and number of IFN-γ-producing CD8^+^ effector T cells (**Fig. 2F, fig. S2J**).

Our results demonstrate that the I3-01v9 SApNP itself is immunogenic, and adjuvants can further enhance vaccine-induced NAb responses in plasma by up to 35-fold. The I3-01v9 SApNP, when formulated with the STING or TLR9 agonist, yielded the highest ID_50_ titers with robust CD4^+^ and CD8^+^ T-cell responses, highlighting their potential as adjuvants in the development of more effective SARS-CoV-2 vaccines.

### Diverse variant-neutralizing mouse antibody lineages identified by single-cell analysis

Although plasma neutralization confirmed the effectiveness of our newly designed SARS-CoV-2 vaccines (*41*) against variants, the nature of this response was unclear. It might result from multiple NAb lineages that each target a specific strain (non-overlapping), a few bNAb lineages that are each able to block multiple strains (overlapping), or a combination of both. Previously, we used antigen-specific single-cell sorting to identify potent mouse NAbs elicited by an I3-01 SApNP that presents 20 stabilized HIV-1 Env trimers (*66*). Here, we applied a similar strategy to decipher NAb responses induced by SARS-CoV-2 vaccines using mouse samples from our previous study (*41*), for which potent plasma neutralization against four variants has been verified (**Fig. 1C**).

Spleen samples from M4 in the spike group (S2GΔHR2-5GS-1TD0) and M2 in the spike-SApNP group (S2GΔHR2-10GS-I3-01v9-L7P), along with a control sample from M2 in the RBD (RBD-5GS-1TD0) group were analyzed. Two probes, RBD-5GS-foldon-Avi and S2GΔHR2-5GS-foldon-Avi, were produced, biotinylated, and purified to facilitate antigen-specific B-cell sorting (**fig. S3A, B**). Following antibody cloning, reconstituted mouse mAbs were tested for neutralizing activity against the wildtype strain, Wuhan-Hu-1, in SARS-CoV-2-pp assays. A total of 20 mAbs, four from the RBD group (**fig. S3C**), six from the spike group (**fig. S3D**), and 10 from the I3-01v9 SApNP group (**fig. S3E**), were found to be NAbs. The genetic analysis of mAb sequences revealed some salient features of the vaccine-induced NAb response in mice (**Fig. 3A**). Overall, these mAbs evolved from diverse germline origins. The RBD-elicited mAbs appeared to use distinct germline variable (V) genes for both heavy chain (HC) and κ-light chain (KC), or V_H_ and V_K_, respectively, whereas the spike and I3-01v9 SApNP-elicited mAbs shared some common V_H_ genes, such as IGHV14-1/3 and IGHV1S81. This result was not unexpected because the RBD vaccine presents a structurally distinct antigen to the immune system compared with the spike and I3-01v9 vaccines, which both present the S2GΔHR2 spike. These mAbs showed low levels of somatic hypermutation (SHM) with respect to their germline genes. Heavy chain complementarity-determining region (HCDR3) loops ranged from 4 to 12 aa in length, whereas most KCs contained 9 aa KCDR3 loops. Collectively, diverse germline genes and HCDR3 loops, accompanied by low degrees of SHM, suggest that many antibody lineages must have been generated upon vaccination, and some could achieve neutralizing activity without an extensive maturation process.

**Fig. 3.**
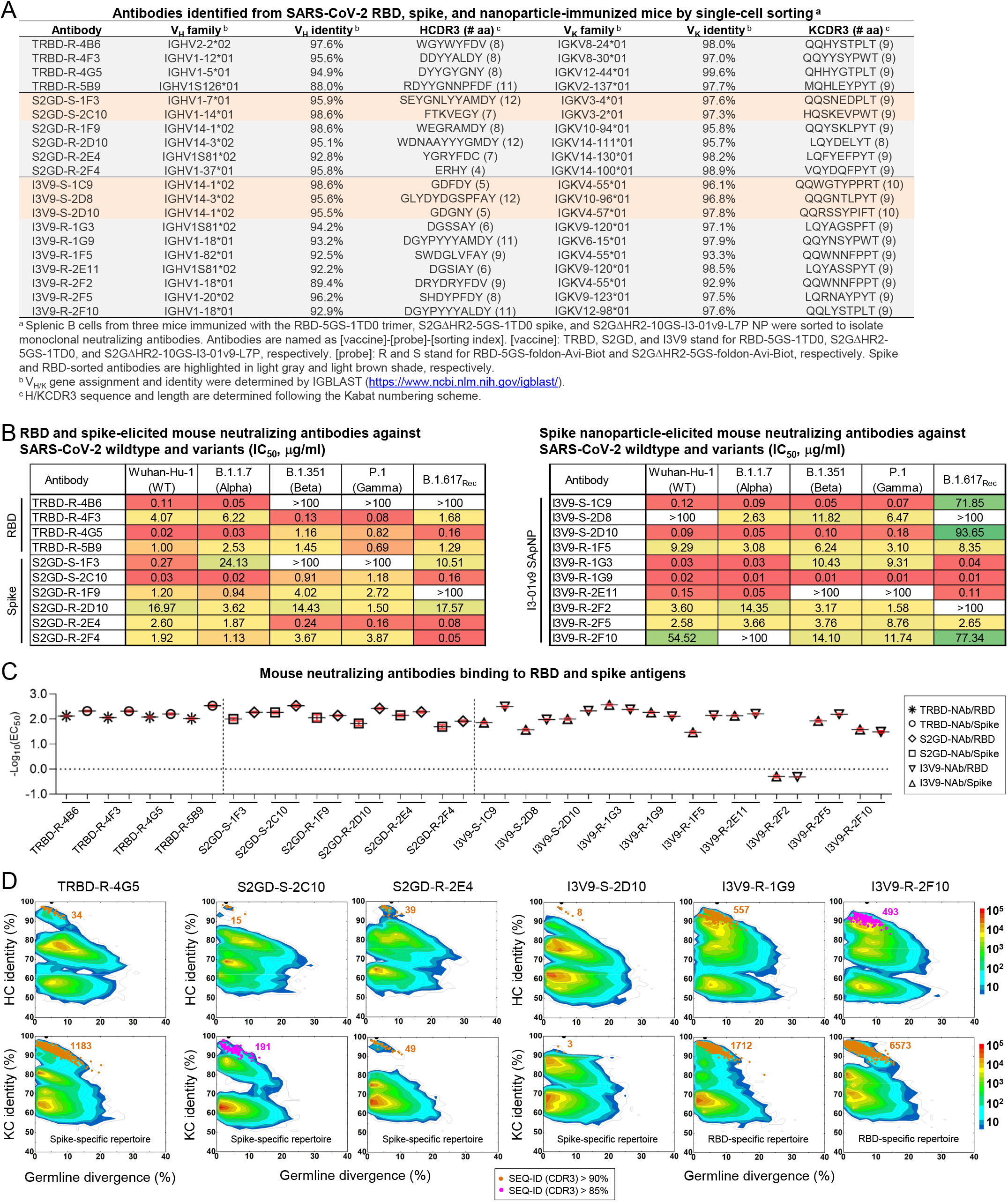
Single-cell isolation identifies vaccine-elicited mouse neutralizing antibody lineages with diverse breadth and potency. (**A**) Genetic analysis of 20 mouse antibodies identified from M2 in the RBD-5GS-1TD0 trimer group (four), M4 in the S2GΔHR2-5GS-1TD0 spike group (six), and M2 in the S2GΔHR2-10GS-I3-01v9-L7P SApNP group (ten). Antibodies isolated by the RBD and spike probes are highlighted in light gray and orange shade, respectively. (**B**) Neutralization of five SARS-CoV-2 strains by 10 RBD and spike-elicited mouse antibodies (left) and 10 SApNP-elicited mouse antibodies (right). The IC_50_ values were calculated with the %neutralization range constrained within 0.0-100.0% and color-coded (white: IC_50_ > 100 μg/ml; green to red: low to high). (**C**) EC_50_ (μg/ml) values of 20 mouse antibodies binding to the two SARS-CoV-2 antigens, the RBD monomer and S2GΔHR2-5GS-1TD0 spike, both with the Wuhuan-Hu-1 backbone. Antigen binding was measured by ELISA in duplicate, with mean value and standard deviation (SD) shown as black and red lines, respectively. (**D**) Divergence-identity analysis of selected mouse NAbs in the context of RBD/spike-specific splenic B cells. HCs and KCs are plotted as a function of sequence identity to the template and sequence divergence from putative germline genes. Color coding denotes sequence density. The template and sequences identified based on the *V* gene assignment and a CDR3 identity of 90%/85% or greater to the template are shown as black and orange/magenta dots on the 2D plots, with the number of related sequences labeled accordingly. The 2D plots for other NAbs are shown in **fig. S4D-F**.

We then examined the biological function of these mouse mAbs. Neutralizing activity was assessed in SARS-CoV-2-pp assays against the wildtype strain and four variants (**Fig. 3B, fig. S3F**). Overall, diverse yet consistent patterns were observed for the three sets of mAbs. Both the RBD vaccine (an RBD scaffold) and the two spike vaccines, albeit in different forms, appeared to elicit potent NAbs against the wildtype strain. MAbs TRBD-R-4G5, S2GD-S-2C10, and I3V9-R-1G9 showed similar IC_50_ values (0.02-0.03 μg/ml) against Wuhan-Hu-1, on par with the human NAbs CB6 (*67*) and CC12.1/3 (*68*) (**Fig. 1F**). All three vaccines elicited bNAb responses, despite variation in potency for different mAbs against different strains. Notably, I3V9-R-1G9, which was isolated from an I3-01v9 SApNP-immunized mouse, demonstrated high potency across all four variants (IC_50_: 0.01-0.02 μg/ml). This bNAb provided evidence that individual bNAb lineages may critically contribute to the plasma neutralization of diverse variants (**Fig. 1C**). All three vaccines generated NAbs that preferentially neutralize specific SARS-CoV-2 strains. For example, TRBD-R-4B6 was more effective against the wildtype strain and an early VOC, B.1.1.7, whereas S2GD-R-2E4 neutralized B.1.351, P.1, and B.1.617_Rec_ with greater potency. Notably, more than 60% (13) of the mAbs exhibited different patterns in the neutralization of B.1.617_Rec_ *vs*. VOCs B.1.351 and P.1, as indicated by the fold change in IC_50_, suggesting that B.1.617 may represent a distinct SARS-CoV-2 lineage. Although RBD-isolated NAbs likely neutralized SARS-CoV-2 by blocking its receptor binding, those spike-isolated NAbs could target the RBD, N-terminal domain (NTD), or epitopes in the S2 subunit. Thus, we tested these mAbs in an enzyme-linked immunosorbent assay (ELISA) against the RBD monomer and S2GΔHR2-5GS-1TD0 spike, both based on the wildtype Wuhan-Hu-1 backbone (**Fig. 3C, fig. S3G, H**). Overall, all of the NAbs bound the RBD and spike with a half-maximal concentration (EC_50_) of 0.034 μg/ml or lower, except for I3V9-R-2F2 (1.973 μg/ml for the RBD and 2.051 μg/ml for the spike). Most (15) NAbs showed greater binding affinity (or lower EC_50_ values) for the spike, suggesting that the two arms of the immunoglobulin (Ig) can each interact with one RBD of the spike, resulting in an avidity effect. Notably, diverse binding patterns were observed for I3-01v9 SApNP-elicited NAbs. Although I3V9-S-1C9 and I3V9-S-1F5 bound to the spike more favorably than the RBD, as indicated by a 4.7 to 4.9-fold reduction of their EC_50_ values, three NAbs from this group (I3V9-R-1G3, I3V9-R-1G9, and I3V9-R-2F10) preferred the RBD monomer over the spike. This preference might be explained by steric hindrance when these NAbs approach the RBDs on a trimeric spike at specific angles.

Lastly, we characterized these mouse NAbs in antigen-specific B-cell repertoires by next-generation sequencing (NGS), as previously demonstrated for NAbs isolated from HIV-1 SApNP-immunized mice and rabbits (*66*). Using the same RBD and spike probes (**fig. S3A**), ∼1500 splenic B cells were bulk-sorted from each of the three mice that were analyzed by single-cell sorting for mAb isolation (**fig. S4A**). Unbiased mouse antibody HC and KC libraries were constructed and sequenced on an Ion S5 platform, which yielded up to 4 million raw reads (**fig. S4B**). The antibody NGS data were then processed using a mouse antibodyomics pipeline (*69*) to remove low-quality reads, resulting in 0.11-0.41 full-length HCs and KCs (**fig. S4B**). Quantitative profiles of critical antibody properties, such as germline gene usage, the degree of SHM, and CDR3 loop length, were determined for the RBD and spike-specific B-cell populations (**fig. S4C**). All 20 single-cell-sorted mouse NAbs could well fall in the range of these repertoire profiles, but some V_H_/V_K_ genes that accounted for large portions of antigen-specific B cells, such as IGHV9 and IGHV5, were not used by any NAbs, suggesting that they might give rise to non-neutralizing binding antibodies. Two-dimensional (2D) divergence/identity plots were generated to visualize these NAbs in the context of NGS-derived B-cell repertoires (**Fig. 3D, fig. S4D-F**). Somatic variants were identified for each NAb by searching for sequences of the same V_H_/V_K_ gene with a CDR3 identity cutoff of 90% (or 85% for evolutionarily more remote variants). For the most potent NAb, TRBD-R-4G5, from an RBD-immunized mouse (M2), 34 HC variants were identified that overlapped with an “island” of high sequence similarity to TRBD-R-4G5 on the plot, whereas more KC variants (1183) were found, likely due to the lack of diversity in the KCDR3 region. A similar pattern was observed for the potent bNAb, I3V9-R-1G9, from an SApNP-immunized mouse (M2). By comparison, fewer putative somatic variants were identified for other NAbs in the antigen-specific B-cell repertoires regardless of the sorting probe used (**fig. S4D-F**), suggesting that these NAbs either were from less prevalent lineages or were generated in response to a previous injection (each mouse received four doses) (*41*). Similar observations were reported for the vaccination of nonhuman primates and humans in longitudinal repertoire analyses of single-cell-sorted NAbs (*70, 71*).

Single-cell isolation identified a panel of mouse mAbs with different neutralization breadth and potency against the wildtype SARS-CoV-2 strain and four major variants. The ELISA analysis suggested that the I3-01v9 SApNP can elicit NAbs with more diverse angles of approach to their epitopes than the RBD and soluble spike vaccine. Structural analysis by crystallography and EM will provide a more detailed understanding of epitope recognition by these mouse mAbs.

### Distribution and trafficking of I3-01v9 SApNP in mouse lymph node

After validating these vaccines against variants at both the plasma and mAb levels, we studied *in vivo* behaviors of the S2GΔHR2 spike and two large 60-meric SApNPs to understand why SApNPs outperform soluble spikes in bNAb elicitation. In principle, these SApNPs need to be transported to lymph nodes, retained, and presented to various immune cell populations to induce robust innate and adaptive immune responses. Here, we first examined the transport and distribution of I3-01v9 SApNPs in mouse lymph nodes via footpad injections (10 μg/footpad). The mice were sacrificed 12 h after single-dose (**Fig. 4A**) and prime-boost (**Fig. 4B**) regimens. The axillary, brachial, and popliteal sentinel lymph nodes were isolated for histological analysis. The lymph node tissues were stained with the human anti-spike antibody P2B-2F6 (*72*) to characterize SARS-CoV-2 spikes presented on the I3-01v9 SApNPs. Consistent with our previous study (*73*), SApNPs accumulated in lymph node follicles, regardless of the number of doses. SApNPs were sequestrated in the center of lymph node follicles after a single dose (**Fig. 4A**, images on the left, schematics on the right) but were located along the outer layer of expanded lymph node follicles after the second injection due to preexisting humoral immunity (i.e., GC reactions) that was induced by the first dose (**Fig. 4B**, images on the left, schematics on the right). Overall, the majority of SApNPs accumulated in lymph node follicles, but their distribution differed slightly, depending on the doses.

**Fig. 4.**
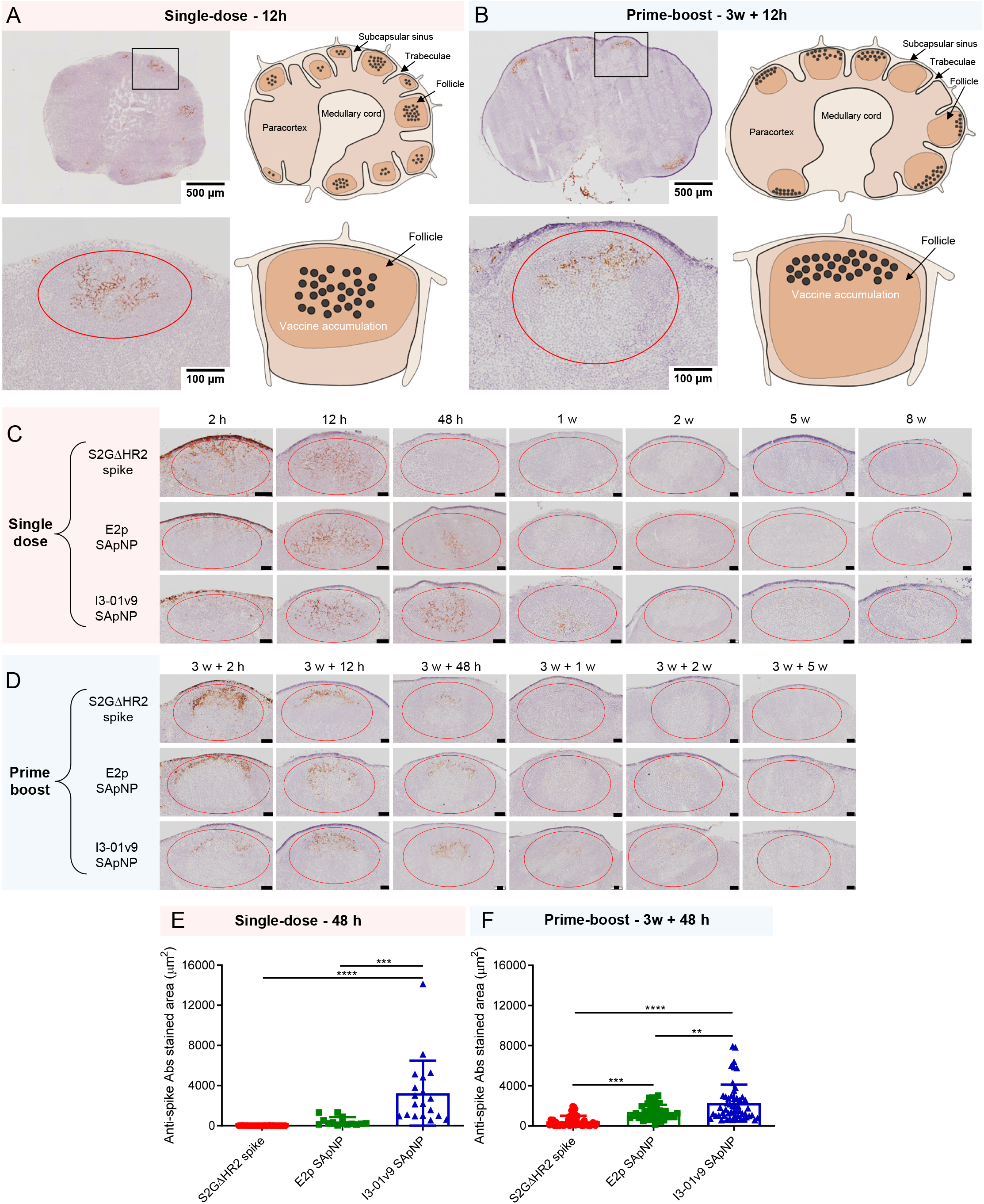
SARS-CoV-2 SApNP vaccines induce long-term lymph node follicle retention. (**A, B**) S2GΔHR2-presenting I3-01v9 SApNP vaccine distribution in a lymph node 12 h after (**A**) a single-dose or (**B**) prime-boost footpad injections (10 μg/footpad, 40 μg/mouse). A schematic illustration of SApNPs in lymph node follicles is shown. (**C, D**) Histological images of the S2GΔHR2 spike and S2GΔHR2-presenting E2p and I3-01 SApNP vaccine trafficking and retention in lymph node follicles 2 h to 8 weeks after (**C**) single-dose or (**D**) prime-boost injections, with a scale bar of 50 μm shown for each image. (**E, F**) Quantification of vaccine accumulation in lymph node follicles 48 h after (**E**) a single-dose or (**F**) prime-boost injections. Data were collected from more than 10 lymph node follicles (*n* = 3-4 mice/group). The data points are expressed as mean ± SD. The data were analyzed using one-way ANOVA followed by Tukey’s multiple comparison *post hoc* test. \*\**p* < 0.01, \*\*\**p* < 0.001, \*\*\*\**p* < 0.0001.

In this context, we examined patterns of trafficking and lymph node follicle retention for soluble S2GΔHR2 spike *vs*. the S2GΔHR2-presenting E2p and I3-01v9 SApNPs. To facilitate this analysis, the mice were sacrificed 2 h to 8 weeks after a single dose (**Fig. 4C**) and 2 h to 5 weeks after the boost (**Fig. 4D**). The injection dose was normalized to the total amount of protein (10 μg) per injection into each footpad (40 μg/mouse). As shown in **Fig. 4C**, the S2GΔHR2 spikes that trafficked into lymph node follicles at 2 h cleared within 48 h. In contrast, the two large SApNPs accumulated in the subcapsular sinus at 2 h and then trafficked into follicles 12 h after the single-dose injection. Remarkably, I3-01v9 SApNPs remained detectable in lymph node follicles after 2 weeks, suggesting 6-fold longer retention than the S2GΔHR2 spike (**Fig. 4C**). The results for these protein nanoparticles are thus consistent with the pattern of size dependency that was observed for ovalbumin-conjugated gold nanoparticles in our previous study (*73*), in which small (5-15 nm) nanoparticles cleared shortly after the injection, whereas large (50-100 nm) nanoparticles were retained in lymph node follicles for weeks. Similar patterns of antigen retention were observed after the second injection, although the boost appeared to exert a more positive effect on the soluble spike, which could be detected in lymph node follicles at 48 h (**Fig. 4D**). Nonetheless, prolonged retention was observed for both E2p and I3-01v9 SApNPs 2 weeks after the boost injection. Overall, the multivalent display of S2GΔHR2 spikes on the I3-01v9 SApNP resulted in 325- and 4-fold greater accumulation in lymph node follicles compared with the soluble spike 48 h after the single-dose **(Fig. 4E**) and prime-boost (**Fig. 4F**) injections, respectively. These findings reveal the advantage of a leading vaccine candidate identified in our previous study, S2GΔHR2-10GS-I3-01v9-L7P (*41*), in terms of spike antigen retention in lymph node follicles.

### Retention and presentation of I3-01v9 SApNP on follicular dendritic cell dendrites

Antigen retention and presentation in lymph node follicles are prerequisites to the stimulation of robust B cell responses and GC reactions (*34, 36*). Resident cells spatially rearrange antigens and present them to B cells. Follicular dendritic cells (FDCs) are resident stromal cells in follicles and retain soluble antigens, immune complexes, virus-like particles (VLPs), viruses, and bacteria (*73-76*). FDCs are also key to GC initiation, maintenance, and B-cell affinity maturation (*37, 77, 78*). Here, we hypothesized that FDCs comprise the major cell population in lymph node follicles that retain SARS-CoV-2 spikes and spike-presenting SApNPs. To test this hypothesis, we administered vaccines via footpad injections and collected mouse lymph nodes at the peak of accumulation (12 h) after single-dose (**Fig. 5A**) and prime-boost (**Fig. 5B**) injections. Lymph node tissue samples were stained with the anti-spike antibody P2B-2F6 (*72*) for the S2GΔHR2 spike, as well as anti-CD21 and anti-CD169 antibodies for FDCs and subcapsular sinus macrophages, respectively. The spike and SApNP (E2p or I3-01v9) signals colocalized with FDC (CD21^+^) networks in lymph node follicles (**Fig. 5A, B**). This result confirmed the critical role of FDC networks in mediating vaccine retention in lymph node follicles.

**Fig. 5.**
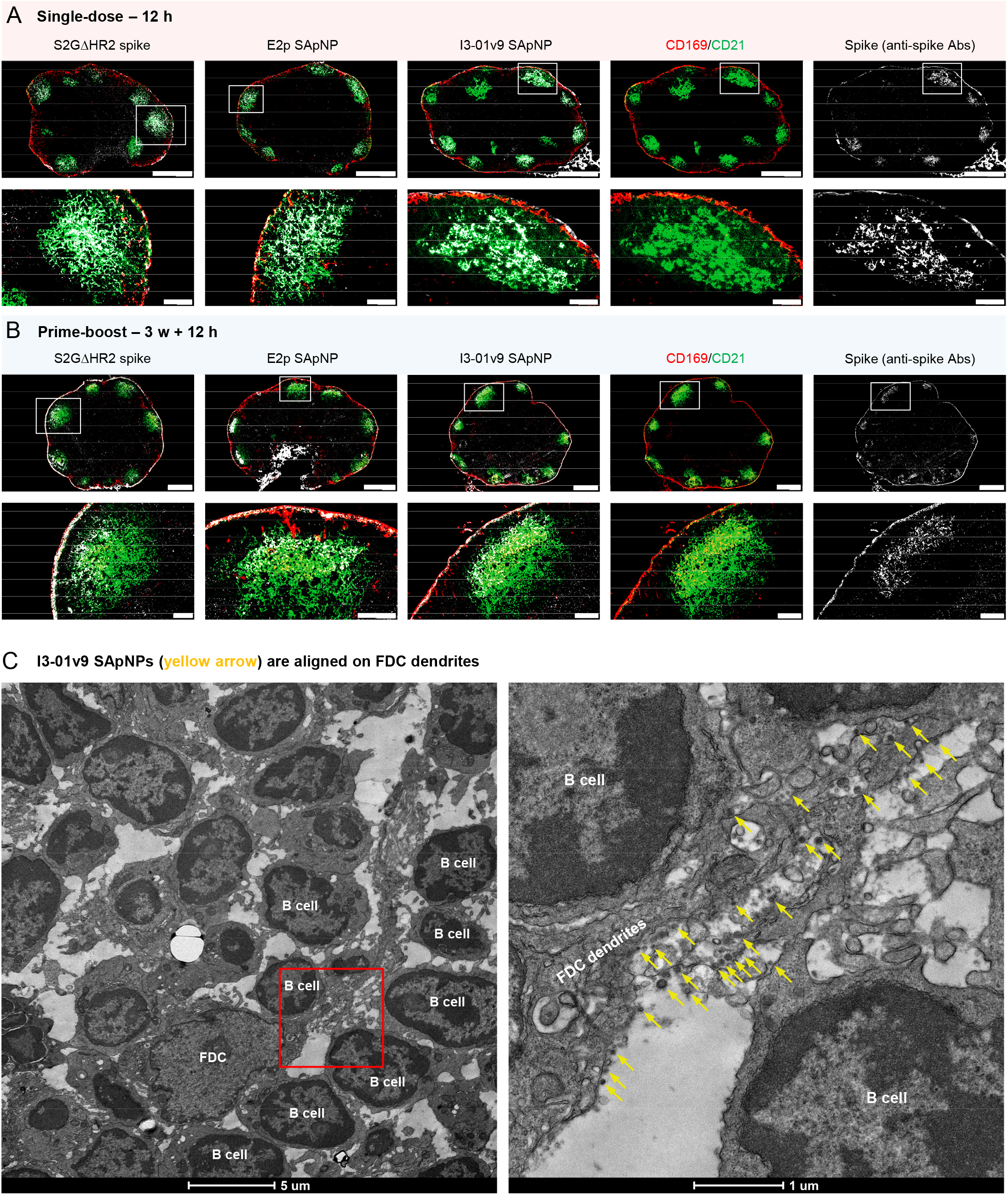
SARS-CoV-2 SApNP vaccines interact with follicular dendritic cells (FDCs) and are presented on FDC dendrites to B cells. (**A, B**) S2GΔHR2 spike and S2GΔHR2-presenting E2p and I3-01 SApNP vaccine interaction with FDC networks in lymph node follicles 12 h after (**A**) a single-dose or (**B**) prime-boost injections (10 μg/footpad, 40 μg/mouse). Vaccine antigens (the S2GΔHR2 spike and S2GΔHR2-presenting E2p and I3-01 SApNPs) colocalized with FDC networks. Immunostaining is color-coded (Green: CD21; Red: CD169; White: anti-spike), with scale bars of 500 μm and 100 μm shown for a complete lymph node and an enlarged image of a follicle, respectively. (**C**) Representative TEM images of an FDC surrounded by multiple B cells. S2GΔHR2-presenting I3-01 SApNPs (yellow arrows) presented on FDC dendrites.

The induction of potent bNAb responses by spike-presenting SApNPs in mice suggests the effective activation of naïve B cells and subsequent recalls by crosslinking B cell receptors (*76, 79, 80*). We visualized the interface between FDC networks and B cells to better understand how FDC networks present SApNPs to engage B cells. Briefly, fresh lymph nodes were isolated and directly immersed in fixative. The processed tissue samples were sectioned and stained on copper grids for TEM analysis. We first determined whether SApNPs, such as the S2GΔHR2-presenting I3-01v9 SApNP, remain intact *in vivo* (**fig. S5**). Mouse lymph nodes were isolated 2 h after the injection of a high dose (50 μg) of the non-adjuvanted I3-01v9 SApNP. The TEM images revealed that round-shape granules corresponding to intact SApNP aligned on the macrophage surface or inside endolysosomes of the macrophage in a lymph node (**fig. S5**). We next studied the relative location between FDCs and I3-01v9 SApNPs and how FDCs present SApNPs to B cells. Mouse lymph nodes were collected 2, 12, and 48 h after a single-dose (50 μg) and 12 h after the boost of the I3-01v9 SApNP vaccine. The FDCs exhibited the characteristic morphology of long dendrites that surrounded and interacted with B cells in lymph node follicles (**Fig. 5C, fig. S6**). Few I3-01v9 SApNPs were observed on FDC dendrites at 2 h (**fig. S6D**), whereas notably more nanoparticles migrated to and aligned on FDC dendrites at 12 and 48 h (**Fig. 5C, fig. S6A-C**, yellow arrows). The TEM images indicated that FDCs can present many SApNPs to neighboring B cells in this “hugging mode”, in which their long dendrites brace B cells to maximize interactions between multivalently displayed spikes and B cell receptors. These results demonstrated the intrinsic nature of FDCs as a reservoir for the sequestration, retention, and presentation of virus-like particles, or SApNPs with similar molecular traits, to initiate GC reactions.

### Robust germinal center reactions induced by spike-presenting SApNPs

Long-lived GC reactions induce immune stimulation for B-cell selection and affinity maturation, as well as production of immune memory and bNAb responses (*34, 35, 40*). Here, we investigated whether the prolonged retention of S2GΔHR2-presenting E2p and I3-01v9 SApNPs induces more robust GCs in lymph node follicles than the soluble S2GΔHR2 spike. Immunohistological analysis was performed to characterize GC B cells (GL7^+^) and T follicular helper (T_fh_) cells (CD4^+^Bcl6^+^). For the I3-01v9 SApNP, 2 weeks after immunization, we observed robust GCs in lymph node B cell follicles (B220^+^) with well-formed dark zone (DZ) and light zone (LZ) compartments, which contain GC B cells, FDCs, and T_fh_ cells (*35, 81-83*) (**Fig. 6A**). We then extended the analysis to the S2GΔHR2 spike and spike-presenting SApNPs 2, 5, and 8 weeks after the single-dose injection (**Fig. 6B, fig. S7A-C**) and 2 and 5 weeks after the boost (**Fig. 6C, fig. S7D, E**). Two metrics, the GC/FDC ratio (i.e., whether GC formation is associated with an FDC network, %) and GC size (i.e., occupied area), were used. Overall, the soluble spike and both large SApNPs induced robust GCs 2 weeks after immunization (**Fig. 6B, fig. S7A**). The E2p and I3-01v9 SApNPs that present 20 spikes induced robust, long-lived GCs, whereas the spike alone failed to sustain robust GCs at week 8 with either the single-dose (**Fig. 6B, D**) or prime-boost (**Fig. 6C, E**) injections. The I3-01v9 SApNP generated larger GCs than the soluble spike, 2.0-fold larger after the single dose (**Fig. 6B, D**) and 2.4-fold larger after the boost (**Fig. 6C, E**), measured at week 8.

**Fig. 6.**
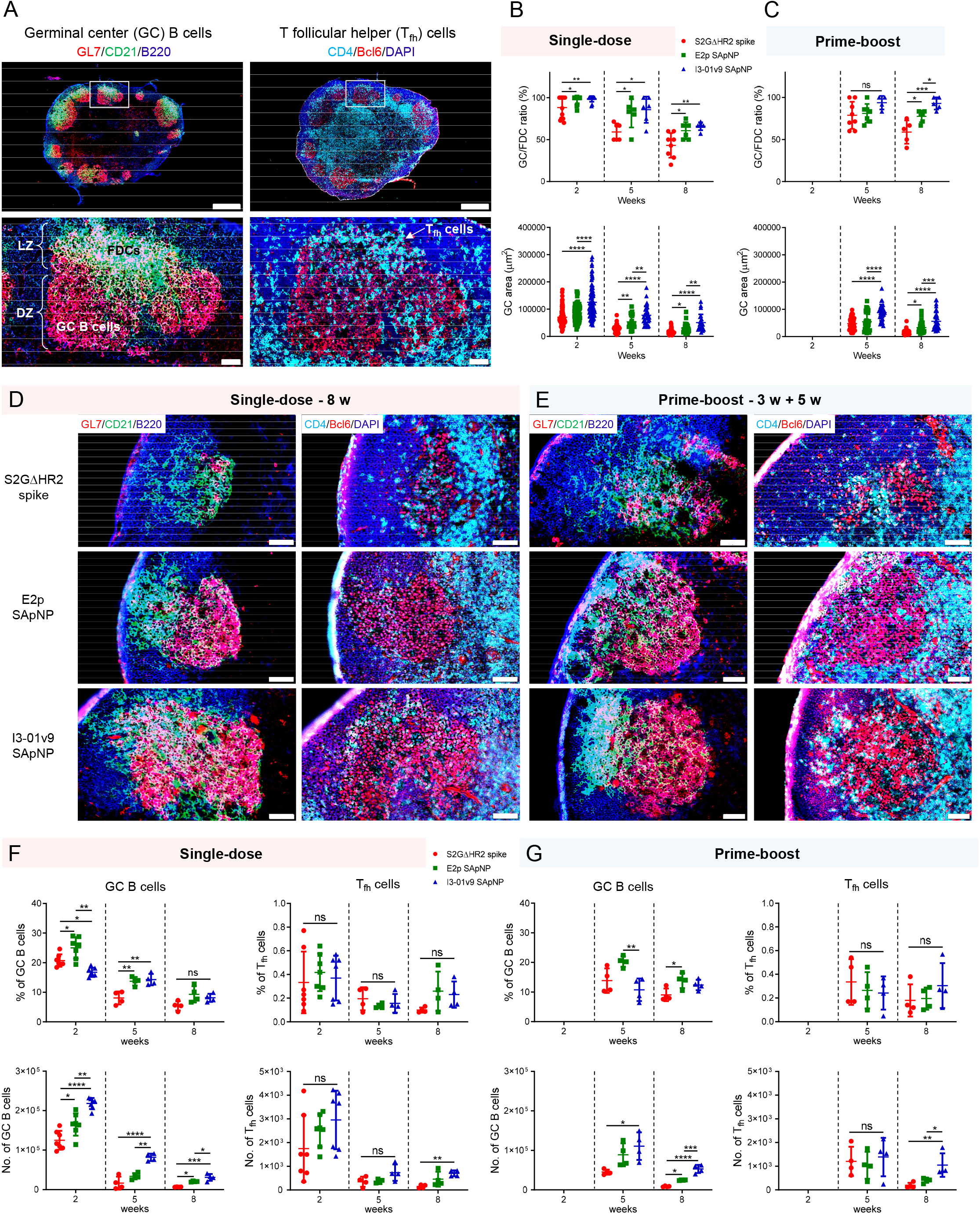
SARS-CoV-2 SApNP vaccines induce robust long-lived germinal centers. (**A**) Top: Representative immunohistological images of germinal centers at week 2 after a single-dose injection of the S2GΔHR2-presenting I3-01 SApNP vaccine (10 μg/injection, 40 μg/mouse). Bottom: Germinal center B cells (GL7^+^, red) adjacent to FDCs (CD21^+^, green) in lymph node follicles (left) and T_fh_ cells in the light zone (LZ) of germinal centers (right). Scale bars of 500 μm and 50 μm are shown for a complete lymph node and an enlarged image of a follicle, respectively. (**B, C**) Quantification of germinal center reactions using immunofluorescent images: GC/FDC ratio and sizes of germinal centers 2, 5, and 8 weeks after (**B**) single-dose or (**C**) prime-boost injections (*n* = 4-7 mice/group). The GC/FDC ratio is defined as whether the germinal center formation is associated with an FDC network (%). **(D, E)** Representative immunohistological images of germinal centers in mice immunized using S2GΔHR2 spike or S2GΔHR2-presenting E2p and I3-01 SApNP vaccines at week 8 after (**D**) single-dose or (**E**) prime-boost injections, with a scale bar of 50 μm shown for each image. (**F, G**) Quantification of germinal center reactions using flow cytometry: percentage and number of germinal center B cells and T_fh_ cells 2, 5, and 8 weeks after (**F**) single-dose or (**G**) prime-boost injections. The data points are shown as mean ± SD. The data were analyzed using one-way ANOVA followed by Tukey’s multiple comparison *post hoc* test for each timepoint. ns (not significant), \**p* < 0.05, \*\**p* < 0.01, \*\*\**p* < 0.001, \*\*\*\**p* < 0.0001.

We further characterized GC reactions by flow cytometry. Fresh mouse lymph nodes were disaggregated into a single cell suspension and stained with an antibody cocktail to quantify GC B cells and T_fh_ cells (**fig. S8A**). The results were consistent with the immunohistological analysis, in which all spike-based vaccine antigens, including the S2GΔHR2 spike and SApNPs, showed robust GCs at week 2 after the injection that declined over time, as measured at weeks 5 and 8 (**Fig. 6F**). The E2p and I3-01v9 SApNPs generated a larger population of GC B cells than both the S2P_ECTO_ and S2GΔHR2 spikes at week 2 (**fig. S8B, C**). Although the boost dose had little impact on the frequency of GC B cells and T_fh_ cells, it appeared to extend GC formation within lymph nodes (**Fig. 6F, G**), which may promote B cell development toward bNAbs. Notably, the GC B cell and T_fh_ cell populations elicited by the soluble S2GΔHR2 spike were barely detectable 5 weeks after immunization (**Fig. 6F, G**). This result was reminiscent of a recent study of an mRNA vaccine, in which GC reactions diminished to baseline levels at week 4 after a single-dose injection (*84*). The S2GΔHR2-presenting I3-01v9 SApNP generated 3.7/5.2-fold more GC B cells and 3.7/4.4-fold more T_fh_ cells than the soluble S2GΔHR2 spike at week 8 after one/two-dose immunization (**Fig. 6F, G**). Therefore, SApNPs that were retained on FDC dendrites could present NAb epitopes to enable more effective B cell recognition than the soluble spike, and consequently induce more robust and long-lived GCs in lymph nodes. Patterns of trafficking and retention may be specific to antigen size, as shown previously (*73*) and in the present study (**Figs. 4** and **5**), but GC reactions are largely determined by vaccine adjuvants. This effect was briefly demonstrated for the E2p and I3-01v9 SApNPs, which were previously formulated with the AddaVax and AP adjuvants (*41*). At week 2 after a single-dose injection, the adjuvanted SApNPs induced stronger GC reactions than the non-adjuvanted groups (**fig. S9**). This result can also explain the differences in plasma neutralization between the adjuvanted and non-adjuvanted I3-01v9 SApNPs **(Fig. 2)**.

NGS has been used to assess vaccine-draining lymph node B-cell responses (*85*). Here, we characterized lymph node B cells at the repertoire level for three groups of mice immunized with two doses (3.3 μg each) of the S2GΔHR2 spike, E2p, and I301v9 SApNPs via footpad injections. At this dosage, the spike showed less effective plasma neutralization of variants than the large SApNPs (**Fig. 1E**). Given their differences in retention, presentation, and GC reaction (**Figs. 4-6**), they were expected to yield different lymph node B-cell profiles. Interestingly, antigen-specific sorting identified more spike-targeting lymph node B cells from the I3-01v9 SApNP group than both the spike and E2p SApNP groups (**fig. S10A**). The antibody NGS data were processed by the mouse antibodyomics pipeline (*69*) (**fig. S10B**) to derive quantitative B-cell profiles (**fig. S10C-E**). Compared with the spike, the I3-01v9 SApNP appeared to activate fewer V_H_/V_K_ genes (**fig. S10F**, left two), while generating a larger population of spike-specific lymph node B cells (**fig. S10A**). The three vaccine groups exhibited a similar degree of SHM for V_H_ genes, with the I3-01v9 SApNP showing the highest SHM for V_K_ genes (**fig. S10F**, middle two). A highly uniform HCDR3 loop length distribution (∼10 aa) was observed for mice in the I3-01v9 SApNP group with little variation, as measured by the root-mean-square fluctuation (RMSF) (**fig. S10F**, right two). In our previous studies (*86, 87*), a similar approach was applied to assess hepatitis C virus (HCV) and Ebola virus (EBOV) vaccine-induced B cell responses in the spleen, a major lymphoid organ (*88*), after mice received four i.p. injections. We observed distinct B-cell profiles associated with the viral antigen and NP platform (*86, 87*). Here, the lymph node B-cell profiles appeared to be rather different, revealing the complex inner workings of another primary site for vaccine-induced immunity. Notably, I3-01v9 SANP exhibited more “focused” B-cell activation and development in vaccine-draining lymph nodes, as indicated by fewer activated germline genes and a narrower HCDR3 length distribution. More in-depth studies are needed to investigate the effect of injection route, adjuvant, and lymphoid organ, in addition to viral antigen and NP platform, on the resulting B-cell profiles. Single-cell immune profiling and antibody isolation (*89*) may provide further insights into the clonality of vaccine-induced B-cell lineages within lymph nodes.

## DISCUSSION

To end the COVID-19 pandemic, vaccines need to effectively block current and emerging SARS-CoV-2 variants that evade NAb responses by mutating key epitopes on the viral spike (*31*). To overcome this challenge, some suggested that COVID-19 vaccines need to be updated on a regular basis (*30-32*), whereas others developed mosaic or cocktail vaccines for related sarbecoviruses (*46, 90*). These vaccine strategies need to be evaluated for long-term protection, because SARS-CoV-2 is evolving rapidly and may acquire new mutations to evade vaccine-induced immunity (e.g., B.1.617) (*11*). In our previous study (*41*), the spike-presenting SApNPs induced a potent NAb response to SARS-CoV-1, which is evolutionarily much more distant to the wildtype SARS-CoV-2 strain, Wuhan-Hu-1, than all its circulating variants. Emerging data from human serum analysis suggested that vaccines derived from early pandemic strains may provide broad protection against current variants (*33*). Based on these findings, we hypothesized that SApNPs presenting stabilized ancestral Wuhan-Hu-1 spikes may provide an effective vaccine against SARS-CoV-2 variants. In the present study, we sought to confirm this hypothesis by testing four major variants and, if proven true, investigate the mechanism underlying such a broadly protective vaccine.

We explored several critical aspects related to the vaccine response, with a focus on the lead candidate identified in our previous study, S2GΔHR2-10GS-I3-01v9-L7P (*41*). We first tested vaccine-induced mouse plasma, which represents a polyclonal response, against four SARS-CoV-2 variants. Mouse plasma generated previously (*41*) and in new studies using different regimens (e.g., injection route, dosage, and adjuvant) potently neutralized the variants. Notably, SApNPs retained their high ID_50_ titers at a dosage as low as 3.3 μg, whereas formulations with the STING and TLR9 agonists further enhanced the I3-01v9 SApNP-induced neutralizing response. While plasma neutralization data may be interpreted with caution due to assay variation (*44*), single-cell-sorted mAbs provided unambiguous evidence of the vaccine-induced bNAb response. Our results revealed that a plethora of NAb lineages were generated upon vaccination, with I3-01v9 SApNP being the most effective at eliciting bNAbs. Additionally, our results confirmed the necessity of a prime-boost strategy for eliciting a potent NAb response, regardless of the regimen (e.g., injection route, dosage, and adjuvant). Such an NAb response, once generated, can persist for an extended period of time post vaccination. Although SARS-CoV-2 challenge in relevant animal models gives more accurate assessment of vaccine protection (*91*), NAb titers have been found to be highly predictive of immune protection from symptomatic infections in a large cohort study (*92*). Protein vaccines, despite the well-established records of safety and effectiveness, have yet to be deployed to mitigate the COVID-19 pandemic (*93-95*). One protein vaccine, NVX-CoV2373 (micelle-attached spikes formulated with the Matrix-M™ adjuvant), showed ∼90% efficacy in human trials (*19*). Our study indicates that SApNPs displaying 20 stabilized spikes provide a promising protein vaccine candidate that can be used either alone or as a booster for nucleic acid (e.g., mRNA and viral vector) vaccines in the battle against emerging SARS-CoV-2 variants (*11*).

We explored the mechanism of SApNP *vs*. spike vaccines following the previously used strategy to analyze the *in vivo* behaviors of antigen-attached gold nanoparticles (*73*). In principle, SApNP vaccines must induce long-lasting GCs to facilitate the development of bNAbs. Effective vaccine retention and presentation are critical for inducing and sustaining GC reactions, which in turn promote the proliferation and affinity maturation of antigen-specific B cells. Indeed, we found that the I3-01v9 SApNP, our leading vaccine candidate (*41*), elicited 6-fold longer retention and 4-fold greater accumulation in lymph node follicles than the stabilized S2GΔHR2 spike alone with a prime-boost regimen. This can be attributed to the intrinsic physiological properties of lymph nodes that mediate vaccine trafficking and retention in follicles in a size-dependent manner, which would favor retaining large (> 50 nm) virus-like particles (*73-75, 80, 96*). Supporting this notion are the TEM images of retained SApNPs aligned on long FDC dendrites, suggesting that such protein nanoparticles can present spike antigens to B cells for rapid initiation and then sustain GC reactions in lymph node follicles for an extended period of time. Specifically, the I3-01v9 SApNP generated 2.4-fold larger GCs and greater numbers of GC B cells (5.2-fold) and T_fh_ cells (4.4-fold) than the soluble S2GΔHR2 spike with the prime-boost regimen. These findings provide quantitative evidence that spike-presenting SApNPs are uniquely suited for inducing long-lived robust GCs in lymph node follicles. Our analyses thus shed light on the mechanism by which the I3-01v9 SApNP can elicit a more effective bNAb response than the soluble spike.

Rational design of next-generation COVID-19 vaccines requires an in-depth understanding of bNAb elicitation (*31*). Superior NAb (but not necessarily bNAb) responses have been reported for several vaccine candidates that employ particulate display (*90, 97-104*). The I3-01v9 SApNP elicited a potent bNAb response to four variants, overcoming a major challenge facing the current COVID-19 vaccines. Mechanistic studies of vaccine trafficking, retention, presentation, and GC reactions provided valuable insights into the spike and SApNP-induced immunity (*95, 105, 106*). Such knowledge, if can be obtained for other vaccine platforms (e.g., inactivated whole virions, mRNAs, and viral vectors) will facilitate rational selection of the most effective vaccine candidates to mitigate the pandemic and ultimately stop the spread of SARS-CoV-2.

## MATERIALS AND METHODS

### SARS-CoV-2 spike and SApNP vaccine antigens

The design, expression, and purification of a stabilized SARS-CoV-2 spike, S2GΔHR2, and three SApNPs that present either 8 or 20 S2GΔHR2 spikes were described in our recent study (*41*). Briefly, the spike gene of the SARS-CoV-2 isolate Wuhan-Hu-1 (GenBank accession no. MN908947) was modified to include the mutations ^682^GSAGSV^687^ and K986G/V987G, in addition to truncation of the HR2 stalk (ΔE1150-Q1208). The viral capsid protein SHP (Protein Data Bank: 1TD0) was added as a C-terminal trimerization motif to stabilize the S2GΔHR2 trimer, resulting in a soluble S2GΔHR2-5GS-1TD0 spike (*41*). The S2GΔHR2 spike was genetically fused to ferritin (FR), multilayered E2p, and multilayered I3-01v9 with 5GS, 5GS, and 10GS linkers, respectively, resulting in three S2GΔHR2-presenting SApNPs (*41*). An S2P_ECTO_-5GS-1TD0 spike construct that contained the mutations ^682^GSAGSV^687^ and K986G/V987G but without HR2 deletion (*41*) was included for comparison. All vaccine antigens were transiently expressed in ExpiCHO cells and purified by a CR3022 antibody column and size-exclusion chromatography (SEC) as described previously (*41*). Briefly, ExpiCHO cells were thawed and incubated with ExpiCHO™ Expression Medium (Thermo Fisher) in a shaker incubator at 37 °C at 135 rotations per minute (rpm) with 8% CO_2_. When the cells reached a density of 10×10^6^ ml^-1^, ExpiCHO™ Expression Medium was added to reduce cell density to 6×10^6^ ml^-1^ for transfection. The ExpiFectamine™ CHO/plasmid DNA complexes were prepared for 100-ml transfection in ExpiCHO cells according to the manufacturer’s instructions. For a given construct, 100 μg of plasmid and 320 μl of ExpiFectamine™ CHO reagent were mixed in 7.7 ml of cold OptiPRO™ medium (Thermo Fisher). After the first feed on day 1, ExpiCHO cells were cultured in a shaker incubator at 33 °C at 115 rpm with 8% CO_2_ according to the Max Titer protocol with an additional feed on day 5 (Thermo Fisher). Culture supernatants were harvested 13-14 days after transfection, clarified by centrifugation at 4000 rpm for 25 min, and filtered using a 0.45 μm filter (Thermo Fisher). The CR3022 antibody column was used to extract SARS-CoV-2 antigens from the supernatants, followed by SEC on a Superdex 200 10/300 GL column (for scaffolded RBD trimers), a Superose 6 16/600 GL column (for the S2GΔHR2 spike, with and without Avi-tag), or a Superose 6 10/300 GL column (for SApNPs). Protein concentration was determined using UV_280_ absorbance with theoretical extinction coefficients.

### Animal immunization and sample collection

Similar immunization protocols were reported in our previous vaccine studies (*41, 86, 87*). Briefly, Institutional Animal Care and Use Committee (IACUC) guidelines were followed for all of the animal studies. BALB/c mice (6 weeks old) were purchased from the Jackson Laboratory and kept in ventilated cages in environmentally controlled rooms at The Scripps Research Institute. The mouse studies were conducted according to Association for the Assessment and Accreditation of Laboratory Animal Care guidelines, and the protocols were approved by the IACUC. For the immunogenicity study, the mice were intraperitoneally immunized at weeks 0 and 3 with 200 μl of antigen/adjuvant mix containing 5-50 μg of vaccine antigen and 100 μl of adjuvant (*41*), or intradermally immunized at weeks 0 and 3 with 80 μl of antigen/adjuvant mix containing 3.3 μg of vaccine antigen and 40 μl of adjuvant. The intradermal (i.d.) immunization was done through injections into four footpads, each with 20 μl of antigen/adjuvant mix. For the mechanistic study of vaccine trafficking, retention, and induced GCs, the mice were immunized at weeks 0 and 3 with 80 μl of antigen/adjuvant mix containing 40 μg of vaccine antigen per mouse. To visualize the I3-01v9 SApNPs in lymph node tissues using TEM, each mouse was immunized at weeks 0 and 3 with 140 μl of antigen/adjuvant mix containing 100 μg of vaccine antigen (40 μl of adjuvant) into the two hind footpads. Vaccines were intradermally administered into mouse footpads using a 29-gauge insulin needle under 3% isoflurane anesthesia with oxygen. Blood was drawn from the maxillary/facial vein into an ethylenediaminetetraacetic acid (EDTA)-coated tube 2 weeks after each immunization. Plasma was isolated from blood after centrifugation at 14000 rpm for 10 min. Plasma was heat-inactivated at 56°C for 30 min, with the supernatant collected after centrifugation at 8000 rpm for 10 min. Plasma was used in pseudovirus neutralization assays to determine the vaccine-induced NAb responses. The axillary, brachial, and popliteal sentinel lymph nodes were collected at the end timepoint for further analysis.

### Experimental adjuvants and formulation

The adjuvants squalene-oil-in-water (AddaVax), aluminum hydroxide (AH), aluminum phosphate (AP), 2’3’-c-di-AM(PS)2 (Rp,Rp) (STING ligand), monophosphoryl lipid A from S. minnesota (MPLA-SM) R595 (TLR4 agonist), imidazoquinoline compound R848 (TLR7/8 agonist), and CpG ODN 1826, Class B (murine) (TLR9 agonist) were purchased from InvivoGen. PIKA, a TLR3 agonist with enhanced T cell and antibody responses reported for a Phase I rabies vaccine trial (*107*), was used as an adjuvant. PIKA was generously provided by Yisheng Biopharma and included in this study as an adjuvant that activates the TLR3 pathway. Macrophage inhibitors clodronate liposomes (Liposoma BV, catalog no. CP-005-005) were used to eliminate subcapsular sinus macrophages in lymph nodes to promote more robust B-cell activation. Mouse immunization was performed to examine the effects of 16 adjuvants or adjuvant combinations on the I3-01v9 SApNP-induced immune response with respect to the non-adjuvanted vaccine (PBS instead of an adjuvant). Vaccine antigen and adjuvants were mixed thoroughly 10 min before immunization. Each mouse was intradermally immunized at weeks 0, 3, and 6 with 120-140 μl of antigen/adjuvant mix containing 20 μg of vaccine antigen (I3-01v9 SApNP) and 80-100 μl of adjuvant, which was evenly split and injected into four footpads. Mouse blood was isolated at weeks 5 and 8 after two and three intradermal injections, respectively. Spleens and lymph nodes were harvested at week 8 for immunological analyses. Spleen samples were ground through a 70 μm cell strainer to release splenocytes into a cell suspension. Splenocytes were spun down at 400 × *g* for 10 min, washed with PBS, and treated with the ammonium-chloride-potassium (ACK) lysing buffer (Lonza). Splenocytes were then frozen with 3 ml of Bambanker freezing media.

### SARS-CoV-2 pseudovirus neutralization assay

The SARS-CoV-2-pp neutralization assays were described in our previous study (*41*). Briefly, SARS-CoV-2-pps were generated by the co-transfection of HEK293T cells with the HIV-1 pNL4-3.lucR-E-plasmid (obtained from the National Institutes of Health AIDS reagent program; https://www.aidsreagent.org/) and the expression plasmid encoding the *S* gene of five SARS-CoV-2 strains, including the wildtype Wuhan-Hu-1 strain (GenBank accession no. MN908947), three VOCs (GISAID accession no. EPI_ISL_601443, EPI_ISL_678597, and EPI_ISL_792680 for B.1.1.7, B.1.351, and P.1, respectively), and B.1.617_Rec_, a reconstituted strain based on an early analysis of the B.1.617 lineage (*11*). The HEK293T-hACE2 cell line (catalog no. NR-52511) and pcDNA3.1(-) vector containing the *S* gene of the wildtype Wuhan-Hu-1 strain (catalog no. NR52420) were requested from the BEI Resources (https://www.beiresources.org/) on September 23, 2020 and used in the pseudovirus neutralization assays (*43*). Based on sequence alignment, spike mutations were incorporated into the *S* gene of the Wuhan-Hu-1 strain (catalog no. NR52420) to create respective expression plasmids for B.1.1.7, B.1.351, P.1, and B.1.617_Rec_. For B.1.617_Rec_, G142D, L452R, E484Q, D614G and P681R were included as representative spike mutations in this SARS-CoV-2 lineage (*11*). SARS-CoV-2-pp neutralization by immunized mouse plasma and human or mouse mAbs was performed according to our previously described protocol (*41*). Using the same co-transfection expression system as described above for the SARS-CoV-2-pps, we produced pseudoviruses carrying the murine leukemia virus (MLV) Env, MLV-pps, for use as a negative control (*41*). Percent neutralization data were analyzed using GraphPad Prism 9.1.2 software. ID_50_/IC_50_ values were calculated using constraints for percent neutralization (0-100%), whereas unconstrained neutralization plots are shown in **Fig. 1** and **figs. S1-S3**.

### Enzyme-linked immunosorbent assay

Each well of a Costar™ 96-well assay plate (Corning) was first coated with 50 µl of PBS containing 0.2 μg of the appropriate antigens. The plates were incubated overnight at 4 °C, and then washed five times with wash buffer containing PBS and 0.05% (v/v) Tween 20. Each well was then coated with 150 µl of blocking buffer consisting of PBS and 40 mg/ml blotting-grade blocker (Bio-Rad). The plates were incubated with blocking buffer for 1 h at room temperature, and then washed five times with wash buffer. Mouse mAbs, in the immunoglobulin G (IgG) form, were diluted in blocking buffer to a maximum concentration of 10 μg/ml followed by a 10-fold dilution series. For each dilution, a total volume of 50 μl was added to the appropriate wells. Each plate was incubated for 1 h at room temperature and then washed five times with PBS containing 0.05% Tween 20. A 1:5000 dilution of horseradish peroxidase (HRP)-conjugated goat anti-human IgG antibody (Jackson ImmunoResearch Laboratories) was then made in wash buffer (PBS containing 0.05% Tween 20), with 50 μl of this diluted secondary antibody added to each well. The plates were incubated with the secondary antibody for 1 h at room temperature and then washed six times with PBS containing 0.05% Tween 20. Finally, the wells were developed with 50 μl of TMB (Life Sciences) for 3-5 min before stopping the reaction with 50 μl of 2 N sulfuric acid. The resulting plate readouts were measured at a wavelength of 450 nm. The ELISA data were analyzed to calculate EC_50_ values using GraphPad Prism 9.1.2 software.

### Histology, immunostaining, and imaging

The mice were sacrificed 2 h to 8 weeks after a single-dose immunization and 2 h to 5 weeks after the boost immunization. The axillary, brachial, and popliteal sentinel lymph nodes were isolated for histological analysis. Fresh lymph nodes were rapidly merged into frozen section compound (VWR International, catalog no. 95057-838) in a plastic cryomold (Tissue-Tek at VWR, catalog no. 4565) using liquid nitrogen to preserve antigens on the cell membrane and spike. Lymph node samples were stored at -80°C and sent to the Centre for Phenogenomics (http://phenogenomics.ca) on dry ice for sample processing and imaging. Tissue sections (8 μm) were cut on a cryostat (Cryostar NX70) and collected on charged slides. Sections were post-fixed in 10% neutral buffered formalin and permeabilized in PBS containing 0.5% Triton X-100 before immunostaining. Protein Block (Agilent) was used to block nonspecific antibody binding before incubating the sections with primary antibody overnight at 4°C. After washing in TBST, the sections were incubated in fluorophore-conjugated secondary antibodies for 1 h at room temperature. Lymph node tissue sections were stained with human anti-spike antibody P2B-2F6 (*72*) (1:50) and biotinylated goat anti-human secondary antibody (Abcam, catalog no. ab7152, 1:300), followed by streptavidin-HRP reagent (Vectastain Elite ABC-HRP Kit, Vector, catalog no. PK-6100) and diaminobenzidine (DAB) (ImmPACT DAB, Vector, catalog no. SK-4105) to study the distribution and retention of the soluble S2GΔHR2 spike alone and S2GΔHR2 spike-presenting E2p and I3-01v9 SApNPs. For immunofluorescent staining, tissue sections were stained for FDCs using anti-CD21 antibody (Abcam, catalog no. ab75985, 1:1800) followed by anti-rabbit secondary antibody conjugated with Alexa Fluor 555 (Thermo Fisher, catalog no. A21428, 1:200), stained for B cells using anti-B220 antibody (eBioscience, catalog no. 14-0452-82, 1:100) followed by anti-rat secondary antibody conjugated with Alexa Fluor 674 (Thermo Fisher, catalog no. A21247, 1:200), and stained for subcapsular sinus macrophages using anti-sialoadhesin (CD169) antibody (Abcam, catalog no. ab53443, 1:600) followed by anti-rat secondary antibody conjugated with Alexa Fluor 488 (Abcam, catalog no. ab150165, 1:200). Germinal center B cells were labeled using rat anti-GL7 antibody (FITC; BioLegend, catalog no. 144604, 1:250). T_fh_ cells were labeled using anti-CD4 antibody (BioLegend, catalog no. 100402, 1:100) followed by anti-rat secondary antibody conjugated with Alexa Fluor 488 (Abcam, catalog no. ab150165, 1:1000) and Bcl6 antibody (Abcam, catalog no. ab220092, 1:300) followed by anti-rabbit secondary antibody conjugated with Alexa Fluor 555 (Thermo Fisher, catalog no. A21428, 1:1000). Nuclei were then counterstained with 4′,6-diamidino-2-phenylindole (DAPI) (Sigma-Aldrich, catalog no. D9542, 100 ng/ml). The stained tissue sections were scanned using an Olympus VS-120 slide scanner and imaged using a Hamamatsu ORCA-R2 C10600 digital camera for all bright-field and fluorescent images. Bright-field images of stained S2GΔHR2 spike and S2GΔHR2 spike-presenting SApNPs in lymph node follicles and fluorescent images of GCs were quantified using ImageJ software (*108*).

### Electron microscopy analysis of protein nanoparticles and lymph node tissues

Electron microscopy (EM) analysis was performed by the Core Microscopy Facility at The Scripps Research Institute. For the negative-staining EM analysis of protein nanoparticles, the S2GΔHR2-10GS-I3-01v9-L7P SApNP samples were prepared at a concentration of 0.01 mg/ml. Carbon-coated copper grids (400 mesh) were glow-discharged, and 10 µl of each sample was adsorbed for 2 min. Excess sample was wicked away and grids were negatively stained with 2% uranyl formate for 2 min. Excess stain was wicked away and the grids were allowed to dry. For the EM analysis of mouse tissues, the lymph nodes were dissected from each animal and immersed in oxygenated 2.5% glutaraldehyde and 4% paraformaldehyde in 0.1M Na cacodylate buffer (pH 7.4) fixative overnight at 4°C. After washing in 0.1 M sodium cacodylate buffer, the tissue samples were post-fixed in buffered 1% osmium tetroxide and 1.5% potassium ferrocyanide for 1-1.5 h at 4°C, rinsed in the same buffer, and then stained *en bloc* with 0.5% uranyl acetate overnight at 4°C. The tissue samples were washed in double-distilled H_2_O and dehydrated through a graded series of ethanol followed by acetone, infiltrated with LX-112 (Ladd) epoxy resin, and polymerized at 60°C. Ultrathin lymph node sections (at 70-nm thickness) were prepared for imaging. Samples were analyzed at 80 kV with a Talos L120C transmission electron microscope (Thermo Fisher), and images were acquired with a CETA 16M CMOS camera.

### Lymph node disaggregation, cell staining, and flow cytometry

Germinal center reactions, including the percentage of GC B cells (GL7^+^B220^+^) and T_fh_ cells (CD3^+^CD4^+^CXCR5^+^PD-1^+^), and the number of GC B cells and T_fh_ cells were studied by flow cytometry (**fig. S5A**). The mice were sacrificed 2, 5, and 8 weeks after a single-dose immunization and 2 and 5 weeks after the boost immunization. Fresh axillary, brachial, and popliteal sentinel lymph nodes were collected and mechanically disaggregated. These lymph node samples were merged in enzyme digestion solution containing 958 μl of Hanks’ balanced salt solution (HBSS) buffer (Thermo Fisher Scientific, catalog no. 14185052), 40 μl of 10 mg/ml collagenase IV (Sigma-Aldrich, catalog no. C5138), and 2 μl of 10 mg/ml of DNase (Roche, catalog no. 10104159001) in an Eppendorf tube. After incubation at 37°C for 30 min, lymph node samples were filtered through a 70 μm cell strainer and spun down at 400 × *g* for 10 min. The supernatant was discarded, and the cell pellet was resuspended in HBSS blocking solution containing 0.5% (w/v) bovine serum albumin and 2 mM EDTA. The nonspecific binding of Fc receptors was blocked using anti-CD16/32 antibody (BioLegend, catalog no. 101302) on ice for 30 min. Cocktail antibodies, Zombie NIR live/dead stain (BioLegend, catalog no. 423106), Brilliant Violet 510 anti-mouse/human CD45R/B220 antibody (BioLegend, catalog no. 103247), FITC anti-mouse CD3 antibody (BioLegend, catalog no. 100204), Alexa Fluor 700 anti-mouse CD4 antibody (BioLegend, catalog no. 100536), PE anti-mouse/human GL7 antibody (BioLegend, catalog no. 144608), Brilliant Violet 605 anti-mouse CD95 (Fas) antibody (BioLegend, catalog no. 152612), Brilliant Violet 421 anti-mouse CD185 (CXCR5) antibody (BioLegend, catalog no. 145511), and PE/Cyanine7 anti-mouse CD279 (PD-1) antibody (BioLegend, catalog no. 135216) were then mixed with the cells and placed on ice for 30 min. After washing cells with HBSS blocking solution after antibody staining, the samples were fixed using 1.6% paraformaldehyde (Thermo Fisher Scientific, catalog no. 28906) in HBSS on ice for 30 min. The cell samples were stored in HBSS blocking solution for the flow cytometry study. Sample events were acquired by a 5-laser BD Biosciences LSR II analytical flow cytometer with BD FACS Diva 6 software at the Core Facility of The Scripps Research Institute. The data were further processed using FlowJo 10 software.

### DC production, T cell culture, activation, and flow cytometry analysis

Mouse bone marrow (BM) was cultured in RPMI 1640 medium containing 10% fetal bovine serum (FBS) and recombinant mouse Fms-like tyrosine kinase 3 ligand (Flt3L, 50 ng/ml) and stem cell factor (SCF, 10 ng/ml) for 9 days as previously described (*109*). To induce DC activation, immature DCs were incubated with lipopolysaccharide (LPS, 100 ng/ml) plus R848 (Resiquimod, 100 ng/ml) overnight, which activated TLR4 or TLR7/8 signaling, respectively. Cells were harvested for the experiments. CD11c^+^ DCs were sorted using magnetic beads (Miltenyi-Biotech, CA). Splenic mononuclear cells from each group of immunized mice were cultured in the presence of DCs pulsed with or without I3-01v9 SApNP (1 × 10^−7^ mM) in complete IMDM medium containing IL-2 (5.0 ng/ml). Cells were collected 16 h later for intracellular cytokine staining and flow cytometry. All antibodies used for immunofluorescence staining were purchased from eBioscience (San Diego, CA), BioLegend (San Diego, CA) or BD Biosciences (San Jose, CA). Magnetic microbead-conjugated streptavidin was purchased from Miltenyi-Biotech (Auburn, CA). Recombinant human IL-2 protein was purchased from R&D Systems (Minneapolis, MN). Recombinant mouse Flt3 ligand (Flt3L) and mouse SCF were purchased from Shenandoah Biotech (Warwick, PA). Cells were stained with appropriate concentrations of mAbs. Dead cells were excluded using Fixable Viability Dye (eBioscience, CA). Flow cytometry was performed using LSRII (BD Bioscience, CA).

### Bulk and single-cell sorting of SARS-CoV-2 antigen-specific mouse B cells

Spleens or lymph nodes were harvested from mice 15 days after the last immunization, and the cell suspension was prepared. Dead cells were excluded by staining with the Fixable Aqua Dead Cell Stain kit (Thermo Fisher, catalog no. L34957). FcγIII (CD16) and FcγII (CD32) receptors were blocked by adding 20 µl of 2.4G2 mAb (BD Pharmigen, catalog no. N553142). The cells were then incubated with 10 μg of a biotinylated RBD-5GS-foldon-Avi trimer or biotinylated S2GΔHR2-5GS-foldon-Avi spike. Briefly, the probes were generated by the biotinylation of Avi-tagged SARS-CoV-2 antigens using biotin ligase BirA according to the manufacturer’s instructions (Avidity). Biotin excess was removed by SEC on either a Superdex 200 10/300 column (GE Healthcare) for the RBD probe or a HiLoad Superose 6 16/600 column (GE Healthcare) for the spike probe. In the SEC profiles, the probe peak was well separated from the peak of biotin ligase (**fig. S3A**). Cells and biotinylated proteins were incubated for 5 min at 4 °C, followed by the addition of 2.5 µl of anti-mouse IgG fluorescently labeled with FITC (Jackson ImmunoResearch catalog no. 115-095-071) and incubated for 15 min at 4 °C. Finally, 5 µl of premium-grade allophycocyanin (APC)-labeled streptavidin was added to the cells and incubated for 15 min at 4 °C. In each step, the cells were washed with 0.5 ml of PBS and the sorting buffer (PBS with 2% FBS). FITC^+^ APC^+^ probe-specific B cells were sorted using MoFloAstrios EQ (Beckman Coulter). For bulk sorting, positive cells were sorted into an Eppendorf microtube with 20 μl of lysis buffer. For single B-cell sorting, individual positive cells were sorted into the inner wells of a 96-well plate with 20 μl of pre-reverse transcription (RT) lysis mix containing 0.1 μl of NP40 (Sigma-Aldrich), 0.5 μl of RNAse Inhibitor (Thermo Fisher), 5 μl of 5× First Strand Buffer, and 1.25 μl of DTT from the SuperScript IV kit (Invitrogen), with 13.15 μl of H_2_O per well.

### Antibody cloning from Env-specific single B cells and antibody production

The antibody cloning of SARS-CoV2-2 antigen-sorted single B cells was conducted as follows. A mix containing 3 μl of Random Hexamers (GeneLink), 2 μl of dNTPs, and 1 μl of SuperScript IV enzyme (Thermo Fisher) was added to each well of a single-cell-sorted 96-well plate that underwent thermocycling according to the program outlined in the SuperScript IV protocol, resulting in 25 μl of cDNA for each single cell. cDNA (5 μl) was then added to a polymerase chain reaction (PCR) mix containing 12.5 μl of 2× Multiplex PCR mix (Qiagen), 9 μl of H_2_O, 0.5 μl of forward primer mix, and 0.5 μl of reverse mouse primer mix (*110*) for heavy and κ-light chains within each well. A second PCR reaction was then performed using 5 μl of the first PCR as the template and respective mouse primers (*110*) according to the same recipe as the first PCR. The PCR products were run on 1% Agarose gel and those with correct heavy and light chain bands were then used for Gibson ligation (New England Biolabs), cloning into human IgG expression vectors, and transformation into competent cells. Mouse mAbs were expressed by the transient transfection of ExpiCHO cells (Thermo Fisher) with equal amounts of paired heavy and κ-light chain plasmids. Antibody proteins were purified from the culture supernatant after 12-14 days using Protein A bead columns (Thermo Fisher).

### NGS and bioinformatics analysis of mouse B cells

Previously, a 5′-rapid amplification of cDNA ends (RACE)-PCR protocol was developed for the deep sequencing analysis of mouse B-cell repertoires (*69*). In the present study, this protocol was applied to analyze bulk-sorted, RBD/spike-specific mouse B cells. Briefly, 5’
s-RACE cDNA was obtained from bulk-sorted B cells of each mouse with the SMART-Seq v4 Ultra Low Input RNA Kit for Sequencing (TaKaRa). The IgG PCRs were set up with Platinum *Taq* High-Fidelity DNA Polymerase (Life Technologies) in a total volume of 50 µl, with 5 μl of cDNA as the template, 1 μl of 5’-RACE primer, and 1 μl of 10 µM reverse primer. The 5′-RACE primer contained a PGM/S5 P1 adaptor, and the reverse primer contained a PGM/S5 A adaptor. The mouse 3’-C_γ_1-3/3’-C_μ_ inner primers and 3’-mC_κ_ outer primer (*110*) were adapted as reverse primers for the 5’-RACE PCR processing of heavy and κ-light chains. A total of 25 cycles of PCR was performed and the expected PCR products (500-600 bp) were gel purified (Qiagen). NGS was performed on the Ion S5 GeneStudio system. Briefly, heavy and κ-light chain libraries from the same mouse were quantitated using a Qubit 2.0 Fluorometer with the Qubit dsDNA HS Assay Kit and then mixed at a 3:1 ratio before being pooled with antibody libraries of other mice at an equal ratio for sequencing. Template preparation and (Ion 530) chip loading were performed on Ion Chef using the Ion 520/530 Ext Kit, followed by sequencing on the Ion S5 system with default settings (*86*). The mouse Antibodyomics pipeline (*69*) was used to process raw NGS data, derive quantitative profiles for germline gene frequency, the degree of SHM, and CDR3 loop length distribution, and generate 2D divergence/identity plots to visualize mAbs in their respective repertoires (*86*).

### Statistical analysis

Data were collected from 4-7 mice per group. All of the statistical analyses were performed and graphs were generated using GraphPad Prism 9.1.2 software. In the analysis of vaccine-induced plasma neutralization, different vaccine groups were compared using one-way analysis of variance (ANOVA), whereas for a given vaccine group ID_50_ titers of the same plasma sample against different variants were compared using repeated measures one-way ANOVA. In both cases, they were followed by Dunnett’s multiple comparison *post hoc* test. For the vaccine accumulation and GC study, different vaccine groups were compared using one-way ANOVA, followed by Tukey’s multiple comparison *post hoc* test. Statistical significance was indicated as the following: ns (not significant), \**p* < 0.05, \*\**p* < 0.01, \*\*\**p* < 0.001, \*\*\*\**p* < 0.0001.

## SUPPLEMENTARY MATERIALS

Supplementary material for this article is available at http://xxx/xxx/xxx.

**fig. S1**. Spike and spike-presenting SApNP vaccine-induced neutralizing antibody responses against the wildtype SARS-CoV-2 strain and four variants.

**fig. S2**. Immune responses against the wildtype SARS-CoV-2 strain and four variants induced by the I3-01v9 SApNP formulated with different adjuvants.

**fig. S3**. Single-cell isolation and functional evaluation of monoclonal neutralizing antibodies from mice immunized with the RBD, spike, and SApNP vaccines.

**fig. S4**. Unbiased repertoire analysis of bulk-sorted SARS-CoV-2 antigen-specific mouse splenic B cells and tracing of mouse neutralizing antibodies in the NGS-derived repertoires.

**fig. S5**. SARS-CoV-2 spike-presenting SApNP interaction with macrophages in a lymph node.

**fig. S6**. TEM images of SARS-CoV-2 spike-presenting I3-01v9 SApNP interaction with FDCs in a lymph node.

**fig. S7**. Immunohistological analysis of SARS-CoV-2 spike/spike-presenting SApNP vaccine-induced GCs.

**fig. S8**. Flow cytometry analysis of SARS-CoV-2 spike/spike-presenting SApNP vaccine-induced GCs.

**fig. S9**. Adjuvant effect on SARS-CoV-2 spike/spike-presenting SApNP vaccine-induced GCs.

**fig. S10**. NGS analysis of SARS-CoV-2 spike-specific lymph node (LN) B cells from mice immunized with the spike and SApNP vaccines.

## Author contributions

Project design by Y.-N.Z., L.H., and J.Z. SARS-CoV-2 variant plasmid design and processing by L.H. and C.S. Antigen production, purification, and basic characterization by T.N., T.F., and L.H. Antibody and mouse plasma neutralization by J.P., T.F., and L.H. Mouse immunization, plasma collection, and lymph node isolation by Y.-N.Z. Vaccine-induced T-cell response analysis by Y.W., C.A., and Y.Z. B cell sorting, antibody cloning, and synthesis by C.S. and L.H. Antibody expression, purification, and ELISA by C.S. and T.N. NGS and bioinformatics by L.H. and J.Z. Immunohistology, TEM, and flow cytometry by Y.-N.Z. Manuscript written by Y.-N.Z., Y.Z., L.H., and J.Z. All authors commented on the manuscript. This is manuscript number 30082 from The Scripps Research Institute.

## Data and material availability

All data are available in the main text or in the supplementary materials. Additional data related to this paper may be requested from the corresponding author.

## References

1. J. M. Dan et al., Immunological memory to SARS-CoV-2 assessed for up to 8 months after infection. Science 371, eabf4063 (2021).

2. B. Isho et al., Persistence of serum and saliva antibody responses to SARS-CoV-2 spike antigens in COVID-19 patients. Sci. Immunol. 5, eabe5511 (2020).

3. A. S. Iyer et al., Persistence and decay of human antibody responses to the receptor binding domain of SARS-CoV-2 spike protein in COVID-19 patients. Sci. Immunol. 5, eabe0367 (2020).

4. Y. Chen et al., Quick COVID-19 healers sustain anti-SARS-CoV-2 antibody production. Cell 183, 1496-1507.e1416 (2020).

5. R. Burioni, E. J. Topol, Assessing the human immune response to SARS-CoV-2 variants. Nat. Med., Published Ahead-of-Print (2021).

6. H. Tegally et al., Emergence of a SARS-CoV-2 variant of concern with mutations in spike glycoprotein. Nature, Published Ahead-of-Print (2021).

7. R. Challen et al., Risk of mortality in patients infected with SARS-CoV-2 variant of concern 202012/1: matched cohort study. BMJ 372, n579 (2021).

8. N. G. Davies et al., Estimated transmissibility and impact of SARS-CoV-2 lineage B.1.1.7 in England. Science, eabg3055 (2021).

9. E. C. Sabino et al., Resurgence of COVID-19 in Manaus, Brazil, despite high seroprevalence. Lancet 397, 452–455 (2021).

10. J. R. Mascola, B. S. Graham, A. S. Fauci, SARS-CoV-2 viral variants—tackling a moving target. JAMA, Published Ahead-of-Print (2021).

11. S. Cherian et al., Convergent evolution of SARS-CoV-2 spike mutations, L452R, E484Q and P681R, in the second wave of COVID-19 in Maharashtra, India. bioRxiv, 2021.2004.2022.440932 (2021).

12. E. Callaway, Delta coronavirus variant: scientists brace for impact. Nature, Published Ahead-of-Print (2021).

13. C. Zimmer, J. Corum, W. S.-L., in Coronavirus vaccine tracker. (https://www.nytimes.com/interactive/2020/science/coronavirus-vaccine-tracker.html).

14. L. R. Baden et al., Efficacy and safety of the mRNA-1273 SARS-CoV-2 vaccine. N. Engl. J. Med. 384, 403–416 (2020).

15. T. C. Williams, W. A. Burgers, SARS-CoV-2 evolution and vaccines: cause for concern? Lancet Respir. Med., Published Ahead-of-Print (2021).

16. D. Y. Logunov et al., Safety and efficacy of an rAd26 and rAd5 vector-based heterologous prime-boost COVID-19 vaccine: an interim analysis of a randomised controlled phase 3 trial in Russia. Lancet 397, 671–681 (2021).

17. F. P. Polack et al., Safety and efficacy of the BNT162b2 mRNA Covid-19 vaccine. N. Engl. J. Med. 383, 2603–2615 (2020).

18. M. G. Thompson et al., Prevention and attenuation of Covid-19 with the BNT162b2 and mRNA-1273 vaccines. N. Engl. J. Med., Published Ahead-of-Print (2021).

19. P. T. Heath et al., Safety and efficacy of NVX-CoV2373 Covid-19 vaccine. N. Engl. J. Med., Published Ahead-of-Print (2021).

20. Q. Li et al., The impact of mutations in SARS-CoV-2 spike on viral infectivity and antigenicity. Cell 182, 1284-1294.e1289 (2020).

21. C. Rees-Spear et al., The impact of spike mutations on SARS-CoV-2 neutralization. bioRxiv, 2021.2001.2015.426849 (2021).

22. C. K. Wibmer et al., SARS-CoV-2 501Y.V2 escapes neutralization by South African COVID-19 donor plasma. bioRxiv, 2021.2001.2018.427166 (2021).

23. E. Andreano et al., SARS-CoV-2 escape in vitro from a highly neutralizing COVID-19 convalescent plasma. bioRxiv, 2020.2012.2028.424451 (2020).

24. M. Hoffmann et al., SARS-CoV-2 variants B.1.351 and P.1 escape from neutralizing antibodies. Cell, Published Ahead-of-Print (2021).

25. W. F. Garcia-Beltran et al., Multiple SARS-CoV-2 variants escape neutralization by vaccine-induced humoral immunity. Cell, Published Ahead-of-Print (2021).

26. P. Wang et al., Antibody resistance of SARS-CoV-2 variants B.1.351 and B.1.1.7. bioRxiv, 2021.2001.2025.428137 (2021).

27. P. Supasa et al., Reduced neutralization of SARS-CoV-2 B.1.1.7 variant by convalescent and vaccine sera. Cell, Published Ahead-of-Print (2021).

28. D. A. Collier et al., SARS-CoV-2 B.1.1.7 sensitivity to mRNA vaccine-elicited, convalescent and monoclonal antibodies. medRxiv, 2021.2001.2019.21249840 (2021).

29. D. Planas et al., Sensitivity of infectious SARS-CoV-2 B.1.1.7 and B.1.351 variants to neutralizing antibodies. Nat. Med. 27, 917–924 (2021).

30. Z. Wang et al., mRNA vaccine-elicited antibodies to SARS-CoV-2 and circulating variants. Nature, Published Ahead-of-Print (2021).

31. D. R. Burton, E. J. Topol, Variant-proof vaccines - invest now for the next pandemic. Nature 590, 386–388 (2021).

32. Q. Li et al., No higher infectivity but immune escape of SARS-CoV-2 501Y.V2 variants. Cell, Published Ahead-of-Print (2021).

33. C. Liu et al., Reduced neutralization of SARS-CoV-2 B.1.617 by vaccine and convalescent serum. Cell, (2021).

34. A. Singh, Eliciting B cell immunity against infectious diseases using nanovaccines. Nat. Nanotechnol. 16, 16–24 (2021).

35. G. D. Victora, M. C. Nussenzweig, Germinal centers. Annu. Rev. Immunol. 30, 429–457 (2012).

36. R. Rappuoli, Glycoconjugate vaccines: principles and mechanisms. Sci. Transl. Med. 10, eaat4615 (2018).

37. M. Akkaya, K. Kwak, S. K. Pierce, B cell memory: building two walls of protection against pathogens. Nat. Rev. Immunol. 20, 229–238 (2020).

38. O. Bannard, J. G. Cyster, Germinal centers: programmed for affinity maturation and antibody diversification. Curr. Opin. Immunol. 45, 21–30 (2017).

39. P. A. Robert, A. L. J. Marschall, M. Meyer-Hermann, Induction of broadly neutralizing antibodies in germinal centre simulations. Curr. Opin. Biotechnol. 51, 137–145 (2018).

40. D. R. Burton, L. Hangartner, Broadly neutralizing antibodies to HIV and their role in vaccine design. Annu. Rev. Immunol. 34, 635–659 (2016).

41. L. He et al., Single-component, self-assembling, protien nanoparticles presenting the receptor binding domain and stabilized spike as SARS-CoV-2 vaccine candidates. Sci. Adv. 7, eabf1591 (2021).

42. D. Wrapp et al., Cryo-EM structure of the 2019-nCoV spike in the prefusion conformation. Science 367, 1260–1263 (2020).

43. K. H. D. Crawford et al., Protocol and reagents for pseudotyping lentiviral particles with SARS-CoV-2 spike protein for neutralization assays. Viruses 12, 513 (2020).

44. A. M. Sholukh et al., Evaluation of SARS-CoV-2 neutralization assays for antibody monitoring in natural infection and vaccine trials. medRxiv, 2020.2012.2007.20245431 (2020).

45. D. Pinto et al., Cross-neutralization of SARS-CoV-2 by a human monoclonal SARS-CoV antibody. Nature 583, 290–295 (2020).

46. A. C. Walls et al., Elicitation of broadly protective sarbecovirus immunity by receptor-binding domain nanoparticle vaccines. bioRxiv, 2021.2003.2015.435528 (2021).

47. B. Pulendran, R. Ahmed, Translating innate immunity into immunological memory: implications for vaccine development. Cell 124, 849–863 (2006).

48. S. G. Reed, M. T. Orr, C. B. Fox, Key roles of adjuvants in modern vaccines. Nat. Med. 19, 1597–1608 (2013).

49. B. Pulendran, R. Ahmed, Immunological mechanisms of vaccination. Nat. Immunol. 12, 509–517 (2011).

50. K. A. Fitzgerald, J. C. Kagan, Toll-like receptors and the control of immunity. Cell 180, 1044–1066 (2020).

51. T. Kawai, S. Akira, The role of pattern-recognition receptors in innate immunity: update on Toll-like receptors. Nat. Immunol. 11, 373–384 (2010).

52. S. P. Kasturi et al., Programming the magnitude and persistence of antibody responses with innate immunity. Nature 470, 543–547 (2011).

53. T. L. Flach et al., Alum interaction with dendritic cell membrane lipids is essential for its adjuvanticity. Nat. Med. 17, 479–487 (2011).

54. P. J. Hotez, D. B. Corry, U. Strych, M. E. Bottazzi, COVID-19 vaccines: neutralizing antibodies and the alum advantage. Nat. Rev. Immunol. 20, 399–400 (2020).

55. R. Cantisani et al., Vaccine adjuvant MF59 promotes retention of unprocessed antigen in lymph node macrophage compartments and follicular dendritic cells. J. Immunol. 194, 1717–1725 (2015).

56. G. N. Barber, STING: infection, inflammation and cancer. Nat. Rev. Immunol. 15, 760–770 (2015).

57. B. Guy, The perfect mix: recent progress in adjuvant research. Nat. Rev. Microbiol. 5, 396–397 (2007).

58. M. S. Duthie, H. P. Windish, C. B. Fox, S. G. Reed, Use of defined TLR ligands as adjuvants within human vaccines. Immunol. Rev. 239, 178–196 (2011).

59. M. P. Steinbuck et al., A lymph node–targeted Amphiphile vaccine induces potent cellular and humoral immunity to SARS-CoV-2. Sci. Adv. 7, eabe5819 (2021).

60. Y.-N. Zhang, W. Poon, E. Sefton, W. C. W. Chan, Suppressing subcapsular sinus macrophages enhances transport of nanovaccines to lymph node follicles for robust humoral immunity. ACS Nano 14, 9478–9490 (2020).

61. J. G. Liang et al., S-Trimer, a COVID-19 subunit vaccine candidate, induces protective immunity in nonhuman primates. Nat. Commun. 12, 1346 (2021).

62. P. Richmond et al., Safety and immunogenicity of S-Trimer (SCB-2019), a protein subunit vaccine candidate for COVID-19 in healthy adults: a phase 1, randomised, double-blind, placebo-controlled trial. Lancet 397, 682–694 (2021).

63. E. S. Rosenberg et al., Vigorous HIV-1-specific CD4+ T cell responses associated with control of viremia. Science 278, 1447–1450 (1997).

64. L. M. Snell et al., Overcoming CD4 Th1 cell fate restrictions to sustain antiviral CD8 T cells and control persistent virus infection. Cell Rep. 16, 3286–3296 (2016).

65. J. Zhu, H. Yamane, W. E. Paul, Differentiation of effector CD4 T cell populations (*). Annu. Rev. Immunol. 28, 445–489 (2010).

66. S. Kumar et al., Neutralizing antibodies induced by first-generation gp41-stabilized HIV-1 envelope trimers and nanoparticles. mBio 12, e00429–00421 (2021).

67. R. Shi et al., A human neutralizing antibody targets the receptor-binding site of SARS-CoV-2. Nature 584, 120–124 (2020).

68. T. F. Rogers et al., Isolation of potent SARS-CoV-2 neutralizing antibodies and protection from disease in a small animal model. Science 369, 956–963 (2020).

69. C. D. Morris et al., Differential antibody responses to conserved HIV-1 neutralizing epitopes in the context of multivalent scaffolds and native-like gp140 trimers. mBio 8, e00036–00017 (2017).

70. F. Chen et al., Functional convergence of a germline-encoded neutralizing antibody response in rhesus macaques immunized with HCV envelope glycoproteins. Immunity 54, 781-796.e784 (2021).

71. G. P. Wen et al., Quantitative evaluation of protective antibody response induced by hepatitis E vaccine in humans. Nat. Commun. 11, 3971 (2020).

72. B. Ju et al., Human neutralizing antibodies elicited by SARS-CoV-2 infection. Nature 584, 115–119 (2020).

73. Y.-N. Zhang et al., Nanoparticle size influences antigen retention and presentation in lymph node follicles for humoral immunity. Nano Lett. 19, 7226–7235 (2019).

74. B. A. Heesters, R. C. Myers, M. C. Carroll, Follicular dendritic cells: dynamic antigen libraries. Nat. Rev. Immunol. 14, 495–504 (2014).

75. J. G. Cyster, B cell follicles and antigen encounters of the third kind. Nat. Immunol. 11, 989–996 (2010).

76. R. Rappuoli, D. Serruto, Self-assembling nanoparticles usher in a new era of vaccine design. Cell 176, 1245–1247 (2019).

77. C. D. C. Allen, T. Okada, J. G. Cyster, Germinal-center organization and cellular dynamics. Immunity 27, 190–202 (2007).

78. L. Mesin, J. Ersching, Gabriel D. Victora, Germinal center B cell dynamics. Immunity 45, 471–482 (2016).

79. J. López-Sagaseta, E. Malito, R. Rappuoli, M. J. Bottomley, Self-assembling protein nanoparticles in the design of vaccines. Comput. Struct. Biotechnol. J. 14, 58–68 (2016).

80. Y. Kato et al., Multifaceted effects of antigen valency on B cell response composition and differentiation in vivo. Immunity 53, 548-563.e548 (2020).

81. S. Crotty, Follicular helper CD4 T cells (TFH). Annu. Rev. Immunol. 29, 621–663 (2011).

82. J. Merkenschlager et al., Dynamic regulation of TFH selection during the germinal centre reaction. Nature 591, 458–463 (2021).

83. C. Viant et al., Antibody affinity shapes the choice between memory and germinal center B cell fates. Cell 183, 1298-1311.e1211 (2020).

84. K. Lederer et al., SARS-CoV-2 mRNA vaccines foster potent antigen-specific germinal center responses associated with neutralizing antibody generation. Immunity 53, 1281-1295.e1285 (2020).

85. G. S. Shukla, Y. J. Sun, S. C. Pero, G. S. Sholler, D. N. Krag, Immunization with tumor neoantigens displayed on T7 phage nanoparticles elicits plasma antibody and vaccine-draining lymph node B cell responses. J. Immunol. Methods 460, 51–62 (2018).

86. L. He et al., Single-component multilayered self-assembling nanoparticles presenting rationally designed glycoprotein trimers as Ebola virus vaccines. Nat. Commun. 12, 2633 (2021).

87. L. He et al., Proof of concept for rational design of hepatitis C virus E2 core nanoparticle vaccines. Sci. Adv. 6, eaaz6225 (2020).

88. S. M. Lewis, A. Williams, S. C. Eisenbarth, Structure and function of the immune system in the spleen. Sci. Immunol. 4, (2019).

89. Y. L. Cao et al., Potent neutralizing antibodies against SARS-CoV-2 identified by high-throughput single-cell sequencing of convalescent patients’ B cells. Cell 182, 73-84.e16 (2020).

90. A. A. Cohen et al., Mosaic nanoparticles elicit cross-reactive immune responses to zoonotic coronaviruses in mice. Science 371, 735–741 (2021).

91. C. Munoz-Fontela et al., Animal models for COVID-19. Nature 586, 509–515 (2020).

92. D. S. Khoury et al., Neutralizing antibody levels are highly predictive of immune protection from symptomatic SARS-CoV-2 infection. Nat. Med., (2021).

93. R. Rappuoli et al., Vaccinology in the post−COVID-19 era. Proc. Natl. Acad. Sci. U.S.A. 118, e2020368118 (2021).

94. L. Corey, J. R. Mascola, A. S. Fauci, F. S. Collins, A strategic approach to COVID-19 vaccine R&D. Science 368, 948–950 (2020).

95. L. DeFrancesco, Whither COVID-19 vaccines? Nat. Biotechnol. 38, 1132–1145 (2020).

96. T. Tokatlian et al., Innate immune recognition of glycans targets HIV nanoparticle immunogens to germinal centers. Science 363, 649–654 (2019).

97. B. Zhang et al., A platform incorporating trimeric antigens into self-assembling nanoparticles reveals SARS-CoV-2-spike nanoparticles to elicit substantially higher neutralizing responses than spike alone. Sci. Rep. 10, 18149 (2020).

98. A. C. Walls et al., Elicitation of potent neutralizing antibody responses by designed protein nanoparticle vaccines for SARS-CoV-2. Cell 183, 1367-1382.e1317 (2020).

99. A. E. Powell et al., A single immunization with spike-functionalized ferritin vaccines elicits neutralizing antibody responses against SARS-CoV-2 in mice. ACS Cent. Sci. 7, 183–199 (2021).

100. P. J. M. Brouwer et al., Two-component spike nanoparticle vaccine protects macaques from SARS-CoV-2 infection. Cell 184, 1188-1200.e1119 (2021).

101. Y.-F. Kang et al., Rapid development of SARS-CoV-2 spike protein receptor-binding domain self-assembled nanoparticle vaccine candidates. ACS Nano 15, 2738–2752 (2021).

102. T. K. Tan et al., A COVID-19 vaccine candidate using SpyCatcher multimerization of the SARS-CoV-2 spike protein receptor-binding domain induces potent neutralising antibody responses. Nat. Commun. 12, 542 (2021).

103. X. Ma et al., Nanoparticle vaccines based on the receptor binding domain (RBD) and heptad repeat (HR) of SARS-CoV-2 elicit robust protective immune responses. Immunity 53, 1315-1330.e1319 (2020).

104. B. Nguyen, N. H. Tolia, Protein-based antigen presentation platforms for nanoparticle vaccines. npj Vaccines 6, 70 (2021).

105. I. Quast, D. Tarlinton, B cell memory: understanding COVID-19. Immunity 54, 205–210 (2021).

106. M. D. Shin et al., COVID-19 vaccine development and a potential nanomaterial path forward. Nat. Nanotechnol. 15, 646–655 (2020).

107. L. Wijaya et al., An accelerated rabies vaccine schedule based on toll-like receptor 3 (TLR3) agonist PIKA adjuvant augments rabies virus specific antibody and T cell response in healthy adult volunteers. Vaccine 35, 1175–1183 (2017).

108. C. A. Schneider, W. S. Rasband, K. W. Eliceiri, NIH Image to ImageJ: 25 years of image analysis. Nat. Methods 9, 671–675 (2012).

109. K. Mochizuki et al., Programming of donor T cells using allogeneic δ-like ligand 4-positive dendritic cells to reduce GVHD in mice. Blood 127, 3270–3280 (2016).

110. T. Tiller, C. E. Busse, H. Wardemann, Cloning and expression of murine Ig genes from single B cells. J. Immunol. Methods 350, 183–193 (2009).

