## Supplementary information for "Mechanism of a COVID-19 nanoparticle vaccine candidate that elicits a broadly neutralizing antibody response to SARS-CoV-2 variants"

### fig. S1

A **Week 5 mouse plasma neutralization against SARS-CoV-2 strains Wuhan-Hu-1, B.1.1.7, B1.351, P.1 and B.1.617$_{Rec}$**

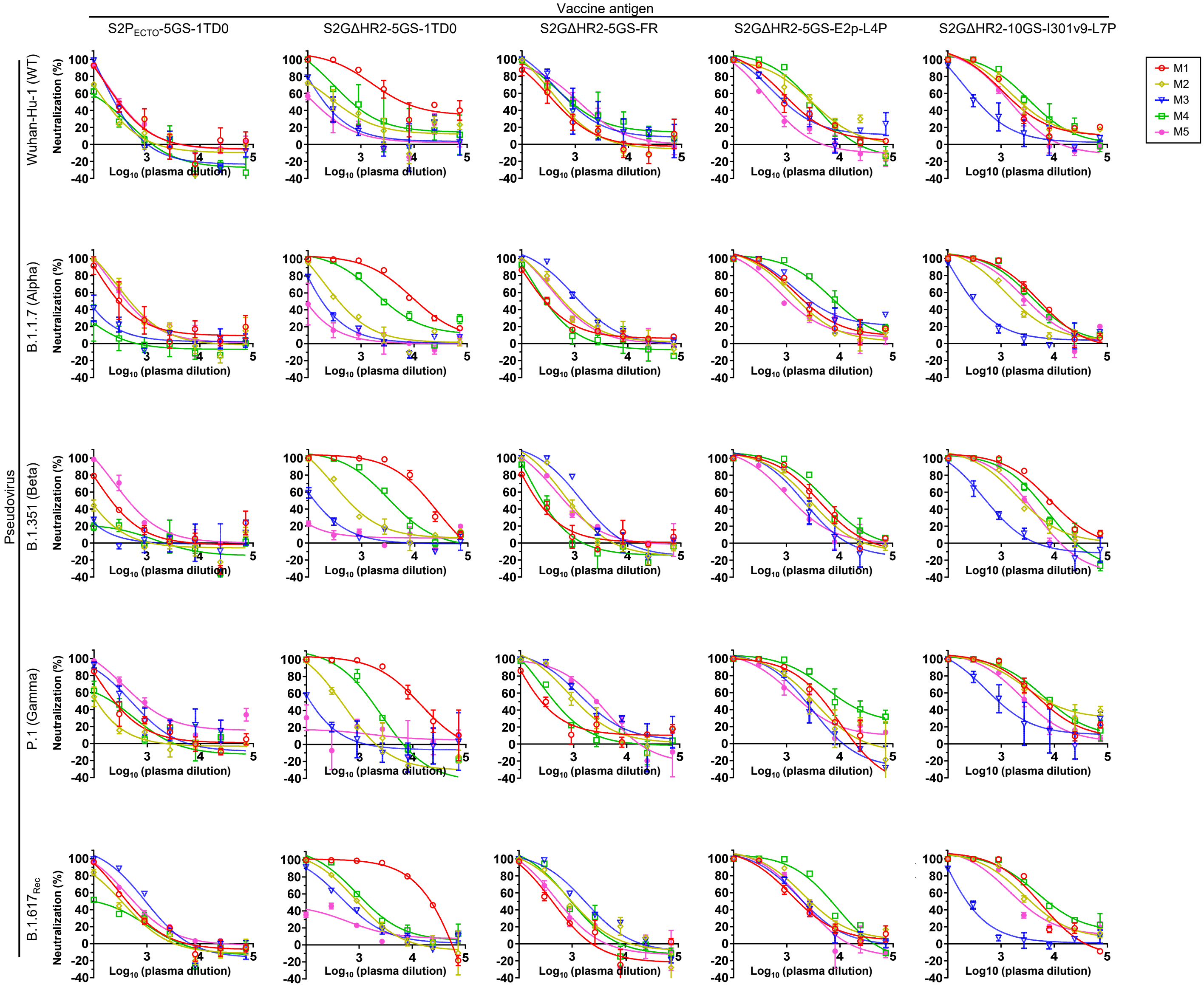


B **Week 5 mouse plasma neutralization ID$_{50}$ titers**

| | Wuhan-Hu-1 (WT) | | | | | B.1.1.7 (Alpha) | | | | | B.1.351 (Beta) | | | | | P.1 (Gamma) | | | | | B.1.617$_{Rec}$ | | | | |
|---|---|---|---|---|---|---|---|---|---|---|---|---|---|---|---|---|---|---|---|---|---|---|---|---|---|
| Vaccine antigen | M1 | M2 | M3 | M4 | M5 | M1 | M2 | M3 | M4 | M5 | M1 | M2 | M3 | M4 | M5 | M1 | M2 | M3 | M4 | M5 | M1 | M2 | M3 | M4 | M5 |
| S2PECTO-5GS-1TD0 (50 µg, i.p.) | 316 | 150 | 233 | 123 | 326 | 377 | 541 | <100 | <100 | 487 | 223 | <100 | <100 | <100 | 506 | 274 | <100 | 465 | 186 | 967 | 443 | 302 | 913 | 151 | 610 |
| S2GΔHR2-5GS-1TD0 (50 µg, i.p.) | 7719 | 370 | 227 | 911 | 131 | 11868 | 444 | 202 | 3252 | <100 | 17560 | 654 | 123 | 3550 | <100 | 12563 | 387 | 101 | 1467 | <100 | 17029 | 980 | 607 | 1530 | 124 |
| S2GΔHR2-5GS-FR (50 µg, i.p.) | 425 | 544 | 962 | 1135 | 1154 | 326 | 700 | 1327 | 293 | 629 | 215 | 764 | 1353 | 236 | 695 | 316 | 1244 | 1978 | 488 | 2355 | 379 | 994 | 1701 | 954 | 535 |
| S2GΔHR2-5GS-E2p-L4P (50 µg, i.p.) | 1442 | 3020 | 1107 | 2849 | 501 | 2458 | 1846 | 3473 | 8633 | 1331 | 4018 | 2757 | 2002 | 5971 | 1375 | 3650 | 3085 | 1966 | 14185 | 2104 | 1803 | 3045 | 2105 | 6394 | 1941 |
| S2GΔHR2-10GS-I301v9-L7P (50 µg, i.p.) | 1992 | 2648 | 428 | 3910 | 1470 | 5313 | 1953 | 355 | 4715 | 3337 | 8641 | 2602 | 506 | 4336 | 2480 | 6461 | 9520 | 1070 | 8834 | 3180 | 4969 | 4031 | 259 | 8562 | 2612 |

#### C Week 5 mouse plasma neutralization against SARS-CoV-2 strains Wuhan-Hu-1, B.1.1.7, B1.351, P.1 and B.1.617$_{Rec}$

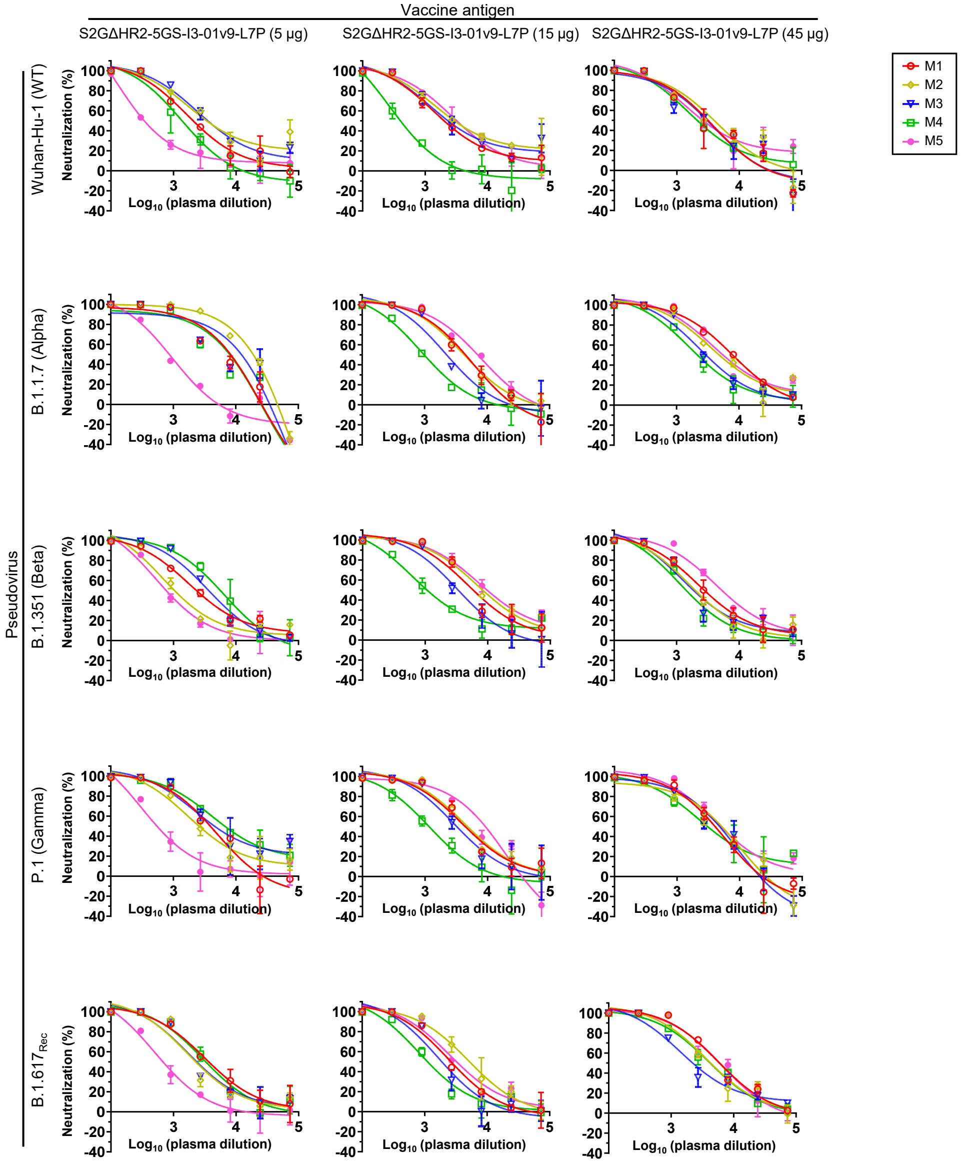


#### D Week 5 mouse plasma neutralization $ID_{50}$ titers

| | Wuhan-Hu-1 (WT) | | | | | B.1.1.7 (Alpha) | | | | | B.1.351 (Beta) | | | | | P.1 (Gamma) | | | | | B.1.617$_{Rec}$ | | | | |
|---|---|---|---|---|---|---|---|---|---|---|---|---|---|---|---|---|---|---|---|---|---|---|---|---|---|
| Vaccine antigen | M1 | M2 | M3 | M4 | M5 | M1 | M2 | M3 | M4 | M5 | M1 | M2 | M3 | M4 | M5 | M1 | M2 | M3 | M4 | M5 | M1 | M2 | M3 | M4 | M5 |
| S2GΔHR2-10GS-I3-01v9-L7P (5 µg, i.p.) | 2287 | 4102 | 4023 | 1364 | 455 | 4806 | 12463 | 5425 | 4080 | 796 | 2624 | 1163 | 3711 | 5420 | 848 | 3550 | 2939 | 5351 | 7806 | 620 | 3911 | 2467 | 2591 | 3412 | 731 |
| S2GΔHR2-10GS-I3-01v9-L7P (15 µg, i.p.) | 2381 | 3946 | 2911 | 425 | 3493 | 4050 | 4354 | 2280 | 1011 | 6750 | 6220 | 8930 | 3425 | 1290 | 10607 | 4560 | 5007 | 3325 | 1143 | 6478 | 2721 | 5353 | 1941 | 1220 | 3513 |
| S2GΔHR2-10GS-I3-01v9-L7P (45 µg, i.p.) | 2772 | 3753 | 2644 | 2433 | 3186 | 8072 | 5394 | 3222 | 2401 | 6157 | 3062 | 2141 | 2119 | 1538 | 5770 | 3627 | 3585 | 3469 | 3503 | 5235 | 6128 | 4567 | 2683 | 4285 | 5841 |

### fig. S1

E **Week 5 mouse plasma neutralization against SARS-CoV-2 strains Wuhan-Hu-1, B.1.1.7, B1.351, P.1 and B.1.617$_{Rec}$**

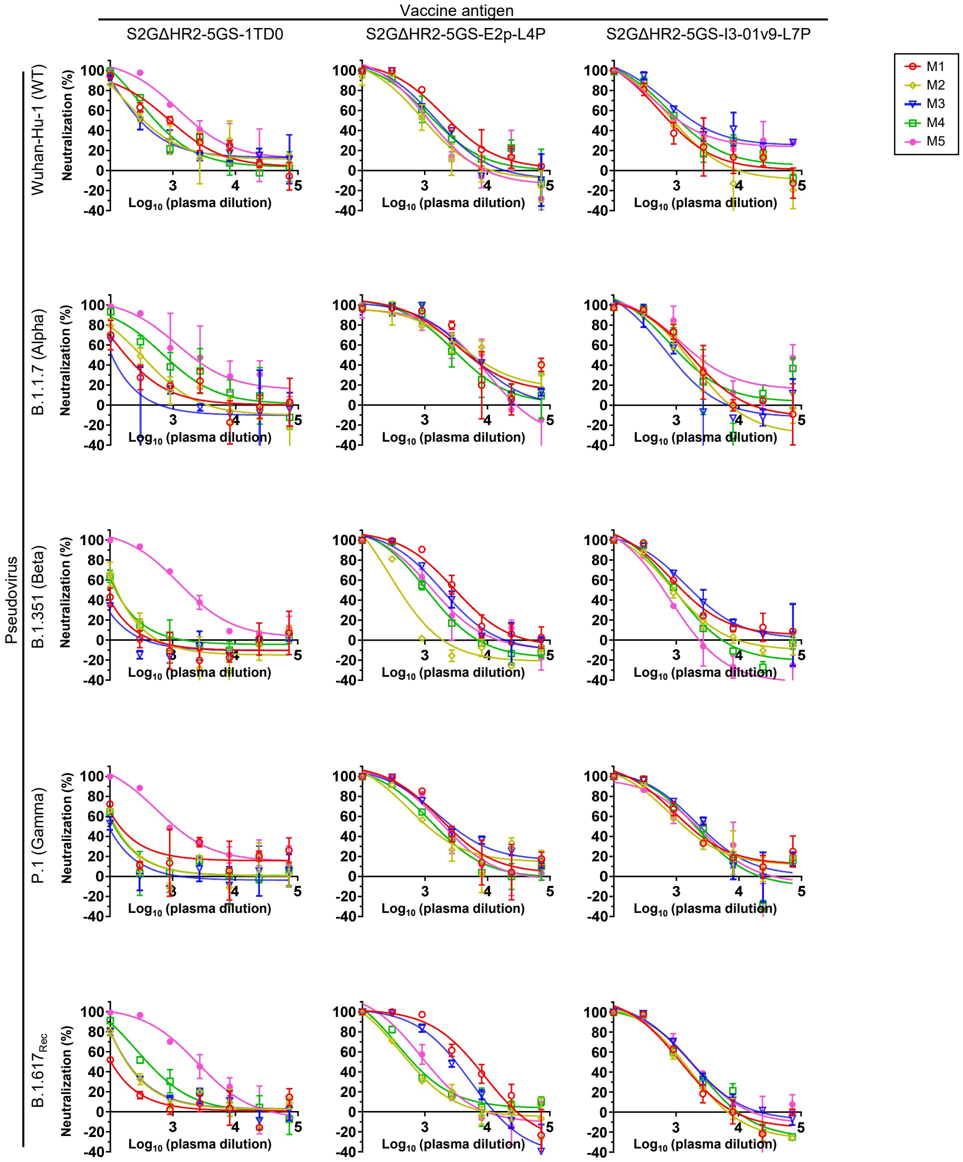


F **Week 5 mouse plasma neutralization ID$_{50}$ titers**

| | Wuhan-Hu-1 (WT) | | | | | B.1.1.7 (Alpha) | | | | | B.1.351 (Beta) | | | | | P.1 (Gamma) | | | | | B.1.617$_{Rec}$ | | | | |
|---|---|---|---|---|---|---|---|---|---|---|---|---|---|---|---|---|---|---|---|---|---|---|---|---|---|
| Vaccine antigen | M1 | M2 | M3 | M4 | M5 | M1 | M2 | M3 | M4 | M5 | M1 | M2 | M3 | M4 | M5 | M1 | M2 | M3 | M4 | M5 | M1 | M2 | M3 | M4 | M5 |
| S2GΔHR2-5GS-1TD0 (3.3 µg, i.d.) | 923 | 435 | 416 | 640 | 2345 | 179 | 278 | <100 | 721 | 2440 | <100 | <100 | <100 | 105 | 1787 | 146 | 100 | <100 | <100 | 1446 | <100 | 208 | 213 | 429 | 2329 |
| S2GΔHR2-5GS-E2p-L4P (3.3 µg, i.d.) | 2680 | 988 | 1586 | 1530 | 1178 | 5243 | 6549 | 5467 | 3676 | 4300 | 3149 | 377 | 1884 | 1048 | 1372 | 2533 | 1335 | 3524 | 1573 | 2120 | 4788 | 599 | 2385 | 734 | 1048 |
| S2GΔHR2-10GS-I3-01v9-L7P (3.3 µg, i.d.) | 851 | 884 | 2593 | 1263 | 1504 | 1617 | 1271 | 776 | 1267 | 2292 | 1418 | 921 | 1866 | 891 | 522 | 1923 | 1616 | 2530 | 1980 | 1968 | 1171 | 1140 | 1746 | 1460 | 1696 |

G **Week 26 mouse plasma neutralization against SARS-CoV-2 Wuhan-Hu-1 isolate**

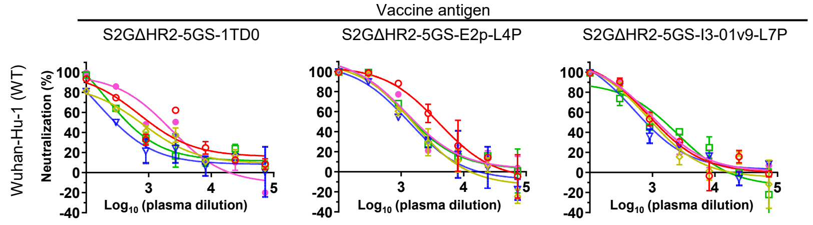


H **Week 26 mouse plasma neutralization ID$_{50}$ titers**

| | Wuhan-Hu-1 (WT) | | | | |
|---|---|---|---|---|---|
| Vaccine antigen | M1 | M2 | M3 | M4 | M5 |
| S2GΔHR2-5GS-1TD0 (3.3 µg, i.d.) | 1451 | 748 | 348 | 608 | 1502 |
| S2GΔHR2-5GS-E2p-L4P (3.3 µg, i.d.) | 3800 | 1446 | 1309 | 1728 | 1844 |
| S2GΔHR2-10GS-I3-01v9-L7P (3.3 µg, i.d.) | 1192 | 1024 | 901 | 1147 | 1323 |

**I** **Human monoclonal antibody neutralization against SARS-CoV-2 strains Wuhan-Hu-1, B.1.1.7, B1.351, P.1 and B.1.617$_{Rec}$**

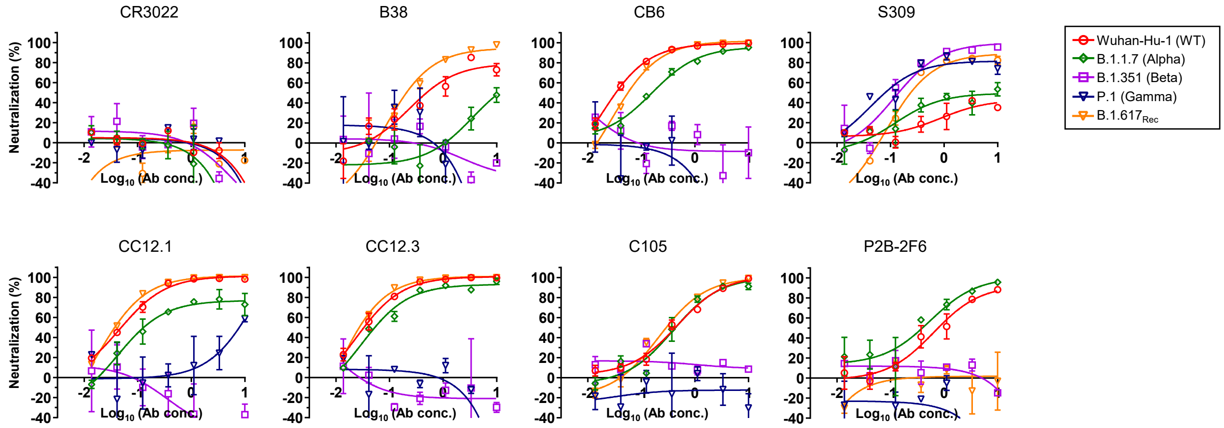


**J** **Week 5 mouse plasma neutralization against MLV-pp**

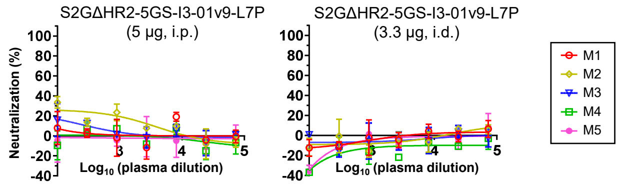


**K** **Week 5 mouse plasma neutralization ID$_{50}$ titers**

| | MLV-pp | | | | |
|---|---|---|---|---|---|
| Vaccine antigen | M1 | M2 | M3 | M4 | M5 |
| S2GΔHR2-10GS-I3-01v9-L7P (5 μg, i.p.) | <100 | <100 | <100 | <100 | <100 |
| S2GΔHR2-10GS-I3-01v9-L7P (3.3 μg, i.d.) | <100 | <100 | <100 | <100 | <100 |

**fig. S1. Spike and spike-presenting SApNP vaccine-induced neutralizing antibody responses against the wildtype SARS-CoV-2 strain and four variants.** (**A**) Neutralization curves of mouse plasma from 5 vaccine groups at week 5 after 2 intraperitoneal (i.p.) injections. The plasma samples were generated in the previous study (Ref 41), where mice were immunized with 50 μg of adjuvanted vaccine antigen. The five vaccines include two spikes (S2P$_{ECTO}$-5GS-1TD0 and S2GΔHR2-5GS-1TD0) and three SApNPs (S2GΔHR2-5GS-FR, S2GΔHR2-5GS-E2p-LD4-PADRE (L4P), and S2GΔHR2-10GS-I3-01v9-LD7-PADRE (L7P)). (**B**) Summary of ID$_{50}$ titers measured for five SARS-CoV-2 spike-based vaccine groups in (A). (**C**) Neutralization curves of mouse plasma induced by the S2GΔHR2-presenting I3-01v9 SApNP vaccine at week 5 after two i.p. injections with different antigen doses (5 μg, 15 μg, and 45 μg). (**D**) Summary of ID$_{50}$ titers measured for the S2GΔHR2-presenting I3-01v9 SApNP vaccine in (C). (**E**) Neutralization curves of mouse plasma induced by the S2GΔHR2 spike and two S2GΔHR2-presenting SApNPs (E2p and I3-01v9) at week 5 after two intradermal (i.d.) footpad injections (0.8 μg per injection site, totaling 3.3 μg per mouse). (**F**) Summary of ID$_{50}$ titers measured for the three S2GΔSHR2-based vaccines against SARS-CoV-2-pps in (E). (**G**) Neutralization curves of mouse plasma induced by the S2GΔHR2 spike and two S2GΔHR2-presenting SApNPs at week 26 after two i.d. footpad injections (0.8 μg per injection site, totaling 3.3 μg per mouse). (**H**) Summary of ID$_{50}$ titers measured for the three S2GΔSHR2-based vaccines against SARS-CoV-2-pps in (G). (**I**) Neutralization curves of human monoclonal antibodies (mAbs). In (A)-(G), SARS-CoV-2-pps that carry spikes of five strains, including the wildtype Wuhan-Hu-1 strain and four variants, B.1.1.7, B.1.351, P.1, and B.1.617$_{Rec}$ were tested in neutralization assays. (**J**) Neutralization curves of mouse plasma from two S2GΔHR2-presenting I3-01v9 SApNP vaccine groups against MLV-pps. One group was taken from (C), where mice were given 5 μg of adjuvanted antigen via i.p. injection and the other group was taken from (E), where mice were given 3.3 μg of adjuvanted antigen via i.d. injection. (**K**) Summary of ID$_{50}$ titers measured for two S2GΔHR2-presenting I3-01v9 SApNP vaccine groups against MLV-pps. In all tables, the ID$_{50}$ values were calculated with the %neutralization range constrained within 0.0-100.0%. Color coding indicates the level of ID$_{50}$ titer (white: no neutralization; green to red: low to high).

### fig. S2

**A** **Week 2 mouse plasma neutralization against SARS-CoV-2 Wuhan-Hu-1 isolate (antigen for all groups: S2GΔHR2-10GS-I3-01v9-L7P, 20 μg)**

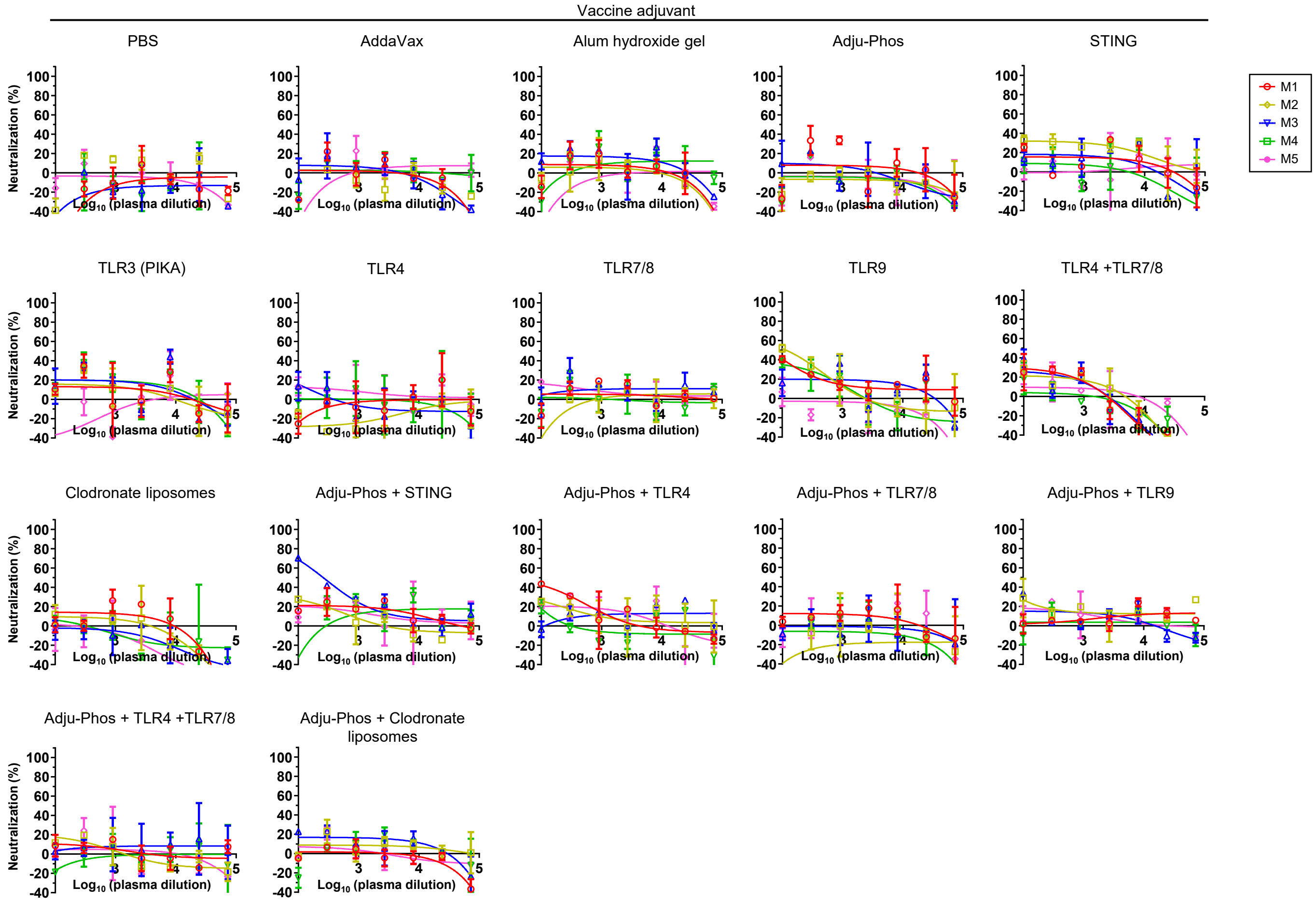


**B** **Week 2 mouse plasma neutralization $ID_{50}$ titers**

| Adjuvant | M1 | M2 | M3 | M4 | M5 | Average |
|---|---|---|---|---|---|---|
| PBS | <100 | <100 | <100 | <100 | <100 | 0 (± 0) |
| AddaVax | <100 | <100 | <100 | <100 | <100 | 0 (± 0) |
| Aluminium hydroxide gel | <100 | <100 | <100 | <100 | <100 | 6 (± 14) |
| Adju-Phos | <100 | <100 | <100 | <100 | <100 | 3 (± 8) |
| STING | <100 | <100 | <100 | <100 | <100 | 32 (± 36) |
| TLR3 (PIKA) | <100 | <100 | <100 | <100 | <100 | 26 (± 17) |
| TLR4 | <100 | <100 | <100 | <100 | <100 | 7 (± 9) |
| TLR7/8 | <100 | <100 | <100 | <100 | <100 | 7 (± 11) |
| TLR9 | <100 | 125 | <100 | <100 | <100 | 65 (± 46) |
| TLR4 +TLR7/8 | <100 | <100 | <100 | <100 | <100 | 41 (± 15) |
| Clodronate liposomes | <100 | <100 | <100 | <100 | <100 | 5 (± 6) |
| Adju-Phos + STING | <100 | <100 | 272 | <100 | <100 | 77 (± 111) |
| Adju-Phos + TLR4 | <100 | <100 | <100 | <100 | <100 | 40 (± 36) |
| Adju-Phos + TLR7/8 | <100 | <100 | <100 | <100 | <100 | 2 (± 4) |
| Adju-Phos + TLR9 | <100 | <100 | <100 | <100 | <100 | 24 (± 21) |
| Adju-Phos + TLR4 +TLR7/8 | <100 | <100 | <100 | <100 | <100 | 9 (± 8) |
| Adju-Phos + Clodronate liposomes | <100 | <100 | <100 | <100 | <100 | 10 (± 19) |

**C Week 5 mouse plasma neutralization against SARS-CoV-2 Wuhan-Hu-1 isolate (antigen for all groups: S2GΔHR2-10GS-I3-01v9-L7P, 20 μg)**

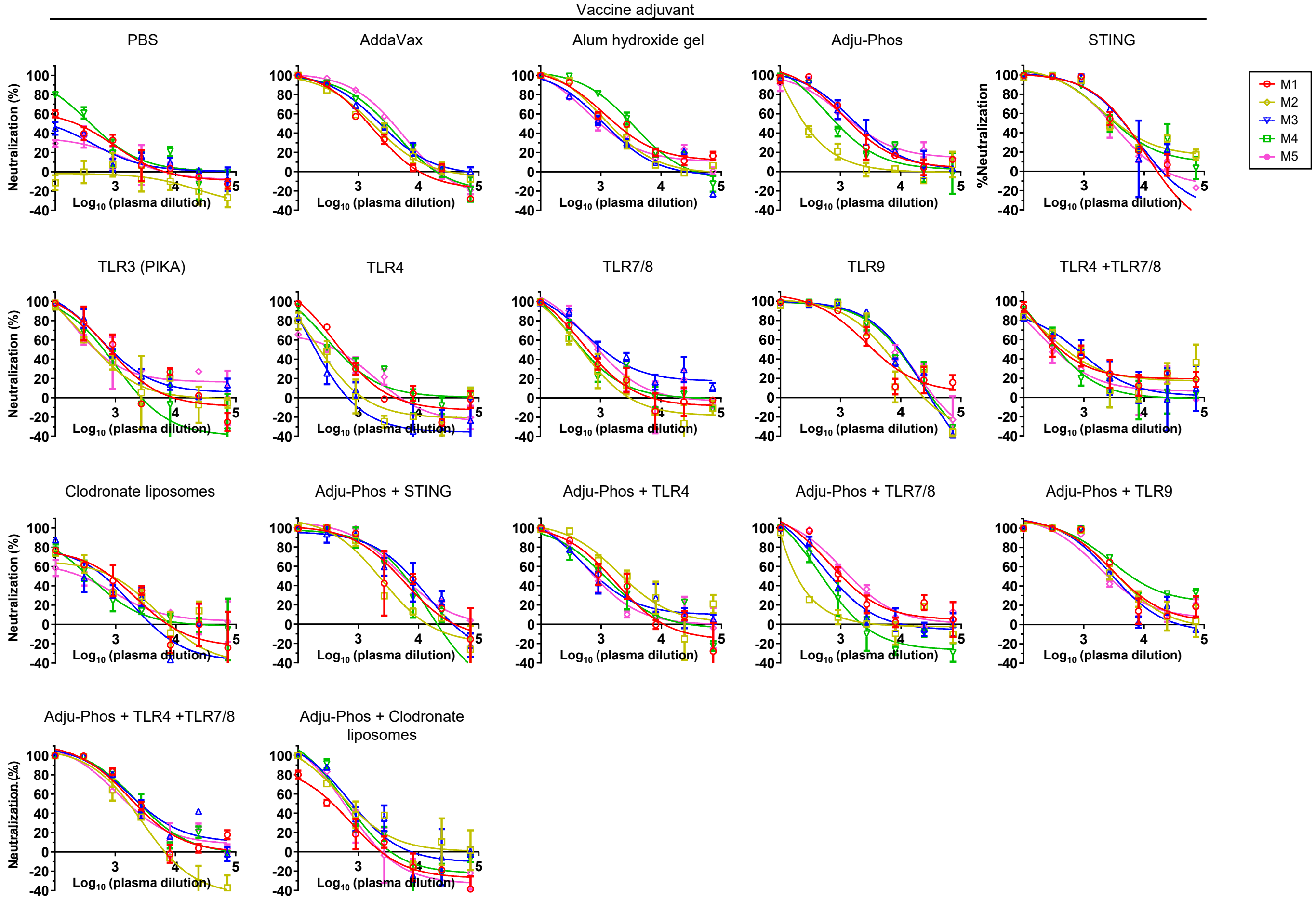


**D Week 5 mouse plasma neutralization $ID_{50}$ titers**

| Adjuvant | M1 | M2 | M3 | M4 | M5 | Average |
|---|---|---|---|---|---|---|
| PBS | 210 | 0 | 122 | 393 | 74 | 160 (± 151) |
| AddaVax | 1308 | 1630 | 2255 | 2435 | 3194 | 2164 (± 735) |
| Aluminium hydroxide gel | 2128 | 1443 | 1140 | 3145 | 1218 | 1815 (± 839) |
| Adju-Phos | 1644 | 305 | 1983 | 997 | 1949 | 1376 (± 717) |
| STING | 3204 | 4146 | 3631 | 3816 | 2715 | 3502 (± 556) |
| TLR3 (PIKA) | 799 | 525 | 971 | 493 | 722 | 702 (± 198) |
| TLR4 | 511 | 196 | 147 | 486 | 304 | 329 (± 165) |
| TLR7/8 | 633 | 453 | 1496 | 588 | 981 | 830 (± 420) |
| TLR9 | 3760 | 4344 | 6632 | 6417 | 6989 | 5628 (± 1468) |
| TLR4 +TLR7/8 | 637 | 591 | 721 | 399 | 382 | 546 (± 150) |
| Clodronate liposomes | 584 | 461 | 386 | 350 | 277 | 412 (± 117) |
| Adju-Phos + STING | 4142 | 2044 | 5268 | 3705 | 5733 | 4178 (± 1448) |
| Adju-Phos + TLR4 | 1236 | 2360 | 1060 | 1114 | 841 | 1322 (± 598) |
| Adju-Phos + TLR7/8 | 1168 | 208 | 721 | 440 | 1538 | 815 (± 540) |
| Adju-Phos + TLR9 | 4017 | 3865 | 3145 | 7203 | 2755 | 4197 (± 1758) |
| Adju-Phos + TLR4 +TLR7/8 | 2293 | 1330 | 3045 | 2677 | 1940 | 2257 (± 663) |
| Adju-Phos + Clodronate liposomes | 279 | 777 | 795 | 618 | 511 | 596 (± 213) |

### fig. S2

E **Week 8 mouse plasma neutralization against SARS-CoV-2 Wuhan-Hu-1 isolate (antigen for all groups: S2GΔHR2-5GS-I3-01v9-L7P, 20 μg)**

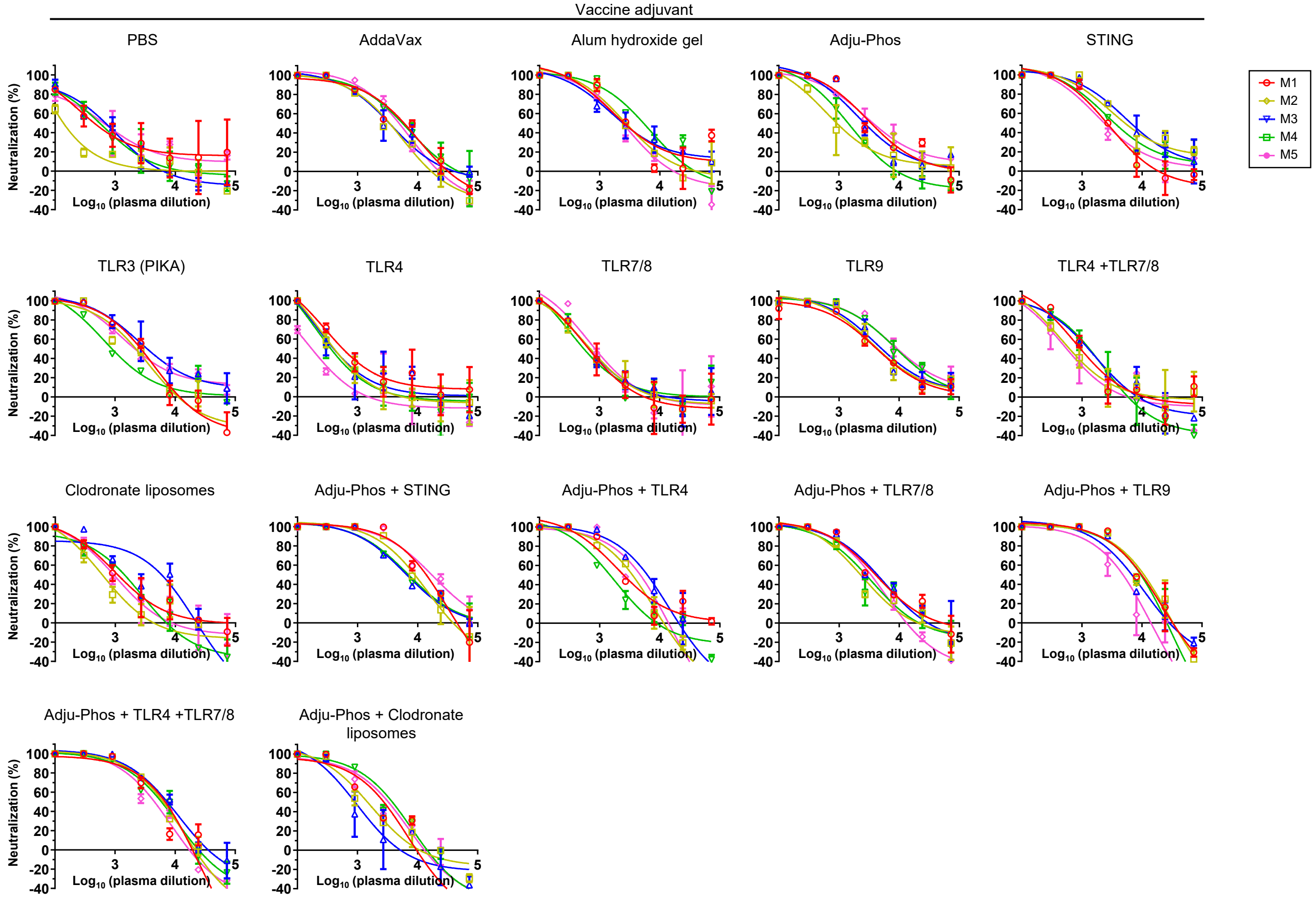


F **Week 8 mouse plasma neutralization $ID_{50}$ titers**

| Adjuvant | M1 | M2 | M3 | M4 | M5 | Average |
|---|---|---|---|---|---|---|
| PBS | 585 | 137 | 572 | 524 | 664 | 496 (± 207) |
| AddaVax | 4508 | 2682 | 3284 | 5056 | 4242 | 3954 (± 958) |
| Aluminium hydroxide gel | 2780 | 2784 | 3079 | 4805 | 2202 | 3130 (± 989) |
| Adju-Phos | 3677 | 1096 | 2867 | 1365 | 4243 | 2650 (± 1388) |
| STING | 2513 | 5913 | 7028 | 3841 | 2562 | 4371 (± 2027) |
| TLR3 (PIKA) | 2156 | 1720 | 3934 | 998 | 2658 | 2293 (± 1101) |
| TLR4 | 709 | 444 | 492 | 399 | 150 | 439 (± 200) |
| TLR7/8 | 663 | 723 | 737 | 618 | 881 | 724 (± 99) |
| TLR9 | 4337 | 4755 | 5524 | 9313 | 9989 | 6784 (± 2663) |
| TLR4 +TLR7/8 | 982 | 679 | 1117 | 1017 | 561 | 871 (± 238) |
| Clodronate liposomes | 1166 | 569 | 2467 | 989 | 886 | 1215 (± 733) |
| Adju-Phos + STING | 11103 | 7838 | 6767 | 7236 | 17251 | 10039 (± 4375) |
| Adju-Phos + TLR4 | 2705 | 2407 | 4301 | 1339 | 2617 | 2674 (± 1061) |
| Adju-Phos + TLR7/8 | 4012 | 2216 | 3825 | 2489 | 2614 | 3031 (± 825) |
| Adju-Phos + TLR9 | 7763 | 7731 | 5947 | 6229 | 2862 | 6106 (± 1997) |
| Adju-Phos + TLR4 +TLR7/8 | 3602 | 4574 | 6443 | 4607 | 3462 | 4538 (± 1190) |
| Adju-Phos + Clodronate liposomes | 1587 | 1351 | 843 | 2515 | 2041 | 1667 (± 641) |

**G** **Week 5 mouse plasma neutralization against SARS-CoV-2 strains Wuhan-Hu-1, B.1.1.7, B1.351, P.1 and B.1.617$_{Rec}$ (antigen for all groups: S2GΔHR2-5GS-I3-01v9-L7P, 20 μg)**

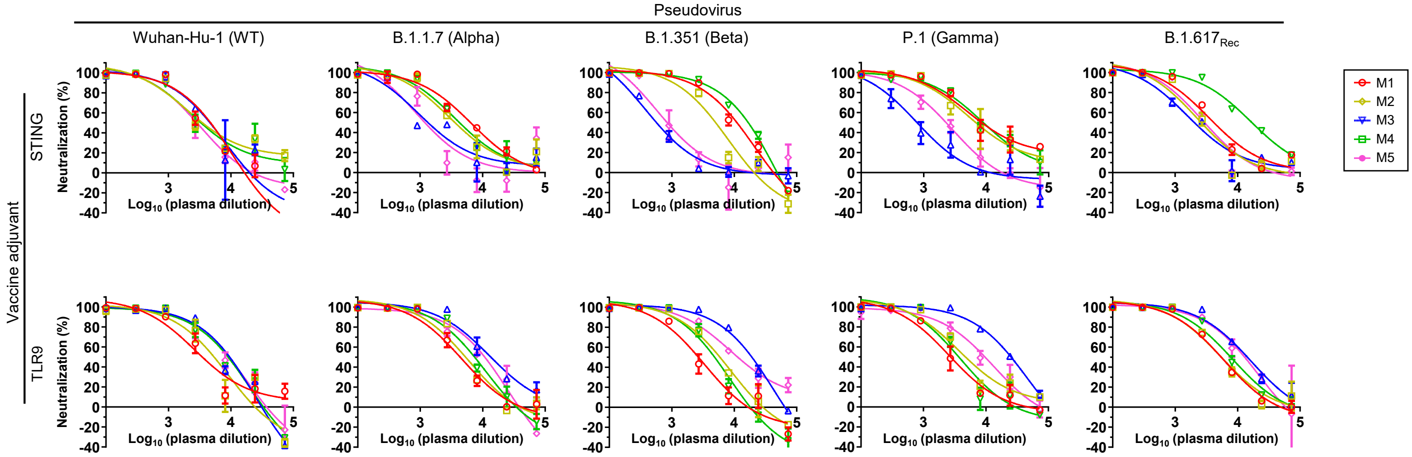


**H** **Week 5 mouse plasma neutralization $ID_{50}$ titers**

| | Wuhan-Hu-1 (WT) | | | | | B.1.1.7 (Alpha) | | | | | B.1.351 (Beta) | | | | | P.1 (Gamma) | | | | | B.1.617$_{Rec}$ | | | | |
|---|---|---|---|---|---|---|---|---|---|---|---|---|---|---|---|---|---|---|---|---|---|---|---|---|---|
| Vaccine adjuvant: | M1 | M2 | M3 | M4 | M5 | M1 | M2 | M3 | M4 | M5 | M1 | M2 | M3 | M4 | M5 | M1 | M2 | M3 | M4 | M5 | M1 | M2 | M3 | M4 | M5 |
| STING | 3204 | 4146 | 3631 | 3816 | 2715 | 6370 | 3845 | 1566 | 4559 | 1258 | 8966 | 4284 | 618 | 12385 | 902 | 9704 | 7650 | 772 | 9137 | 2034 | 4393 | 2719 | 2216 | 17438 | 3010 |
| TLR9 | 3760 | 4344 | 6632 | 6417 | 6989 | 4315 | 5234 | 13994 | 6628 | 7660 | 2420 | 5156 | 16634 | 4019 | 12345 | 2770 | 5077 | 23415 | 3615 | 8614 | 5319 | 5588 | 14219 | 7722 | 11231 |

**I** **Induction of CD4 T-cell responses against spike-presenting I3-01v9 SApNP**

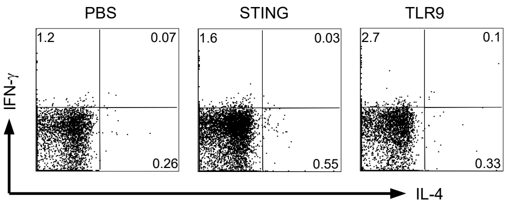


**J** **Induction of CD8 T-cell responses against spike-presenting I3-01v9 SApNP**

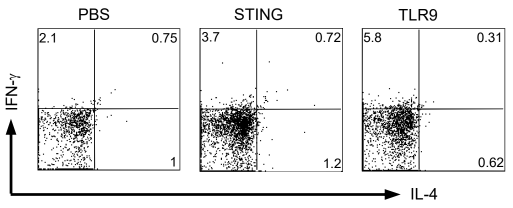


**fig. S2. Immune responses against the wildtype SARS-CoV-2 stain and four variants induced by I3-01v9 SApNP formulated with different adjuvants.** In this study, the S2GΔHR2-10GS-I3-01v9-L7P NP was either non-adjuvanted (PBS control) or formulated with various adjuvants/adjuvant mixes, resulting in 17 vaccine groups. Mice were immunized at w0, w3 and w6 through intradermal (i.d.) footpad injections (5 μg per injection site, totaling 20 μg per mouse). (**A**) Neutralization curves of mouse plasma against the wildtype Wuhan-Hu-1 strain at week 2 after single injection. (**B**) Summary of $ID_{50}$ titers measured for all vaccine groups in (A). (**C**) Neutralization curves of mouse plasma against the wildtype Wuhan-Hu-1 strain at week 5 after two injections. (**D**) Summary of $ID_{50}$ titers measured for all vaccine groups in (C). (**E**) Neutralization curves of mouse plasma against the wildtype Wuhan-Hu-1 strain at week 8 after three injections. (**F**) Summary of $ID_{50}$ titers measured for all vaccine groups in (E). (**G**) Neutralization curves of mouse plasma from the STING and TLR9 adjuvant groups against the wildtype Wuhan-Hu-1 strain and four variants, B.1.1.7, B.1.351, P.1, and B.1.617$_{Rec}$ at week 5 after two injections. (**H**) Summary of $ID_{50}$ titers measured for the two vaccine groups in (G). Representative flow cytometry graphs of (**I**) CD4 T cell, and (**J**) CD8 T cell responses against spike-presenting I3-01v9 SApNP. In all tables, the $ID_{50}$ values were calculated with the %neutralization range constrained within 0.0-100.0%. Color coding indicates the level of $ID_{50}$ titer (white: no neutralization; green to red: low to high). In (B), (D) and (F), average $ID_{50}$ titer and standard deviation are shown to facilitate the comparison of adjuvant effect between different vaccine formulation groups.

### fig. S3

A

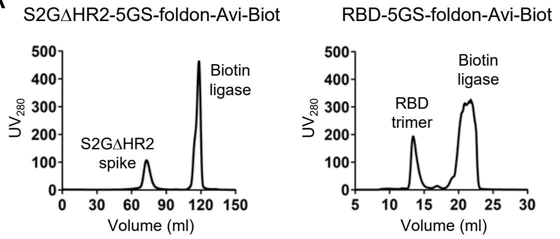


B

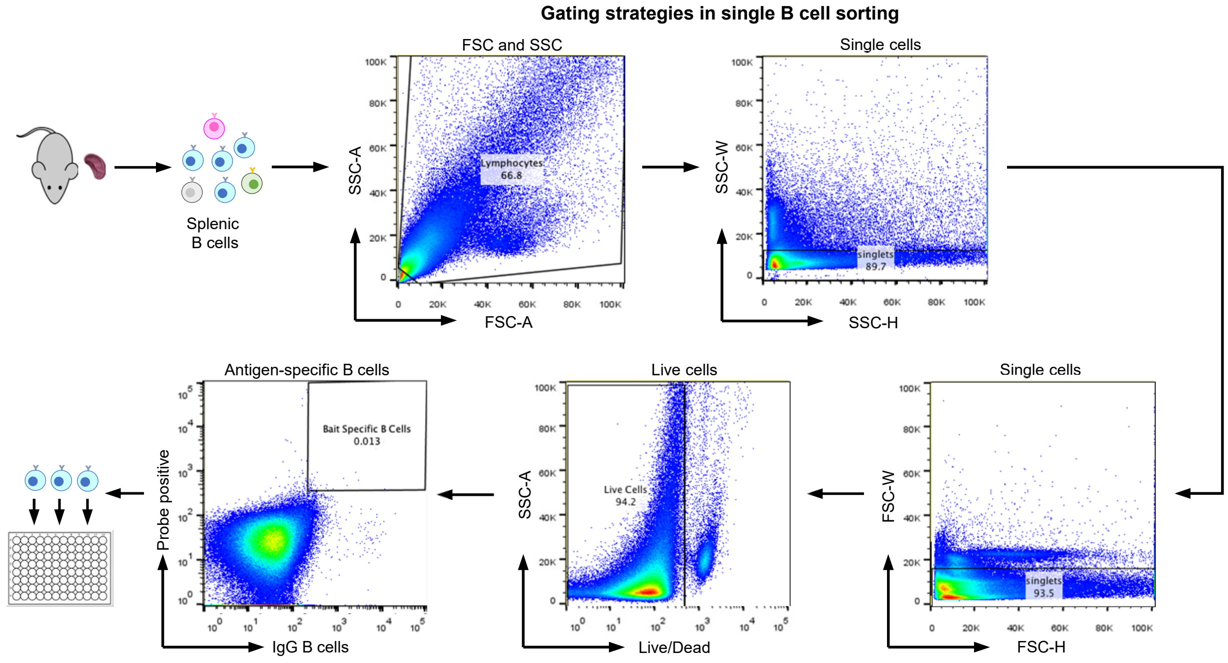


C

**Nucleotide and amino acid sequences of monoclonal neutralizing antibodies isolated from M2 in the RBD-5GS-1TD0 group [a].**

| Heavy chain | Light chain |
| --- | --- |
| **>TRBD-R-4HB6 (D-S14A-G5M2-RBD-4HB6)**<br>GAGGTGAAGCTGGAGGAGTCTGGACCTGGCCTAGTGCAGCCCTCACAGAGCCTGTCCATCACCTGCACAGTCTC<br>TGGTTTCTCATTAACTACCTATGGTGTACACTGGGTTCGCCAGTCTCCAGGAAAGGGTCTGGAGTGGCTGGGAGT<br>GATATGGAGTGGTGGAAGCACAGACTATAATGCAGCTTTCATATCCAGACTGAGCATCAGTAAGGACAATTCCAA<br>GAGCCAAGTTTTCTTTGAAATGAACAGTCTGCAAGCTAATGACACAGCCATATATTACTGTGCCAGATGGGGTTAC<br>TGGTACTTCGATGTCTGGGGCGCAGGGACCACGGTCACCGTCTCCTCA | **>TRBD-R-4KB6 (D-S14A-G5M2-RBD-4KB6)**<br>AATATTGTGCTCACCCAATCTCCATCCTCCCTGGCTATGTCAGTAGGACAGAAGGTCACTATGAGCTGCAA<br>GTCCAGTCAGAGCCTTTTAAATAGTAGCAATCAAAAGAACTATTTGGCCTGGTACCAGCAGAAACCAGGA<br>CAGTCTCCTAAAGTTCTGGTATACTTTACATCCACTAGGGAATCTGGGGTCCCTGATCGCTTCATAGGCAG<br>TGGATCTGGGACAGATTTCACTCTTACCATCAGCAGTGTGCAGGCTGAAGACCTGGCAGATTACTTCTGT<br>CAGCAACATTATAGCACTCCGCTCACGTTCGGTGCTGGGACCAAGCTGGAAATAAAA |
| **>TRBD-R-4HF3 (D-S14A-G5M2-RBD-4HF3)**<br>AAAGTGCAGCTGTTGGAGACTGGGGCTGAGCTGGTGAAGCCTGGGGCCTCAGTGAAGATGTCCTGCAAGGCTTC<br>TGGCTACACATTTACCAGTTCCAATATACACTGGATAAAGCAGACACCTGGACAGGGCCTGGAATGGATTGGAGC<br>TATTTATCCAGGAAATGGTGATACTTCCTACACTCAGAAGTTCAAAGGCAAGGCCACATTGACTGCAGACAAATCC<br>TCCAGCACAGCCTACATGCAGCTCAGCAGCCTGACATCTGAGGACTCTGCGGTCTATTACTGTGCCCTCGACGAT<br>TACTATGCTTTGGACTACTGGGGTCAAGGAACCACGGTCACCGTCTCCTCA | **>TRBD-R-4KF3 (D-S14A-G5M2-RBD-4KF3)**<br>GACATCCTGATGACCCAATCTCCATCCTCCCTAGCTGTGTCAGTTGGAGAGAAGGTTGCTATGAGCTGCA<br>AGTCCAGTCAGAGCCTTTTATACAGTAGCAATCAGAAGAACTACTTGGCCTGGTACCAGCAGAAACCAGG<br>GCAGTCTCCTAAACTGCTGATTTACTGGGCATCCACTAGGGAATCTGGGGTCCCTGATCGCTTCACAGGC<br>AGTGGATCTGGGACAGATTTCACTCTCACCATCAGCAGTGTGAAGGCTGAAGACCTGGCTGTTTATTACT<br>GTCAGCAATATTATAGCTATCCGTGGACGTTCGGTGGAGGCACCAAGCTGGAAATCAAA |
| **>TRBD-R-4HG5 (D-S14A-G5M2-RBD-4HG5)**<br>GAGGTGAAGCTGGTGGAGTCTGGGACTGTGCTGGCAAGGCCTGGGGCTTCAGTGAAGATGTCCTGCAAGGCTT<br>CTGGCTACACCTTTACCAGCTACTGGATGCACTGGGTAAAACAGAGGCCTGGACGGGGTCTGGAATGGATTGGC<br>GCTATTTATCCTGGAAATAGTGATACTAACTATAACCAGAAGTTCAAGGGCAAGGCCAAACTGACTGCAGTCACAT<br>CCACCAGCACTGCCTACATGGACCTCAGCAGCCTGACAAATGAGGACTCTGCGGTCTATTACTGTACAAGAGATT<br>ACTACGGCTACGGGAACTACTGGGGCCAAGGCACCACTCTCACAGTCTCCTCA | **>TRBD-R-4KG5 (D-S14A-G5M2-RBD-4KG5)**<br>AAAATTGTGCTCACTCAGTCTCCAGCCTCCCTATGTGCATCTGTGGGAGAAACTGTCACCATCACATGTCG<br>AGCAAGTGAGAATATTTACAGTTATTTAGCATGGTATCAGCAGAAACAGGGAAAATCTCCTCAGCTCCTGG<br>TCTATAATGCAAAAACCTTAGCAGAAGGTGTGCCATCAAGGTTCAGTGGCAGTGGATCAGGCACACAGTT<br>TTCTCTGAAGATCAACAGCCTGCAGCCTGAAGATTTTGGGAGTTATTACTGTCAACATCATTATGGTACTC<br>CGCTCACGTTCGGTGCTGGGACCAAGCTGGAAATAAAA |
| **>TRBD-R-5HB9 (D-S14A-G5M2-RBD-5HB9)**<br>GACGTGAAGCTGGTGGAGTCTGGGCCTCAGCTGGTTAGGCCTGGGGCTTCAGTGAAGATATCCTGCAAGGCTTC<br>TGGTTACTCATTCACCAGCTACTGGATGCAGTGGTTGAAGCAGAGGCCTGGACAAGGTCTTGAGTGGATTGGCAT<br>GATTGATCCTTCCGATAGTGAAACTAGGTTAAATCAGAAGTTCAAGGACAAGGCCACATTGACTGTAGACAAATCC<br>TCCAGCACAGCCCACATGCAACTCAGCAGCCCGACATCTGAGGACTCTGCGGTCTATTATTGTGCAAGAAGGGA<br>TTACTACGGCAATAATCCCTTTGACTTCTGGGGCCAAGGCACCACTCTCACAGTCTCCTCA | **>TRBD-R-5KB9 (D-S14A-G5M2-RBD-5KB9)**<br>GACATTGTGATGTCACAGTCTCCACCCTCTGTACCTGTCACTCCTGGAGAGTCAGTATCCATCTCCTGCAG<br>GTCTAATAAGAGTCTCCTGCATAGTAATGGCAACACTTACTTGTATTGGTTCCTGCAGAGGCCAGGCCAGT<br>CTCCTCAGCTCCTGATATTTCGGATGTCCAACCTTGCCTCAGGAGTCCCAGACAGGTTCAGTGGCAGTGG<br>GTCAGGAACTGCTTTCACACTGAGAATCAGTAGAGTGGAGGCTGAGGATGTGGGTGTTTATTACTGTATG<br>CAACATCTAGAATATCCGTACACGTTCGGAGGGGGGACCAAGCTGGAAATAAAA |
| **>TRBD-R-4HB6 (D-S14A-G5M2-RBD-4HB6)**<br>EVKLEESGPGLVQPSQSLSITCTVSGFSLTTYGVHWVRQSPGKGLEWLGVIWSGGSTDYNAAFISRLSISKDNSKSQV<br>FFEMNSLQANDTAIYYCARWGYWYFDVWGAGTTVTVSS | **>TRBD-R-4KB6 (D-S14A-G5M2-RBD-4KB6)**<br>NIVLTQSPSSLAMSVGQKVTMSCKSSQSLLNSSNQKNYLAWYQQKPGQSPKVLVYFTSTRESGVPDRFIGSG<br>SGTDFTLTISSVQAEDLADYFCQQHYSTPLTFGAGTKLEIK |
| **>TRBD-R-4HF3 (D-S14A-G5M2-RBD-4HF3)**<br>KVQLLETGAELVKPGASVKMSCKASGYTFTSSNIHWIKQTPGQGLEWIGAIYPGNGDTSYTQKFKGKATLTADKSSST<br>AYMQLSSLTSEDSAVYYCALDDYYALDYWGQGTTVTVSS | **>TRBD-R-4KF3 (D-S14A-G5M2-RBD-4KF3)**<br>DILMTQSPSSLAVSVGEKVAMSCKSSQSLLYSSNQKNYLAWYQQKPGQSPKLLIYWASTRESGVPDRFTGSG<br>SGTDFTLTISSVKAEDLAVYYCQQYYSYPWTFGGGTKLEIK |
| **>TRBD-R-4HG5 (D-S14A-G5M2-RBD-4HG5)**<br>EVKLVESGTVLARPGASVKMSCKASGYTFTSYWMHWVKQRPGRGLEWIGAIYPGNSDTNYNQKFKGKAKLTAVTST<br>STAYMDLSSLTNEDSAVYYCTRDYYGYGNYWGQGTTLTVSS | **>TRBD-R-4KG5 (D-S14A-G5M2-RBD-4KG5)**<br>KIVLTQSPASLCASVGETVTITCRASENIYSYLAWYQQKQGKSPQLLVYNAKTLAEGVPSRFSGSGSGTQFSLK<br>INSLQPEDFGSYYCQHHYGTPLTFGAGTKLEIK |
| **>TRBD-R-5HB9 (D-S14A-G5M2-RBD-5HB9)**<br>DVKLVESGPQLVRPGASVKISCKASGYSFTSYWMQWLKQRPGQGLEWIGMIDPSDSETRLNQKFKDKATLTVDKSSS<br>TAHMQLSSPTSEDSAVYYCARRDYYGNNPFDFWGQGTTLTVSS | **>TRBD-R-5KB9 (D-S14A-G5M2-RBD-5KB9)**<br>DIVMSQSPPSVPVTPGESVSISCRSNKSLLHSNGNTYLYWFLQRPGQSPQLLIFRMSNLASGVPDRFSGSGS<br>GTAFTLRISRVEAEDVGVYYCMQHLEYPYTFGGGTKLEIK |

[a] Single-cell sorted mouse antibodies are named as TRBD-[Probe]-[Antibody index].TRBD stands for trimeric RBD. For this mouse, RBD-Avi-Biot was used as a sorting probe.

### fig. S3

D

**Nucleotide and amino acid sequences of monoclonal neutralizing antibodies isolated from M4 in the S2GΔHR2-5GS-1TD0 group [a].**

| Heavy chain | Light chain |
|---|---|
| **>S2GD-S-1HF3 (D-S14A-G7M4-1HF3)**<br>GAGGTGCAGCTGCAGCAGTCTGGGGCTGAACTGGCAAAACCTGGGGCCTCAGTGAAGATGTCCTGCAAGGCTTC<br>TGGCTACACCTTTACTAGCTACTGGGTGCACTGGGTAAAACAGAGGCCTGGACAGGGTCTGGAATGGATTGGATA<br>CATTTATCCTAGCACTGGTTATACTGACTACAATCAGAAGTTCAAGGACAAGGCCACATTGACTGCAGACAAATCC<br>TCCACCACAGCCTACATGCAACTGAGCAGCCTGACATCTGACCACTCTGCAGTCTATTACTGTGCAAGATCCGAG<br>TATGGTAACCTTTACTATGCTATGGACTACTGGGGTCAAGGAACCTCAGTCACCGTCTCCTCA | **>S2GD-S-1KF3 (D-S14A-G7M4-1KF3)**<br>CAAATTGTTCTCTCCCAGTCTCCAGCTTCTTTGGCTGTGTCTCTAGGGCAGAGGGCCACCATCTCCTGCAA<br>GGCCAGCCAAAGTGTTGATTATGATGGTGATAGTTATATGAACTGGTACCAACAGAAACCAGGACAGCCA<br>CCCAAACTCCTCATCTATTCTGCATCCAATCTAGAATCTGGGATCCCAGCCAGGTTTAGTGGCAGTGGGCC<br>TGGGACAGACTTCACCCTCAACATCCATCCTGTGGAGGAGGAGGATGCTGCAACCTATTACTGTCAACAA<br>AGTAATGAGGATCCTCTCACGTTCGGTGCTGGGACCAAGCTGGAAATAAAA |
| **>S2GD-S-2HC10 (D-S14A-G7M4-2HC10)**<br>CAGCGTGAGCTGCAGCAGTCTGGACCTGAGCTGGTAAAGCCTGGGGCTTCAGTGAAGATGTCCTGCAAGGCTTC<br>TGGATACACATTCACTAACTATGTTATGCACTGGGTGAAGCAGAAGCCTGGGCAGGGCCTTGAGTGGATTGGCTA<br>TATTAATCCTTACAATGATGGTACTAAGTACAATGAGAAGTTCAAAGGCAAGGCCACACTGACTTTAGACAAATCC<br>TCCAGCACAGCCTACATGGAGCTCAGCAGCCTGACCTCTGAGGACTCTGCGGTCTATTACTGTGCAAGATTTACT<br>AAGGTAGAGGGCTACTGGGGCCAAGGCTCCACTCTCACAGTCTCCTCA | **>S2GD-S-2KC10 (D-S14A-G7M4-2KC10)**<br>GACATCTTGCTGACTCAGTCTCCAGCTTCTTTGGCTGTGTCTCTAGGGCAGAGGGCCACCATCTCCTGCA<br>GAGCCAGCGAAAGTGTTGATAATTATGGCATTAGTCTTATGAACTGGTTCCAACAGAAACCAGGACAGCCA<br>CCCCAACTCCTCATCTATGCTGCATCCAACCAAGGATCCGGGGTCCCTGCCAGGTTTAGTGGCAGTGGGT<br>CTGGGACAGACTTCAGCCTCAACATCCACCCTATGGAGGAGGATGATACTGCAATGTATTTCTGTCACCAA<br>AGTAAGGAGGTTCCGTGGACGTTCGGTGGAGGCACCAAGCTGGAAATCAAACGT |
| **>S2GD-R-1HF9 (D-S14A-G7M4-RBD-1HF9)**<br>GAAGTGATGCTGGTGGAGTCTGGGGCTGAGCTTGTGAGGCCAGGGGCCTTAGTCAAGTTGTCCTGCAAAGCTTC<br>TGGCTTCAACATTAAAGACTACTATATGCAGTGGGTGAAGCAGAGGCCTGAACAGGGCCTGGAGTGGATTGGATG<br>GATTGATCCTGAGAATGGTAATACTATATATGACCCGAAGTTCCAGGGCAAGGCCAGTATAACAGCAGACACATC<br>CTCCAACACAGCCTACCTGCAGCTCAGCAGCCTGACATCTGAGGACACTGCCGTCTATTACTGTGCCCTCTGGGA<br>GGGGAGAGCTATGGACTACTGGGGTCAAGGAACCTCAGTCACCGTCTCCTCA | **>S2GD-R-1KF9 (D-S14A-G7M4-RBD-1KF9)**<br>GACATCCAGATGACTCAGTCTCCATCCTCCCTGTCCGCCTCTCTGGGAGACAGAGTCACCATCAGTTGCA<br>GTGCAAGTCAGGGCATCAGCAATTATTTAAATTGGTATCAGCAGAAACCAGATGGAACTGTTAAACTCCTG<br>ATCTATTACACATCAAGCTTACACTCAGGAGTCCCATCAAGGTTCAGTGGCAGTGGGTCTGGGACAGATTA<br>TTCTCTCGCCATCAGCAACCTGGAACCTGAAGATATTGTCACTTACTATTGTCAGCAGTACAGTAAACTTCC<br>GTACACGTTCGGAGGGGGGACCAAGCTGGAGCTGAAA |
| **>S2GD-R-2HD10 (D-S14A-G7M4-RBD-2HD10)**<br>GAAGTGATGCTGGTGGAGTCTGGGGCAGAGCTTGTGAAGCCAGGGGCCTCAGTCAAGTTGTCCTGCACAGCTTC<br>TGGCTTCAACATTAAAGACACCTTTATACACTGGGTGAAACAGAGGCCTGAACAGGGCCTGGAGTGGATTGGAGG<br>GATTGATCCTGCGAATGCTGATCCTAAATATGACCCGAAGTTTCAGGGCAAGGCCACCATAACAGCAGACACATC<br>CTCCAACACAGCCTACCTGCAGTTCAGCAGCCTGACATCTGAGGACACTGCCGTCTATTACTGTGCTCGATGGGA<br>TAATGCGGCCTATTACTATGGTATGGACTACTGGGGTCAAGGCACCTCAGTCACCGTCTCCTCA | **>S2GD-R-2KD10 (D-S14A-G7M4-RBD-2KD10)**<br>GACATCCTGATGACCCAATCTCCATCTTCCATGTATGCATCTCTAGGAGAAAGAGTCACTTTCACCTGCAA<br>GGCGAGTCAGGACATTAATAGGTTTTTAAACTGGTTCCAGCAGAAACCAGGGAAATCTCCTAAGACCCTGA<br>TCTATCGTGCAAACAGATTGGTAGATGGGGTCCCATCAAGGTTCAGTGGCAGTGGATCTGGGCAAGATTA<br>TTCTCTCACCATCAGCAGTTTGGAGTATGAAGATATGGGAATTTATTATTGTCTGCAGTATGATGAGTTGTA<br>CACGTTCGGAGGGGGGACCAAGCTGGAAATAAAA |
| **>S2GD-R-2HE4 (D-S14A-G7M4-RBD-2HE4)**<br>GAGGTGAAGCTGGAGGAGTCTGGGGCTGAACTGGCGAAGCCTGAGGCTTCAGTGAAGTTGTCCTGCAAGGCTTC<br>TGGCTACACCTTCACCAGCTACTTTATGTACTGGGTGAAGCAGAGGCCTGGACAAGGCCTTGAGTGGATTGGACA<br>GATTAACCCTACCAATGGTGGTACTAACTTCAATGAGAAGTTCAAGAGCAAGGCCACACTGACTGTAGACAAATC<br>CTCCAGCACAGCATACATGCAACTCAGCAGCCTGACATCTGAGGACTCTGCGGTCTATTACTGTACAATATATGG<br>AAGGTACTTTGACTGCTGGGGCCAAGGCACCACTCTCACAGTCTCCTCA | **>S2GD-R-2KE4 (D-S14A-G7M4-RBD-2KE4)**<br>GACATCCAGATGAACCAGTCTCCATCCTCTATGTCTGCATCTCTGGGAGGCAGAATAACCATCACTTGCCA<br>GGCAACTCAAGACATTGTTAAGAATTTAAACTGGTATCAGCAGAAACCAGGGAAACCCCCTTCATTCCTGA<br>TCTATTATGCAACTGAACTGGCGGAAGGGGTCCCATCAAGGTTCAGTGGCAGTGGGTCTGGGTCAGACTA<br>TTCTCTGACAATCAGCAACCTGGAGTCTGAAGATTTTGCAGACTATTACTGTCTACAGTTTTACGAGTTTCC<br>GTACACGTTCGGAGGGGGGACCAAGCTGGAAATAAAA |
| **>S2GD-R-2HF4 (D-S14A-G7M4-RBD-2HF4)**<br>GAAGTGATGCTGGTGGAGTCTGGACCTGAGTTGGTGAAGCCTGGGGCTTCAGTGAAGATATCCTGCAAGGCCTC<br>TGGTTACTCATTTACTGGCTACTTTATGAGCTGGGTGAAGCAGAGCCGTGGAAAGAGCCTTGAGTGGATTGGACG<br>TATTAATCCTAACAATGGTGATACTTTCTACAACCAGAAGTTCAAGGGCAAGGCCACATTGACTGTAGACAAATCC<br>TCTAGCACAGCCCACATGGACTTCCTGAGCCTGACATCTGAGGACTCTGCAGTCTATTATTGTGTAAGAGAGAGG<br>CATTACTGGGGCCAAGGCACCACTCTCACAGTCTCCTCA | **>S2GD-R-2KF4 (D-S14A-G7M4-RBD-2KF4)**<br>GAAACAACTGTGACCCAGTCTCCATCCTCCATGTCTGTATCTCTGGGAGACACAGTCAGCATCACTTGCCA<br>TGCAAGTCAGGGCATTAGCAGTAATATAGGGTGGTTGCAGCAGAAACCAGGGAAATCATTTAAGGGCCTG<br>ATCTATCATGGAACCAACTTGGAAGATGGAGTTCCATCAAGGTTCAGTGGCAGTGGATCTGGAGCAGATTA<br>TTCTCTCACCATCAGCAGCCTGGAGTCTGAAGATTTTGCAGACTATTACTGTGTACAGTATGATCAGTTTCC<br>GTACACGTTCGGAGGGGGGACCAAGCTGGAGCTGAAA |
| **>S2GD-S-1HF3 (D-S14A-G7M4-1HF3)**<br>EVQLQQSGAELAKPGASVKMSCKASGYTFTSYWVHWVKQRPGQGLEWIGYIYPSTGYTDYNQKFKDKATLTADKSS<br>TTAYMQLSSLTSDHSAVYYCARSEYGNLYYAMDYWGQGTSVTVSS | **>S2GD-S-1KF3 (D-S14A-G7M4-1KF3)**<br>QIVLSQSPASLAVSLGQRATISCKASQSVDYDGDSYMNWYQQKPGQPPKLLIYSASNLESGIPARFSGSGPGT<br>DFTLNIHPVEEEDAATYYCQQSNEDPLTFGAGTKLEIK |
| **>S2GD-S-2HC10 (D-S14A-G7M4-2HC10)**<br>QRELQQSGPELVKPGASVKMSCKASGYTFTNYVMHWVKQKPGQGLEWIGYINPYNDGTKYNEKFKGKATLTLDKSS<br>STAYMELSSLTSEDSAVYYCARFTKVEGYWGQGSTLTVSS | **>S2GD-S-2KC10 (D-S14A-G7M4-2KC10)**<br>DILLTQSPASLAVSLGQRATISCRASESVDNYGISLMNWFQQKPGQPPQLLIYAASNQGSGVPARFSGSGSGT<br>DFSLNIHPMEEDDTAMYFCHQSKEVPWTFGGGTKLEIKR |
| **>S2GD-R-1HF9 (D-S14A-G7M4-RBD-1HF9)**<br>EVMLVESGAELVRPGALVKLSCKASGFNIKDYYMQWVKQRPEQGLEWIGWIDPENGNTIYDPKFQGKASITADTSSNT<br>AYLQLSSLTSEDTAVYYCALWEGRAMDYWGQGTSVTVSS | **>S2GD-R-1KF9 (D-S14A-G7M4-RBD-1KF9)**<br>DIQMTQSPSSLSASLGDRVTISCSASQGISNYLNWYQQKPDGTVKLLIYYTSSLHSGVPSRFSGSGSGTDYSLA<br>ISNLEPEDIVTYYCQQYSKLPYTFGGGTKLELK |
| **>S2GD-R-2HD10 (D-S14A-G7M4-RBD-2HD10)**<br>EVMLVESGAELVKPGASVKLSCTASGFNIKDTFIHWVKQRPEQGLEWIGGIDPANADPKYDPKFQGKATITADTSSNTA<br>YLQFSSLTSEDTAVYYCARWDNAAYYYGMDYWGQGTSVTVSS | **>S2GD-R-2KD10 (D-S14A-G7M4-RBD-2KD10)**<br>DILMTQSPSSMYASLGERVTFTCKASQDINRFLNWFQQKPGKSPKTLIYRANRLVDGVPSRFSGSGSGQDYSL<br>TISSLEYEDMGIYYCLQYDELYTFGGGTKLEIK |
| **>S2GD-R-2HE4 (D-S14A-G7M4-RBD-2HE4)**<br>EVKLEESGAELAKPEASVKLSCKASGYTFTSYFMYWVKQRPGQGLEWIGQINPTNGGTNFNEKFKSKATLTVDKSSST<br>AYMQLSSLTSEDSAVYYCTIYGRYFDCWGQGTTLTVSS | **>S2GD-R-2KE4 (D-S14A-G7M4-RBD-2KE4)**<br>DIQMNQSPSSMSASLGGRITITCQATQDIVKNLNWYQQKPGKPPSFLIYYATELAEGVPSRFSGSGSGSDYSLT<br>ISNLESEDFADYYCLQFYEFPYTFGGGTKLEIK |
| **>S2GD-R-2HF4 (D-S14A-G7M4-RBD-2HF4)**<br>EVMLVESGPELVKPGASVKISCKASGYSFTGYFMSWVKQSRGKSLEWIGRINPNNGDTFYNQKFKGKATLTVDKSSS<br>TAHMDFLSLTSEDSAVYYCVRERHYWGQGTTLTVSS | **>S2GD-R-2KF4 (D-S14A-G7M4-RBD-2KF4)**<br>ETTVTQSPSSMSVSLGDTVSITCHASQGISSNIGWLQQKPGKSFKGLIYHGTNLEDGVPSRFSGSGSGADYSL<br>TISSLESEDFADYYCVQYDQFPYTFGGGTKLELK |

[a] Single-cell sorted mouse antibodies are named as S2GD-[Probe]-[Antibody index].S2GD stands for the S2GΔHR2 spike, or the full construct name S2GΔHR2-5GS-1TD0. Probe can be S, which stands for spike, or R, which stands for RBD. For this mouse, both RBD-Avi-Biot and S2GΔHR2-5GS-foldon-Avi-Biot were used as sorting probes.

### fig. S3

E

**Nucleotide and amino acid sequences of monoclonal neutralizing antibodies isolated from M2 in the S2GΔHR2-10GS-I3-01v9-LD7-PADRE group [a].**

| Heavy chain | Light chain |
|---|---|
| **>I3V9-S-1HC9 (D-S15G2M2-1HC9)**<br>CAGGTGCAGCTGCAGCAGTCTGGGGCTGAGCTTGTGAGGCCAGGGGCCTTAGTCAAGTTGTCCTGCAAAGCTTC<br>TGGCTTCAACATTAAAGACTACTATATACACTGGGTGAAACAGAGGCCTGAACAGGGCCTGGAGTGGATTGGATG<br>GATTGATCCTGAGAATGGTAATGCTATATATGACCCGAAGTTCCAGGGCAAGGCCAGTATAACAGCAGACACATC<br>CTCCAACACAGCCTACCTGCAGCTCAGCAGCCTGACATCTGAGGACACTGCCGTCTATTACTGTGCTAGGGGGG<br>ACTTTGACTACTGGGGCCAAGGCACCACTCTCACAGTCTCCTCA | **>I3V9-S-1KC9 (D-S15G2M2-1KC9-2)**<br>GATGTTGTGATGACCCAAACTCCAGCAATCATGTCTGCATCTCCAGGGGAGAAGGTCACCATGACCTGCA<br>GTGCCAGCTCAAGTGTAGGTTACATGCACTGGTACCAGCAGAAGCCAGGATCCTCCCCCAGACTCCTGAT<br>TTATGACGCATCCAACCTGGCTTCTGGAGTCCCTGTTCGTTTCAGTGGCAGTGGGTCTGGGACCTCTTACT<br>CTCTCACAATCAGCCGAATGGAGGCTGAAGATGCTGCCACTTATTACTGCCAGCAGTGGGGTACTTACCC<br>ACCCAGGACGTTCGGTGGAGGCACCAAGCTGGAAATCAAA |
| **>I3V9-S-2HD8 (D-S15G2M2-2HD8)**<br>CAGGTGCAGCTGCAGCAGCCTGGGGCAGAGCTTGTGAAGCCGGGGGCCTCAGTCAAGTTGTCCTGCACAGCTT<br>CTGGCTTCAACATTAAAGACACCTATATTCACTGGGTGAATCAGAGGCCTGAACAGGGCCTGGAGTGGATTGGAA<br>GGATTGATCCTGCGAATGGTAATACTAAGTATGACCCTAAATTCCAGGGCAAGGCCACAATAACAGCAGACACCT<br>CCTCCAATACCGCCTACCTGCAGCTCAGCAGCCTGACATCTGAGGACACTGCCGTCTATTACTGTGCTGGAGGAC<br>TCTATGATTACGACGGGAGCCCTTTTGCCTACTGGGGCCAAGGCACCACGATCACCGTCTCCTCA | **>I3V9-S-2KD8 (D-S15G2M2-2KD8)**<br>GATATCCAGATGACACAGACTACATCCTCCCTGTCTGCCTCTCTGGGAGACAGAGTCACCATCAGTTGCA<br>GGGCAAGTCAGGGCATTAGCACTTATTTACACTGGTATCAGCAGAAACCAGATGGAACTCTTAAACTCCTG<br>ATCTACTACACATCAAGATTACACTCAGGTGTCCCGTCAAGGTTCAGTGGCAGTGGGTCTGGAACAGATTA<br>TTCTCTCACCATTAGCAACCTGGAACAAGAGGATGTTGCCACTTACTTTTGCCAACAGGGTAATACGCTTC<br>CGTACACGTTCGGCGGGGGGACCAAGCTGGAGCTGAAA |
| **>I3V9-S-2HD10 (D-S15G2M2-2HD10-2)**<br>GAGGTGAAGCTGGAGGAGTCTGGGGCTGAGCTTGTGAGGCCAGGGGCCTTGGTCAAGTTGTCCTGCAAAGCTTC<br>TGGCTTCAACATTAAAGACTACTATATACACTGGGTGAAGCAGAGGCCTGAACAGGGCCTGGAGTGGATTGGATG<br>GATTGATCCTGTTTATGGTAATTCTATATGTGACCCGAAGTTTCAGGGCAAGGCCACTATAACAGCAGACACATCC<br>TCCAACACAGCCTACCTGCAGCTCAGCAGCCTGACATCTGAGGACACTGCCGTCTATTACTGTGCCATGGGGGAT<br>GGTAACTACTGGGGCCAGGGCACCACTCTCACAGTCTCCTCA | **>I3V9-S-2KD10 (D-S15G2M2-2KD10-2)**<br>GACATCAAGATGACCCAATCTCCAGCAATCATGTCTGCATCTCCAGGGGAGAAGGTCACCATGACCTGCA<br>GTGCCACCTCAAGTGTAGATTACATGCACTGGTTCCAGCAGAAGCCAGGCACTTCTCCCAAACTCTGGATT<br>TATAGCACATCCAACCTGGCTTCTGGAGTCCCTGCTCGCTTCAGTGGCAGCGGATCTGGGACCTCTTACT<br>CTCTCACAATCAGCCGAATGGAGGCTGAAGATGCTGCCACTTATTACTGCCAGCAAAGGAGTAGTTACCC<br>AATATTCACGTTCGGCTCGGGGACCAAGCTGGAGCTGAAA |
| **>I3V9-R-1HG3 (D-S15G2M2-RBD-1HG3)**<br>AAGTTCCAGCTGCAGCAGTCTGGGGCTGAAATGGTGAAGCCTGGGGCTTCAGTGAAGCTGTCCTGCAAGGCTTC<br>TGGCTACACCTTCACCAACTACTGGATGTACTGGATGAAGCAGAGGCCTGGGCAAGGCCTTGAGTGGATTGGAG<br>AGATTAATCCTAGGAACGGTCTTACTAACTACAATGAGAAATTCAAGACCAAGGCCACACTGACTGTAGACAAACC<br>CTCCAGCACAGCCTTCATGCAGCTCAACAGCCTGACATCTGAGGACTCTGCGGTCTATTACTGTGCAAAGGACGG<br>TAGTAGCGCCTACTGGGGCCAAGGCACCACTCTCACAGTCTCCTCA | **>I3V9-R-1KG3 (D-S15G2M2-RBD-1KG3)**<br>GATGTTGTGGTGACTCAAACTCCATCCTCCTTATCTGCCTCTCTAGGAGAAAGAGTCAGTCTCACTTGTCG<br>GGCAAGTCAGGACATTGGTAGTAGCTTAAACTGGCTTCAGCAGGAACCAGATGGAACTATTAAACGCCTG<br>ATCTACGCCACATCCATTTTAGATTCTGGTGTCCCCAAAAGGTTCAGTGGCAGTAGGTCTGGGTCAGATTA<br>TTCTCTCACCATCAGCAGCCTTGAGTCTGAAGACTTTGTAGACTATTACTGTCTACAATATGCTGGTTCTCC<br>ATTCACGTTCGGCTCGGGGACAAAGTTGGAAATAAAA |
| **>I3V9-R-1HG9 (D-S15G2M2-RBD-1HG9)**<br>CAGGTGCAGCTGAAGGAGTCTGGACCTGAGCTGGTGAAGCCTGGGGCTTCAGTGAAGATATCCTGCAAGACTTC<br>TGGATACACATTCACTGAATACACCATGCTCTGGGTGAAGCAGAGCCATGGAAAGAGCCTTAAGTGGATTGGAGG<br>TATTAATCCTAACAATGGTGATACTACCTACAACCAGAAGTTCAAGGGCAAGTTTATATTGACTGTAGACAAGTCCT<br>CCAGCACAGCCTACATGGAGCTCCGCAGCCTGACATCTGAGGATTCTGCAGTCTATTACTGTGCAAGAGATGGTT<br>ACCCCTATTACTATGCTATGGACTACTGGGGTCAAGGAACCTCAGTCACCGTCTCCTCA | **>I3V9-R-1KG9 (D-S15G2M2-RBD-1KG9-1)**<br>TACATCAAGATGACCCAGTCTCCAAAATTCATGTCCACATCAGTAGGAGACAGGGTCAGCGTCACCTGCA<br>AGGCCAGTCAGAATGTGGGTACTAATGTAGCCTGGTATCAAAAGAAACCAGGGCAATCTCCTAAAGCACT<br>GATTTACTCGGCATCCTACCGGTACAGTGGAGTCCCTGATCGCTTCACAGGCAGTGGATCTGGGACAGAT<br>TTCACTCTCACCATCAGCAATGTGCAGTCTGAAGACTTGGCAGAGTATTTCTGTCAACAATATAACAGCTAT<br>CCGTGGACGTTCGGTGGAGGCACCAAGCTGGAAATCAAA |
| **>I3V9-R-1HF5 (D-S15G2M2-RBD-1HF5)**<br>CAGGTGCAGCTGAAGGAGTCTGGACCTGAGCTGGTGAAGCCTGGGGCCTCAGTGAAGATTTCCTGCAAAGTTTC<br>TGGTTCCGCATTCAGTAATTCTTGGATGAACTGGGTGAAGCAGAGGCCTGGACAGGGTCTTGAATGGATTGGACG<br>GATTTATCTTGGAGATGGGGATACTAACTTCAATGGGAAATTCAAGGACAAGGCCACACTGACTGCAGACAAATC<br>CTCCACCACAGCCTACATGCAACTCAGCAGCCTCACCTCTGTGGACTCTGCGGTCTATTTCTGTGCAAATTCCTG<br>GGACGGTCTGGTGTTTGCTTACTGGGGCCAGGGGACTCTGGTCACTGTCTCTGCA | **>I3V9-R-1KF5 (D-S15G2M2-RBD-1KF5)**<br>AACATTGTGATGTCACAGTCTCCAACAATCATGTCTGCATCTCCAGGGGAGAAGGTCACTTTGACCTGCAG<br>TGCCAGCTCAGGTGTTACTTACATCTTCTGGTACCAGCAGAAGCCAGGATCCTCCCCCAGACTCCTGATTT<br>ATGACACATCCATCCTGGCTTCTGGAGTCCCTGTTCGCTTCAGTGGCGGTGGGTCTGGGACCTCTTACTC<br>TCTCACACTCAGCCGAATGGAGGCTGAAGATGCTGCCACTTATTACTGCCAGCAGTGGAATAATTTCCCAC<br>CCACGTTCGGTCCTGGGACCAAGCTGGAGCTGAAA |
| **>I3V9-R-2HE11 (D-S15G2M2-RBD-2HE11)**<br>GAGGTGCAGCTGGTGGAGTCTGGGGCTGAACTGGTGAAGCCTGGGGCTTCAGTGAAGTTGTCCTGCAAGGCTTC<br>TGGCTACAGCTTTACCACCTACTATATATACTGGATGAAGCAGAGGCCTGGACAGGGCCTTGAGTGGATTGGAGA<br>GATTAATCCTAGCAATGGTGGTACTAACTTCAATGAGAGGTTCAAGAGCAAGGCCACACTGACTGTAGACAAATC<br>CTCCAGCACAGCATACATGCAACTCAGCAGCCTGACATCTGAGGACTCTGCGGTCTATTACTGTTCAAGAGATGG<br>TAGTATTGCTTACTGGGGCCAAGGGACTCTGGTCACTGTCTCTGCA | **>I3V9-R-2KE11 (D-S15G2M2-RBD-2KE11-4)**<br>GAAAATGTTCTCACCCAGTCTCCATCCTCCTTATCTGCCTCTCTGGGAGAAAGAGTCAGTCTCACTTGTCG<br>GGCAAGTCAGGACACTGGTAGTAGTTTAAACTGGCTTCAGCAGGAACCAGATGGAACTATTAAACGCCTG<br>ATCTACGCCACATCCAATTTAGATTCTGGTGTCCCCAAAAGGTTCAGTGGCAGTAGGTCTGGGTCAGATTA<br>TTCTCTCACCATCAGCAGCCTTGAGTCTGAAGATTTTGTAGACTATTACTGTCTACAATATGCTAGTTCTCC<br>GTACACGTTCGGAGGGGGGACCAAGCTGGAAATAAAA |
| **>I3V9-R-2HF2 (D-S15G2M2-RBD-2HF2)**<br>GAGGTGAAGCTGGAGGAGTCTGGACCTGAGCTGGTGAAGCCTGGGGCTTCAGTGAAGATTTCCTGCAAGACTTC<br>TGGATTCACATTCACTGAATACACCATGCACTGGGTGAAGCAGAGCCATGGAAAGAGCCTTGAGTGGATTGGAGC<br>TATTAATCTTAACATTGTTGATACCATCTACAATCAGAACTTCCAGGGCAAGGCCACATTTGCTGTAGACAAGTCCT<br>CCAGCACAGCCTACATGGAACTCCGTAGCCTGACATCTGAAGATTCTTCAGTCTACTACTGCGCAAGGGATAGGT<br>ACGACCGGTACTTCGATGTCTGGGGCGCAGGGACCACGGTCACCGTCTCCTCA | **>I3V9-R-2KF2 (D-S15G2M2-RBD-2KF2)**<br>TACATTGTGATGACTCAGTCTCCAACGATCATGTCTGCGTCTCCAGGGGAGAAGGTCACTTTGACCTGCAG<br>TGCCAGCTCAGGTGTTACTTACATCTTCTGGTACCAGCAGAAGCCAGGATCCTCCCCCAGACTCCTGATTT<br>ATGACACATCCATCCTGGCTTCTGGAGTCCCTGTTCGCTTCAGTGGCGGTGGGTCTGGGACCTCTTACTC<br>TCTCACACTCAGCCGAATGGAGGCTGAAGATGCTGCCACTTATTACTGCCAGCAGTGGAATAATTTCCCAC<br>CCACGTTCGGTCCTGGGACCAAGCTGGAAATAAAA |
| **>I3V9-R-2HF5 (D-S15G2M2-RBD-2HF5)**<br>CAGGCTTATCTACAGCAGTCTGGACCTGAGCTGGTGAAGCCTGGGGCTTCAGTGAAGATATCCTGCAAGGCTTCT<br>GGTTACTCTTTTACTGAATACTTTATGAACTGGGTGATGCAGAGCCATGGAAAGAGCCTTGAGTGGATTGGACGTA<br>TTAATCCTTACAATGGTGATACTTTCTACAACCAGAAGTTCAAGGGCAAGGCCACATTGACTGTAGACAAATCATC<br>TAGCACTGCCCACATGGAGTTCCGGAGCCTGGCATCTGAGGACTCCGCAGTCTATTATTGTGCAAGATCCCATGA<br>TTACCCCTTTGACTACTGGGGCCAAGGCACCACTCTCACAGTCTCCTCA | **>I3V9-R-2KF5 (D-S15G2M2-RBD-2KF5)**<br>GACATCCTGATGACCCAATCTCCATCGTCCATGTTTGCCTCTCTGGGAGACAGAGTCAGTCTCTCTTGTCG<br>GGCTAGTCAGGGCATTAGAGGTAATTTAGACTGGTATCAGCAGAAACCAGGTGGAACTATTAAACTCCTGA<br>TCTACTCCACATTTAATTTAAATTTTGGTGTCCCATCAAGGTTCAGTGGCAGTGGGTCTGGGTCAGATTATT<br>CTCTCACCATCAGCAGCCTAGAGTCTGAAGATTTTGCAGACTATTACTGTCTACAGCGTAATGCGTATCCG<br>TACACGTTCGGAGGGGGGACAAAGTTGGAAATAAAA |
| **>I3V9-R-2HF10 (D-S15G2M2-RBD-2HF10)**<br>GAGTTCCAGCTGCAGCAGTCTGGACCTGAACTGGTGAGGCCTGGGGCTTCAGTGAAGATATCCTGCAGGACTTC<br>TGGATACACATTCACTGAATACACCCTGTACTGGGTGAAGCAGAGCCATGGAAAGAGCCTTGAGTGGATTGGAGG<br>TATTAATCCTAACAATGGTGATACTACCTACAACCAGAGGTTTAAGGGCAGGGCCACATTGACTGTAGACAAGTCA<br>TCCAGCACAGCCTACATGGAGCTCCGCAGCCTGACATCTGAGGATTCTGCAGTCTATTACTGTGCGAGAGATGGT<br>TACCCCTATTACTATGCCTTGGACTATTGGGGTCAAGGAACCACTCTCACAGTCTCCTCA | **>I3V9-R-2KF10 (D-S15G2M2-RBD-2KF10-2)**<br>TACATCTTGCTGACTCAGTCTCCAGCCTCTCAGTCTGCATCTCTGGGAGAAAGTGTCACCATCACATGCCT<br>GGCAAGTCAGACCATTGGTACATGGTTAGCATGGTATCAGCAGAAACCAGGGAAATCTCCTCAGCTCCTG<br>ATTTATGCTGCAACCAGCTTGGCAGATGGGGTCCCATCAAGGTTCAGTGGTAGTGGATCTGGCACAAAATT<br>TTCTTTCAAGATCAGCAGCCTACAGGCTGAAGATTTTGTAAGTTATTACTGTCAACAACTTTACAGTACTCC<br>TCTGACGTTCGGTGGAGGCACCAAGCTGGAAATCAAA |
| **>I3V9-S-1HC9 (D-S15G2M2-1HC9)**<br>QVQLQQSGAELVRPGALVKLSCKASGFNIKDYYIHWVKQRPEQGLEWIGWIDPENGNAIYDPKFQGKASITADTSSNT<br>AYLQLSSLTSEDTAVYYCARGDFDYWGQGTTLTVSS | **>I3V9-S-1KC9 (D-S15G2M2-1KC9-2)**<br>DVVMTQTPAIMSASPGEKVTMTCSASSSVGYMHWYQQKPGSSPRLLIYDASNLASGVPVRFSGSGSGTSYSL<br>TISRMEAEDAATYYCQQWGTYPPRTFGGGTKLEIK |
| **>I3V9-S-2HD8 (D-S15G2M2-2HD8)**<br>QVQLQQPGAELVKPGASVKLSCTASGFNIKDTYIHWVNQRPEQGLEWIGRIDPANGNTKYDPKFQGKATITADTSSNT<br>AYLQLSSLTSEDTAVYYCAGGLYDYDGSPFAYWGQGTTITVSS | **>I3V9-S-2KD8 (D-S15G2M2-2KD8)**<br>DIQMTQTTSSLSASLGDRVTISCRASQGISTYLHWYQQKPDGTLKLLIYYTSRLHSGVPSRFSGSGSGTDYSLTI<br>SNLEQEDVATYFCQQGNTLPYTFGGGTKLELK |
| **>I3V9-S-2HD10 (D-S15G2M2-2HD10-2)**<br>EVKLEESGAELVRPGALVKLSCKASGFNIKDYYIHWVKQRPEQGLEWIGWIDPVYGNSICDPKFQGKATITADTSSNTA<br>YLQLSSLTSEDTAVYYCAMGDGNYWGQGTTLTVSS | **>I3V9-S-2KD10 (D-S15G2M2-2KD10-2)**<br>DIKMTQSPAIMSASPGEKVTMTCSATSSVDYMHWFQQKPGTSPKLWIYSTSNLASGVPARFSGSGSGTSYSL<br>TISRMEAEDAATYYCQQRSSYPIFTFGSGTKLELK |
| **>I3V9-R-1HG3 (D-S15G2M2-RBD-1HG3)**<br>KFQLQQSGAEMVKPGASVKLSCKASGYTFTNYWMYWMKQRPGQGLEWIGEINPRNGLTNYNEKFKTKATLTVDKPS<br>STAFMQLNSLTSEDSAVYYCAKDGSSAYWGQGTTLTVSS | **>I3V9-R-1KG3 (D-S15G2M2-RBD-1KG3)**<br>DVVVTQTPSSLSASLGERVSLTCRASQDIGSSLNWLQQEPDGTIKRLIYATSILDSGVPKRFSGSRSGSDYSLTI<br>SSLESEDFVDYYCLQYAGSPFTFGSGTKLEIK |
| **>I3V9-R-1HG9 (D-S15G2M2-RBD-1HG9)**<br>QVQLKESGPELVKPGASVKISCKTSGYTFTEYTMLWVKQSHGKSLKWIGGINPNNGDTTYNQKFKGKFILTVDKSSST<br>AYMELRSLTSEDSAVYYCARDGYPYYYAMDYWGQGTSVTVSS | **>I3V9-R-1KG9 (D-S15G2M2-RBD-1KG9-1)**<br>YIKMTQSPKFMSTSVGDRVSVTCKASQNVGTNVAWYQKKPGQSPKALIYSASYRYSGVPDRFTGSGSGTDFT<br>LTISNVQSEDLAEYFCQQYNSYPWTFGGGTKLEIK |
| **>I3V9-R-1HF5 (D-S15G2M2-RBD-1HF5)**<br>QVQLKESGPELVKPGASVKISCKVSGSAFSNSWMNWVKQRPGQGLEWIGRIYLGDGDTNFNGKFKDKATLTADKSST<br>TAYMQLSSLTSVDSAVYFCANSWDGLVFAYWGQGTLVTVSA | **>I3V9-R-1KF5 (D-S15G2M2-RBD-1KF5)**<br>NIVMSQSPTIMSASPGEKVTLTCSASSGVTYIFWYQQKPGSSPRLLIYDTSILASGVPVRFSGGGSGTSYSLTLS<br>RMEAEDAATYYCQQWNNFPPTFGPGTKLELK |
| **>I3V9-R-2HE11 (D-S15G2M2-RBD-2HE11)**<br>EVQLVESGAELVKPGASVKLSCKASGYSFTTYYIYWMKQRPGQGLEWIGEINPSNGGTNFNERFKSKATLTVDKSSST<br>AYMQLSSLTSEDSAVYYCSRDGSIAYWGQGTLVTVSA | **>I3V9-R-2KE11 (D-S15G2M2-RBD-2KE11-4)**<br>ENVLTQSPSSLSASLGERVSLTCRASQDTGSSLNWLQQEPDGTIKRLIYATSNLDSGVPKRFSGSRSGSDYSL<br>TISSLESEDFVDYYCLQYASSPYTFGGGTKLEIK |
| **>I3V9-R-2HF2 (D-S15G2M2-RBD-2HF2)**<br>EVKLEESGPELVKPGASVKISCKTSGFTFTEYTMHWVKQSHGKSLEWIGAINLNIVDTIYNQNFQGKATFAVDKSSSTA<br>YMELRSLTSEDSSVYYCARDRYDRYFDVWGAGTTVTVSS | **>I3V9-R-2KF2 (D-S15G2M2-RBD-2KF2)**<br>YIVMTQSPTIMSASPGEKVTLTCSASSGVTYIFWYQQKPGSSPRLLIYDTSILASGVPVRFSGGGSGTSYSLTLS<br>RMEAEDAATYYCQQWNNFPPTFGPGTKLEIK |
| **>I3V9-R-2HF5 (D-S15G2M2-RBD-2HF5)**<br>QAYLQQSGPELVKPGASVKISCKASGYSFTEYFMNWVMQSHGKSLEWIGRINPYNGDTFYNQKFKGKATLTVDKSSS<br>TAHMEFRSLASEDSAVYYCARSHDYPFDYWGQGTTLTVSS | **>I3V9-R-2KF5 (D-S15G2M2-RBD-2KF5)**<br>DILMTQSPSSMFASLGDRVSLSCRASQGIRGNLDWYQQKPGGTIKLLIYSTFNLNFGVPSRFSGSGSGSDYSL<br>TISSLESEDFADYYCLQRNAYPYTFGGGTKLEIK |
| **>I3V9-R-2HF10 (D-S15G2M2-RBD-2HF10)**<br>EFQLQQSGPELVRPGASVKISCRTSGYTFTEYTLYWVKQSHGKSLEWIGGINPNNGDTTYNQRFKGRATLTVDKSSST<br>AYMELRSLTSEDSAVYYCARDGYPYYYALDYWGQGTTLTVSS | **>I3V9-R-2KF10 (D-S15G2M2-RBD-2KF10-2)**<br>YILLTQSPASQSASLGESVTITCLASQTIGTWLAWYQQKPGKSPQLLIYAATSLADGVPSRFSGSGSGTKFSFKI<br>SSLQAEDFVSYYCQQLYSTPLTFGGGTKLEIK |

[a] Single-cell sorted mouse antibodies are named as I3V9-[Probe]-[Antibody index].I3V9 stands for the I3-01v9 nanoparticle, or the full construct name S2GΔHR2-10GS-I3-01v9-LD7-PADRE. Probe can be S, which stands for spike, or R, which stands for RBD. For this mouse, both RBD-Avi-Biot and S2GΔHR2-5GS-foldon-Avi-Biot were used as sorting probes.

### fig. S3

**F Mouse antibody neutralization against SARS-CoV-2 strains Wuhan-Hu-1, B.1.1.7, B1.351, P.1 and B.1.617$_{Rec}$**

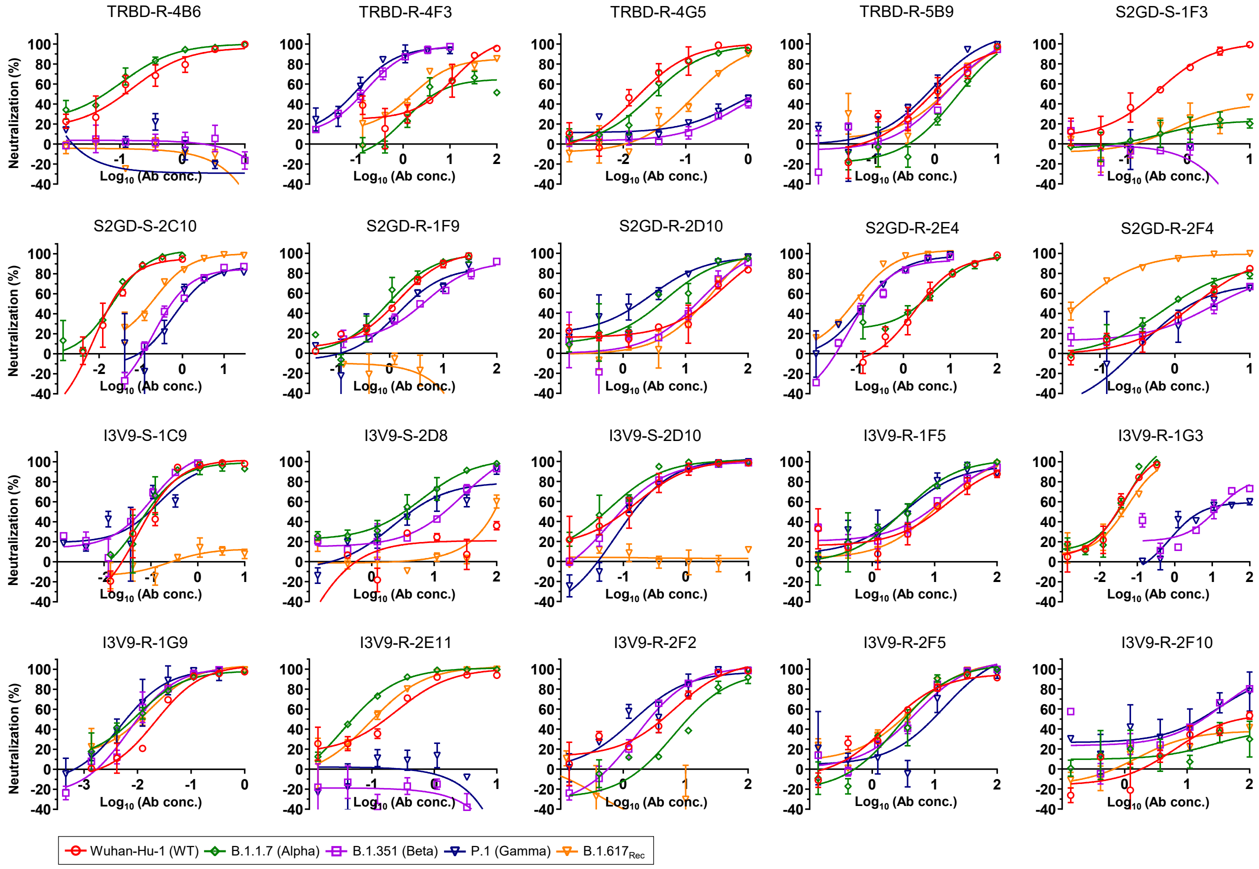


**G Mouse antibodies isolated from mice in three vaccine groups binding to RBD and spike antigens**

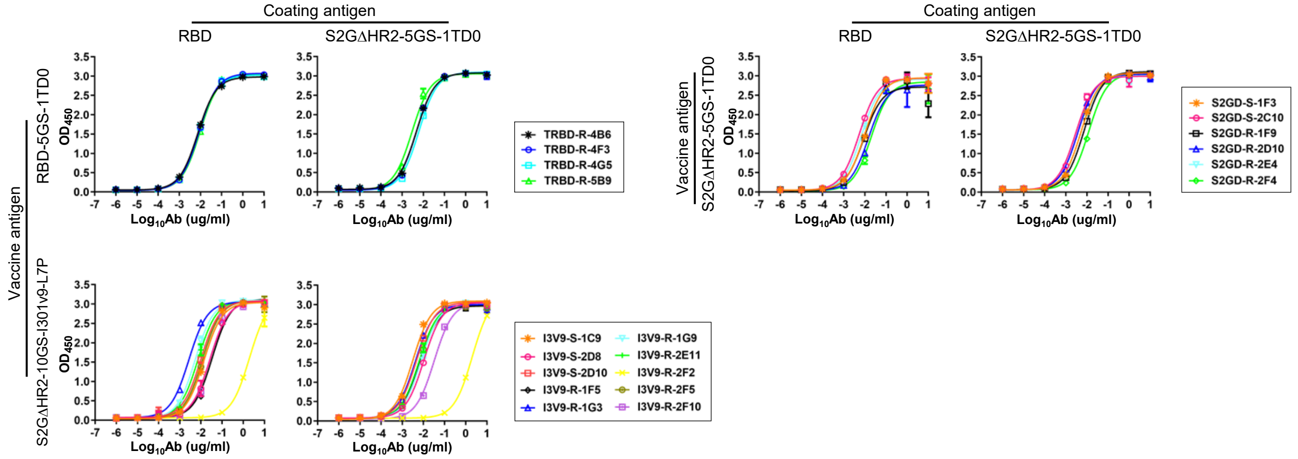

H

**$EC_{50}$ values (μg/ml) of 20 mouse antibodies isolated from mice in three vaccine groups binding to SARS-CoV-2 RBD and spike antigens [a].**

| | TRBD-R-4B6 | TRBD-R-4F3 | TRBD-R-4G5 | TRBD-R-5B9 | S2GD-S-1F3 | S2GD-S-2C10 | S2GD-R-1F9 | S2GD-R-2D10 | S2GD-R-2E4 | S2GD-R-2F4 |
|---|---|---|---|---|---|---|---|---|---|---|
| RBD | 0.007 | 0.009 | 0.008 | 0.010 | 0.010 | 0.005 | 0.009 | 0.015 | 0.007 | 0.020 |
| S2GΔHR2-5GS-1TD0 spike | 0.005 | 0.005 | 0.006 | 0.003 | 0.005 | 0.003 | 0.007 | 0.004 | 0.005 | 0.012 |

| | I3V9-S-1C9 | I3V9-S-2D8 | I3V9-S-2D10 | I3V9-R-1G3 | I3V9-R-1G9 | I3V9-R-1F5 | I3V9-R-2E11 | I3V9-R-2F2 | I3V9-R-2F5 | I3V9-R-2F10 |
|---|---|---|---|---|---|---|---|---|---|---|
| RBD | 0.014 | 0.027 | 0.010 | 0.003 | 0.005 | 0.034 | 0.007 | 1.973 | 0.012 | 0.026 |
| S2GΔHR2-5GS-1TD0 spike | 0.003 | 0.010 | 0.005 | 0.004 | 0.008 | 0.007 | 0.006 | 2.051 | 0.006 | 0.032 |

[a] $EC_{50}$ values were calculated from the besting fitting in GraphPad Prism v9.1.2.

**fig. S3. Single-cell isolation and functional evaluation of monoclonal neutralizing antibodies from mice immunized with the RBD, spike, and SApNP vaccines.** (**A**) SEC profiles of biotinylated Avi-tagged SARS-CoV-2 spike (S2GΔHR2-5GS-foldon-Avi-Biot) and RBD (RBD-5GS-foldon-Avi-Biot) probes from a Superdex 200 Increase 10/300 GL column and a HiLoad Superose 6 16/600 column, respectively. Foldon (PDB: 1RFO) is a trimerization motif used to stabilize the spike and RBD in trimeric conformations and to mask 1TD0-specific B cells (1TD0 is a trimerization motif used in the RBD and spike vaccine constructs). (**B**) Gating strategies used in the antigen-specific single-cell sorting of mouse splenic B cells (Step 1: remove cell debris; Steps 2 and 3: exclude clumped or sticky cells to ensure that only single cells remain; Step 4: remove dead cells; Step 5: identify antigen-specific B cells). Spleen samples from M2 in the RBD-5GS-1TD0 group, M4 in the S2GΔHR2-5GS-1TD0 group, and M2 in the S2GΔHR2-10GS-I3-01v9-L7P group were single-cell sorted using the probes in (A). While only the RBD probe was used to sort B cells from M2 in the RBD-5GS-1TD0 group, both probes were used to sort B cells from the two mice immunized with spike-based vaccines, resulting in a total of 5 sorting experiments. In each sorting experiment, a 96-well plate was used to collect single B cells, which were subjected to antibody cloning and functional validation. Nucleotide and amino acid sequences of monoclonal neutralizing antibodies isolated from (**C**) M2 in the RBD-10GS-1TD0 group (4 antibodies), (**D**) M4 in the S2GΔHR2-5GS-1TD0 group (6 antibodies), and (**E**) M2 in the S2GΔHR2-10GS-I3-01v9-L7P (10 antibodies). Antibodies are named as [Vaccine]-[Probe]-[Antibody index], with heavy and κ-light chains indicated by "H" and "K", respectively. Vaccine can be TRBD, which stands for the RBD-5GS-1TD0 trimer, S2GD, which stands for the S2GΔHR2-5GS-1TD0 spike, and I3V9, which stands for the S2GΔHR2-10GS-I3-01v9-L7P NP. Probe can be S, which stands for spike, and R, which stands for RBD. (**F**) Neutralization curves of 20 mouse monoclonal neutralizing antibodies against SARS-CoV-2-pps that carry spikes of four strains, the wildtype Wuhan-Hu-1 strain and four variants, B.1.1.7, B.1.351, P.1, and $B.1.617_{Rec}$. $IC_{50}$ values were calculated in GraphPad Prism 9.1.2 and are summarized in Fig. 3B. (**G**) ELISA curves of 20 mouse monoclonal neutralizing antibodies binding to the RBD and spike antigens derived from the wildtype Wuhan-Hu-1 strain. All ELISA binding assays were performed in duplicates. (**H**) Summary of $EC_{50}$ values (μg/ml) measured for antibody-antigen binding in (G).

**fig. S4**

A

**Gating strategies in bulk sorting of mouse splenic B cells**

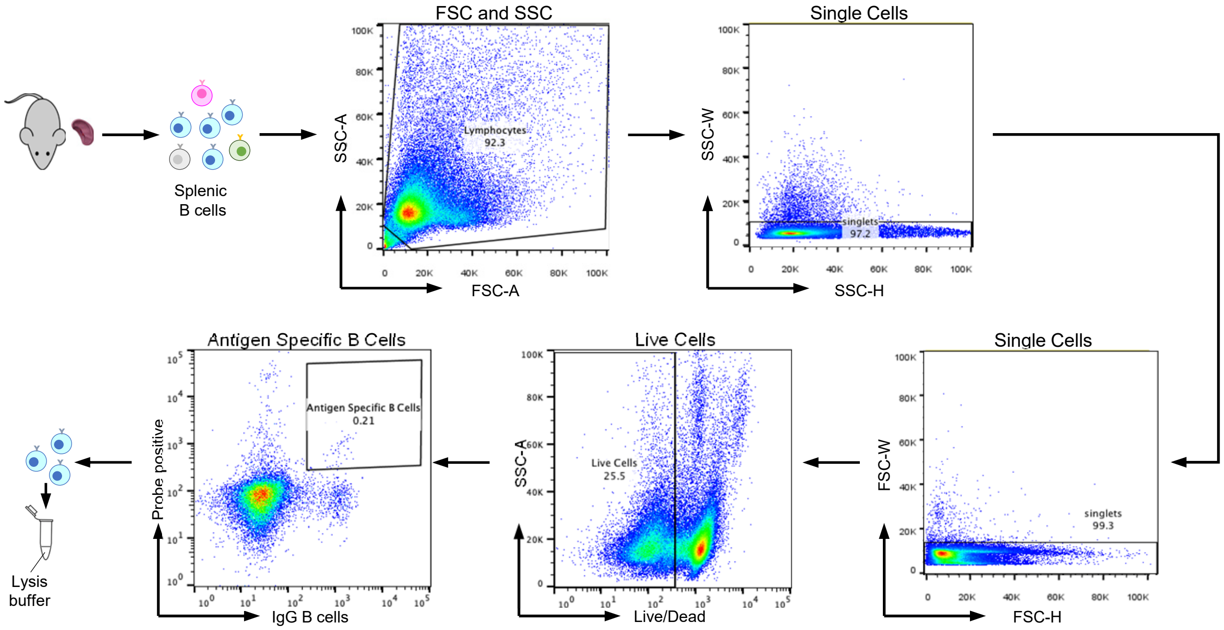


**Antigen-specific sorting of mouse splenic B cells from 3 mice [a]**

| | M2 in the RBD-5GS-1TD0 group | | M4 in the S2GΔHR2-5GS-1TD0 group | | M2 in the S2GΔHR2-10GS-I3-01v9-L7P group | |
|---|---|---|---|---|---|---|
| Probe | Sorted B cells | %Splenic B cells | Sorted B cells | %Splenic B cells | Sorted B cells | %Splenic B cells |
| S2GΔHR2-5GS-foldon-Avi-Biot | 1481 | 0.36370 | 1550 | 0.21626 | 1407 | 0.25887 |
| RBD-5GS-foldon-Avi-Biot | 1565 | 0.23684 | 1550 | 0.27332 | 1402 | 0.37566 |

[a] Spleen samples from three mice in the previous study (Ref 41) were processed for bulk sorting of antigen-specific B cells to facilitate deep sequencing analysis. Two SARS-CoV-2 antigen probes, S2GΔHR2-5GS-foldon-Avi-Biot and RBD-Avi-Biot, were used in the bulk sorting.

B

**Next-generation sequencing (NGS) analysis of 3 mice immunized with RBD, spike, and nanoparticle vaccines [a]**

| Vaccine antigen | Probe [b] | $N_{Raw}$ | $N_{V\text{-}assign}$ | $N_{V\text{-}align\ (250bp)}$ | Chain | $N_{Chain}$ | $N_{Step5}$ | <Length> | $N_{Usable}$ |
|---|---|---|---|---|---|---|---|---|---|
| M2 in the RBD-5GS-1TD0 group | Spike | 2,067,177 | 842,447 | 335,620 | H | 150,323 | 149,768 | 361.0 | 149,767 |
| | | | | | K | 185,297 | 184,846 | 331.5 | 184,845 |
| | RBD | 1,258,194 | 831,049 | 470,912 | H | 208,287 | 207,731 | 359.9 | 207,728 |
| | | | | | K | 262,625 | 262,030 | 335.5 | 262,028 |
| M4 in the S2GΔHR2-5GS-1TD0 group | Spike | 1,431,024 | 771,147 | 376,522 | H | 111,699 | 111,440 | 357.7 | 111,438 |
| | | | | | K | 264,823 | 264,345 | 339.6 | 264,344 |
| | RBD | 1,380,740 | 907,658 | 380,881 | H | 154,863 | 154,571 | 355.4 | 154,570 |
| | | | | | K | 226,017 | 224,892 | 337.9 | 224,892 |
| M2 in the S2GΔHR2-10GS-I3-01v9-L7P group | Spike | 3,952,238 | 1,873,397 | 720,897 | H | 411,322 | 410,074 | 360.2 | 410,068 |
| | | | | | K | 309,575 | 308,842 | 330.3 | 308,842 |
| | RBD | 1,752,190 | 954,410 | 389,625 | H | 200,507 | 197,000 | 361.0 | 196,995 |
| | | | | | K | 189,118 | 188,800 | 329.9 | 188,798 |

[a] Listed items include the vaccine antigen, mouse sample ID, number of raw reads from Ion S5 sequencing, number of reads that can be assigned to a VH/VK gene with an E-value of $10^{-3}$ or lower, number of reads that can be aligned to a VH/VK gene with 250bp or longer, number of VH/VK chains, number of VH/VK chains at the last step (5) of pipeline processing, average read length, and number of usable chains. Of note, to determine usable chains, the 250bp V-gene alignment filter was applied again to remove any problematic sequences detected during the full pipeline processing.

[b] Two SARS-CoV-2 probes were used in antigen-specific bulk sorting of mouse splenic B cells. Spike stands for S2GΔHR2-5GS-foldon-Avi-Biot; RBD stands for RBD-5GS-foldon-Avi-Biot.

### fig. S4

C **Spike/RBD-specific bulk sorting of splenic B cells from three mice**

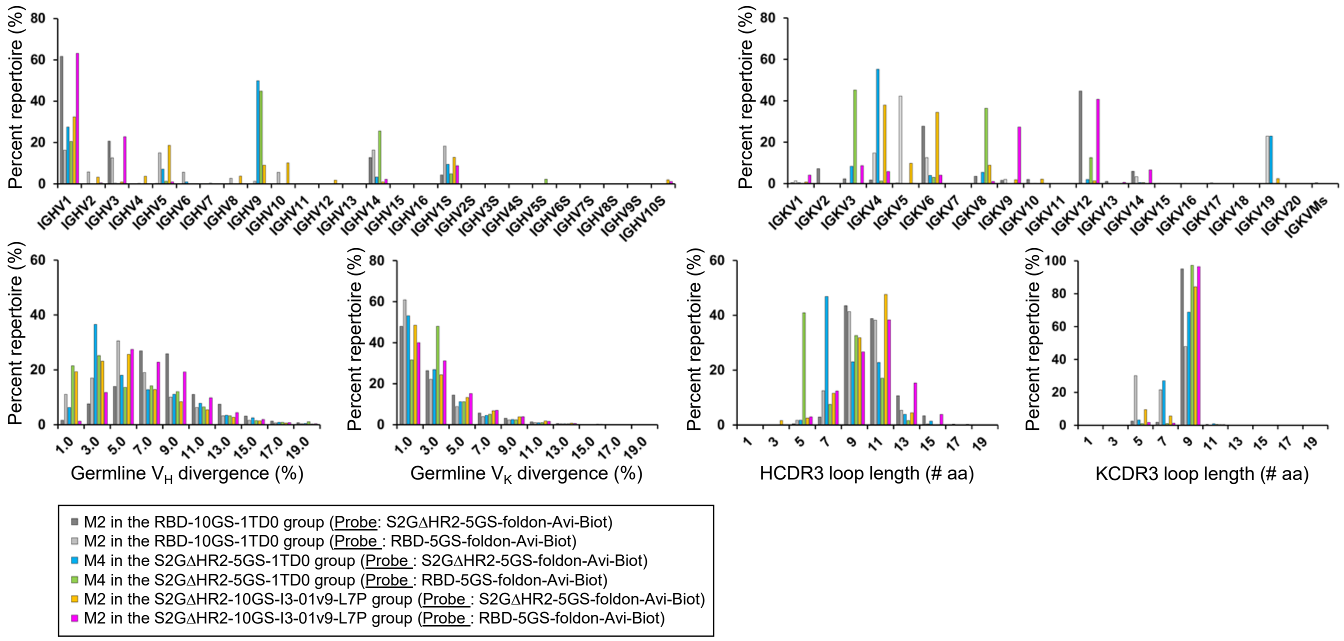


D **Tracing RBD-5GS-1TD0 trimer-elicited NAbs in spike-sorted B cell populations**

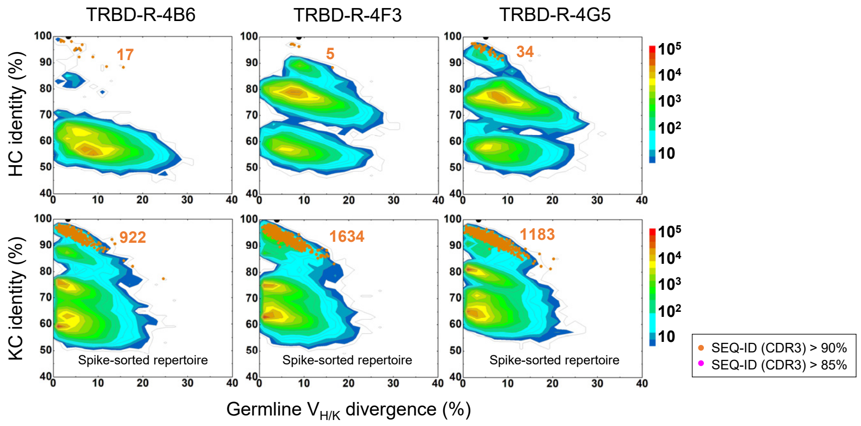


E **Tracing S2GΔHR2-5GS-1TD0 spike-elicited NAbs in spike/RBD-sorted B cell populations**

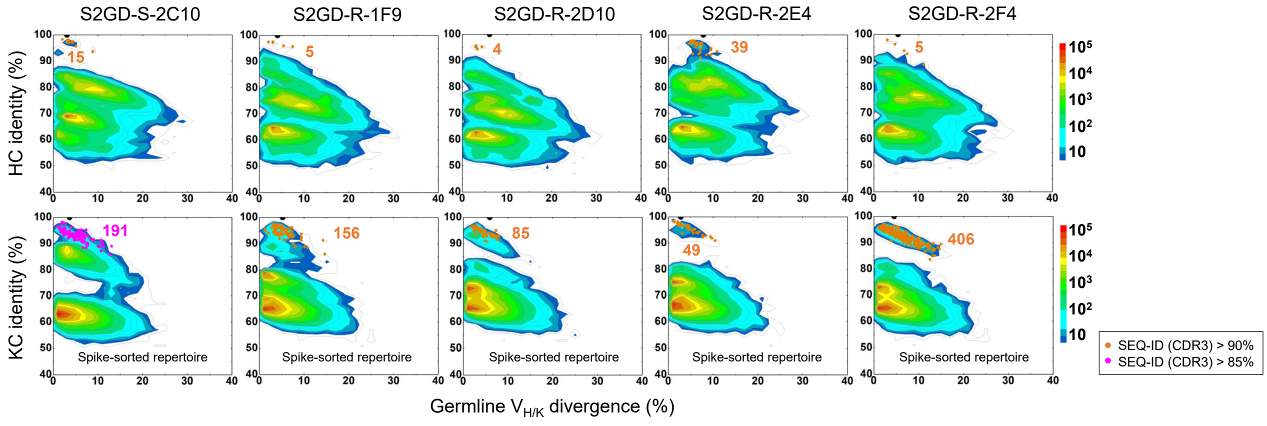

### fig. S4

#### E (continued)

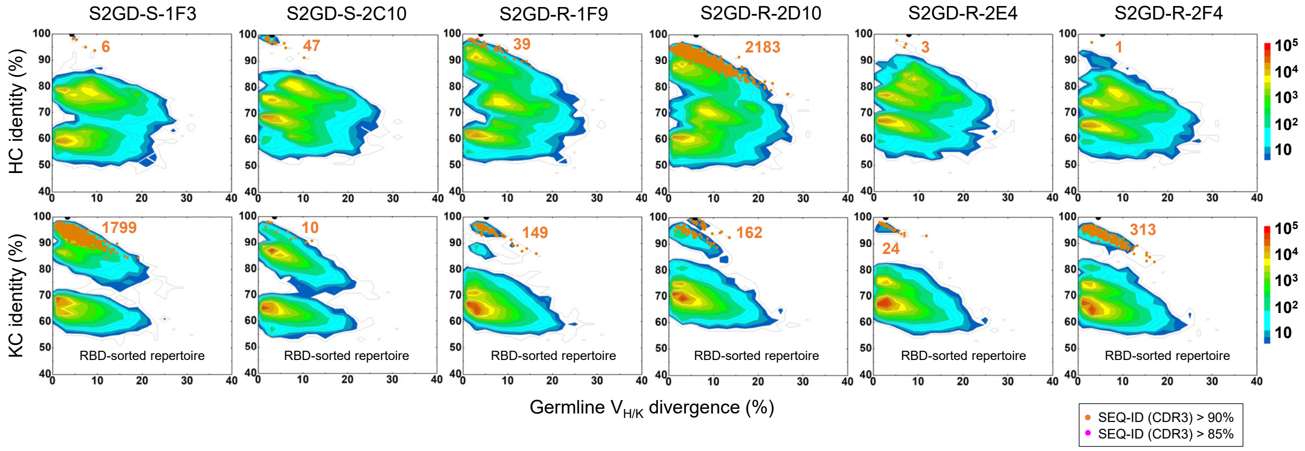


#### F Tracing S2GΔHR2-10GS-I3-01v9-L7P NP-elicited NAbs in spike/RBD-sorted B cell populations

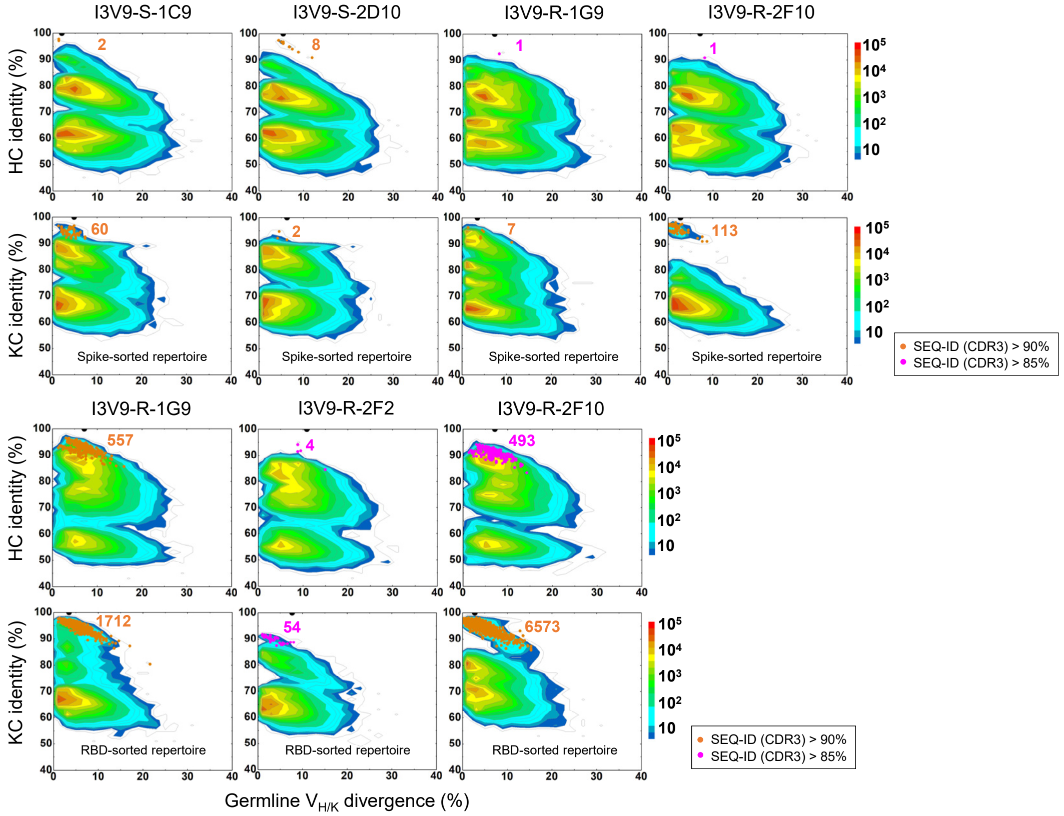

### fig. S4

**fig. S4. Unbiased repertoire analysis of bulk-sorted SARS-CoV-2 antigen-specific mouse splenic B cells and tracing of mouse neutralizing antibodies in the NGS-derived repertoires.** Two SARS-CoV-2 antigen probes in fig. S3A were used to sort mouse splenic B cells. (**A**) Bulk B-cell sorting experiment. Top: gating strategies used in the antigen-specific sorting of mouse splenic B cells (Step 1: remove cell debris; Steps 2 and 3: exclude clumped or sticky cells to ensure that only single cells remain; Step 4: remove dead cells; Step 5: identify antigen-specific B cells). Bottom: Summary of SARS-CoV-2 antigen-specific bulk sorting of mouse splenic B cells from three mice. (**B**) Antibodyomics pipeline analysis of repertoire NGS data obtained for spike and RBD-specific mouse splenic B cells from three mice. (**C**) B-cell repertoire profiles are shown for three mice immunized with RBD-5GS-1TD0, S2GΔHR2-5GS-1TD0, and S2GΔHR2-10GS-I3-01v9-L7P. Top: VH gene usage (left) and VK gene usage (right); Bottom: germline VH/VK divergence (left) and CDRH/K3 loop length (right). **(D)** Divergence-identity analysis of NAbs in the context of RBD-sorted B-cell repertoires for M2 in the RBD-5GS-1TD0 group. (**E**) Divergence-identity analysis of NAbs in the context of spike and RBD-sorted B-cell repertoires for M4 in the S2GΔHR2-5GS-1TD0 group. (**F**) Divergence-identity analysis of NAbs in the context of spike and RBD-sorted B-cell repertoires for M2 in the S2GΔHR2-10GS-I3-01v9-L7P group. HC and KC sequences are plotted as a function of sequence identity to a given template NAb and sequence divergence from putative germline V genes. Color coding indicates sequence density. On the 2D plots, template NAbs are shown as black dots, whereas somatic variants that were identified based on the germline V gene and the CDR3 identity of 85/90% or greater are shown as orange/magenta dots, with the number of sequences labeled accordingly. In (D)-(F), the 2D plots are only shown for NAbs for which both HC and KC somatic variants could be found in the NGS repertoires.

**fig. S5**

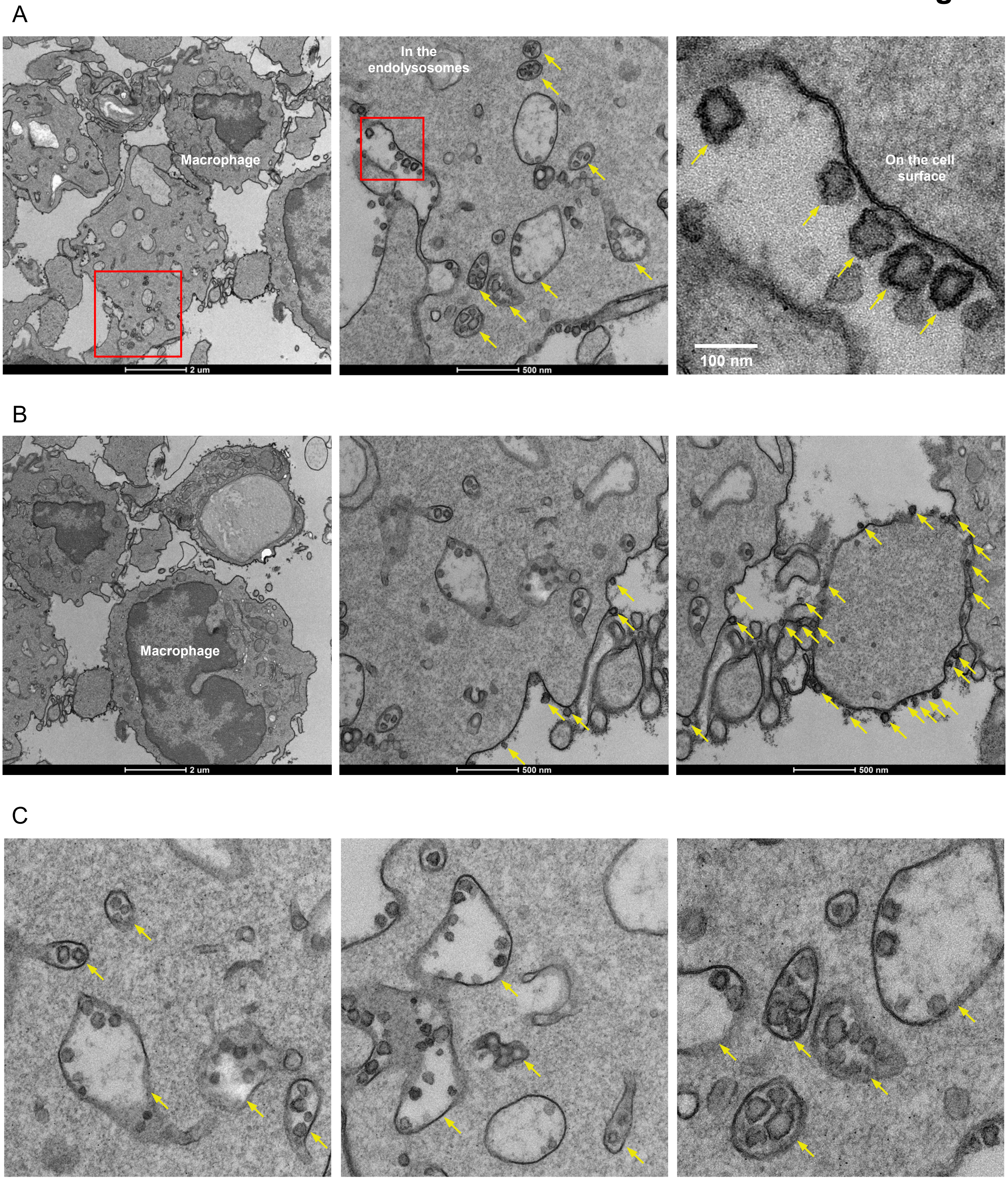


**fig. S5. SARS-CoV-2 spike-presenting I3-01v9 SApNP interaction with macrophages in a lymph node.** (**A**) S2GΔHR2-presenting I3-01v9 SApNPs are sequestered by macrophages in the medullary cord zone of a lymph node after a single-dose injection (50 μg). S2GΔHR2-presenting I3-01v9 SApNPs (**B**) aligned on macrophage surface or (**C**) sequestered inside the endolysosomes of a macrophage. S2GΔHR2-presenting I3-01v9 SApNPs are pointed by yellow arrows.

fig. S6

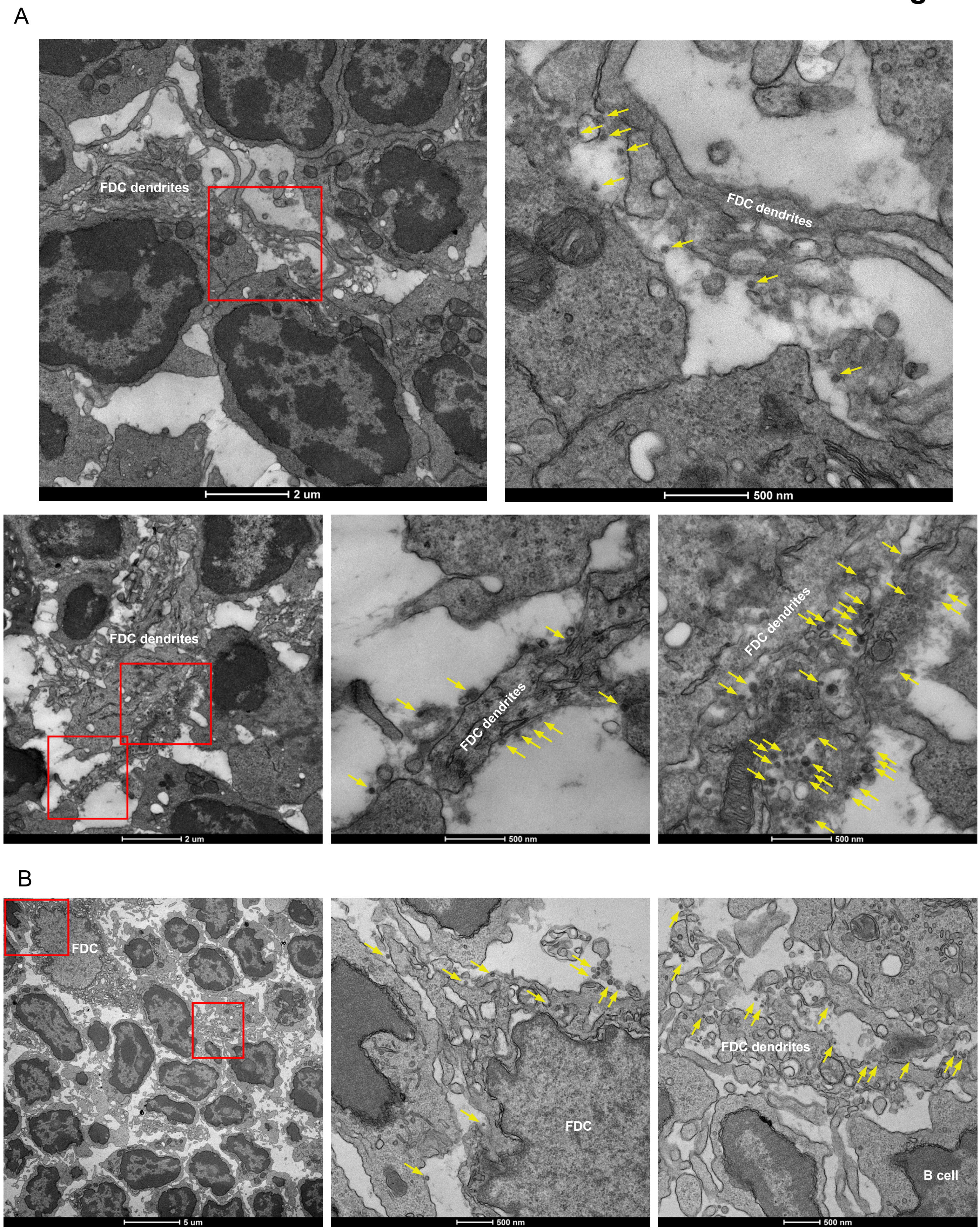

A
FDC dendrites
2 um
FDC dendrites
500 nm
FDC dendrites
2 um
FDC dendrites
500 nm
FDC dendrites
500 nm
B
FDC
5 um
FDC
500 nm
FDC dendrites
B cell
500 nm

fig. S6

C

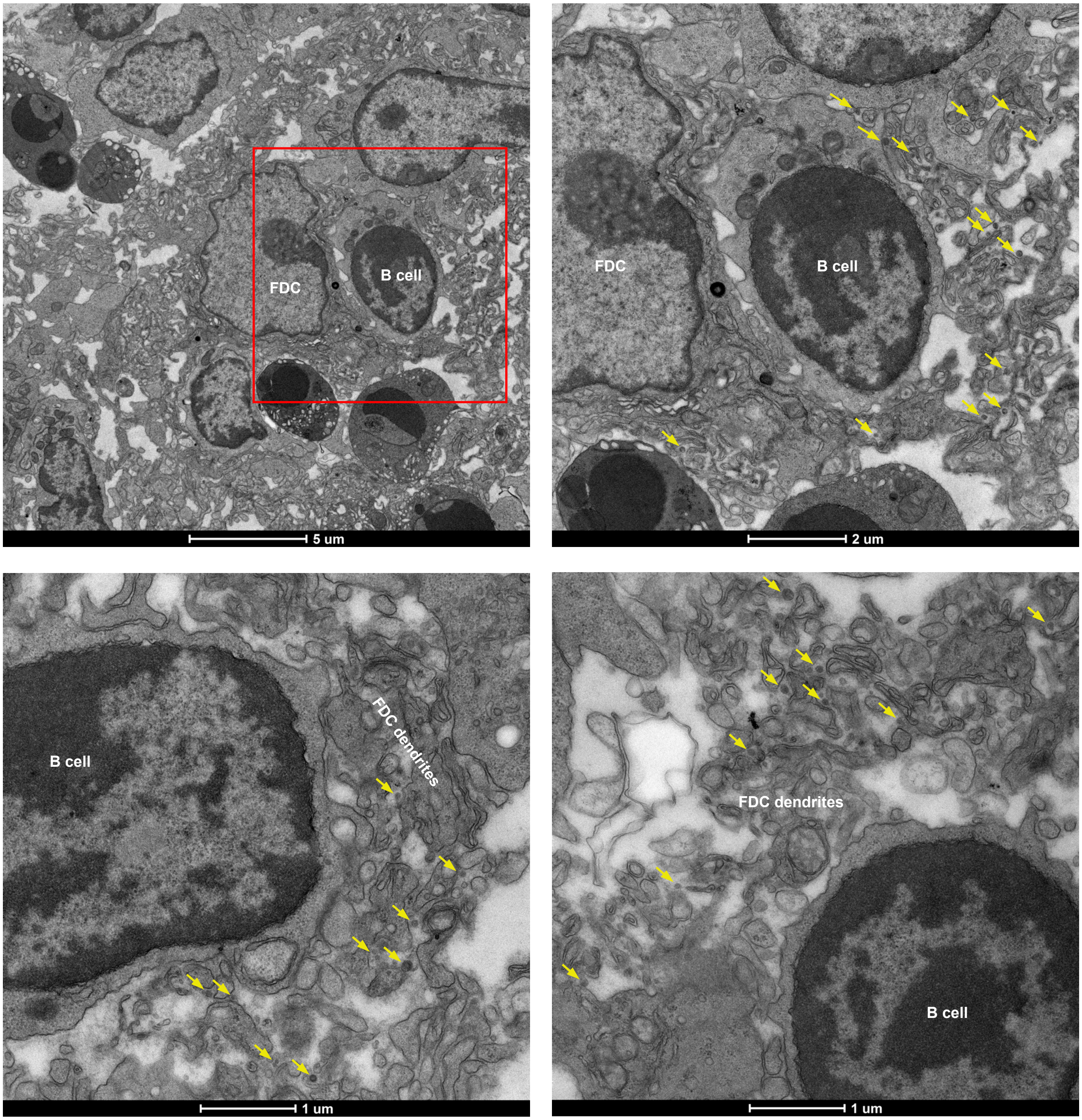

FDC
B cell
5 um
FDC
B cell
2 um
B cell
FDC dendrites
1 um
FDC dendrites
B cell
1 um

### fig. S6

D

**I3-01 nanoparticles injected at 2h**

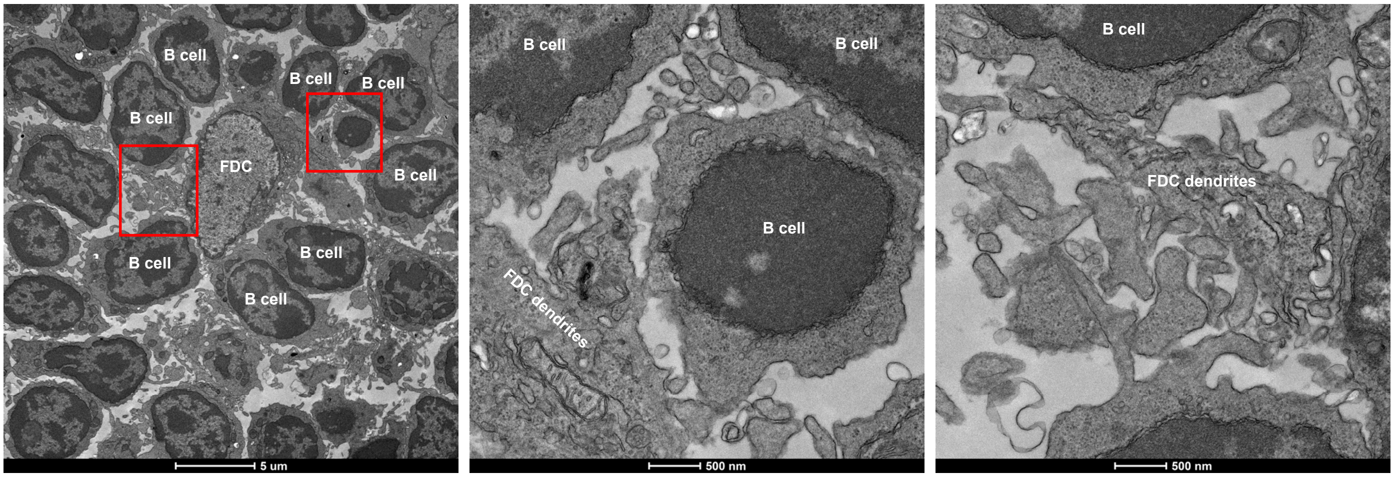


E

**Lymph node from naïve mouse**

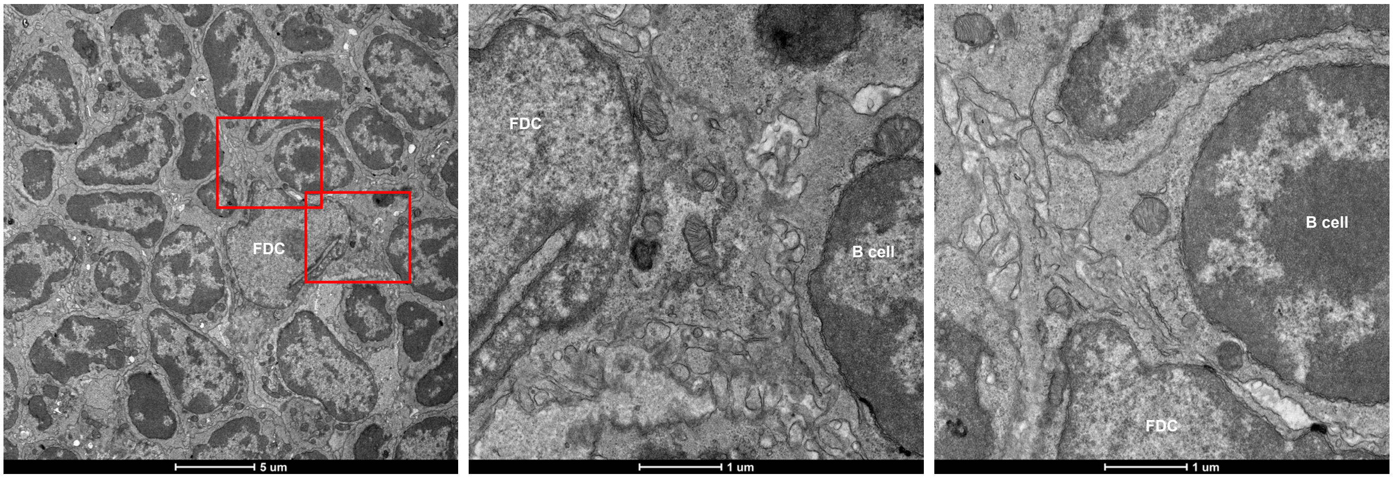


**fig. S6. TEM images of SARS-CoV-2 spike-presenting I3-01v9 SApNP interaction with FDCs in a lymph node.** S2GΔHR2-presenting I3-01v9 SApNPs are aligned on FDC dendrites (**A**) at 12 h after a single-dose injection (50 μg), (**B**) at 48 h after a single-dose injection (50 μg), and (**C**) at 12 h after the boost injection (50 μg). S2GΔHR2-presenting I3-01v9 SApNPs are pointed by yellow arrows. (**D**) S2GΔHR2-presenting I3-01v9 SApNPs were barely observed at 2 h after a single-dose injection (50 μg). (**E**) Lymph node from a naïve, unimmunized mouse.

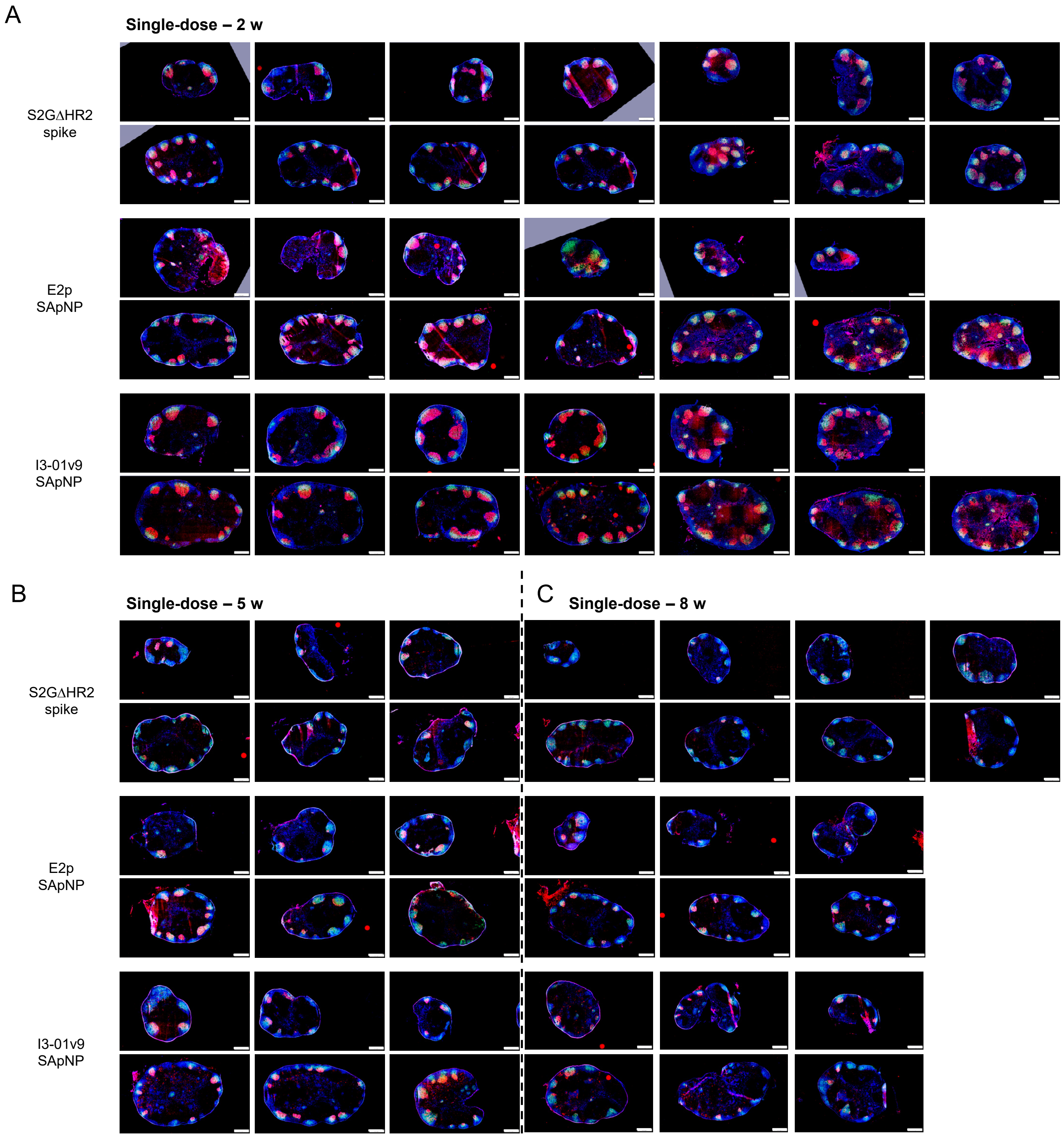
A
Single-dose – 2 w
S2GΔHR2 spike
E2p SApNP
I3-01v9 SApNP
B
Single-dose – 5 w
S2GΔHR2 spike
E2p SApNP
I3-01v9 SApNP
C
Single-dose – 8 w

fig. S7

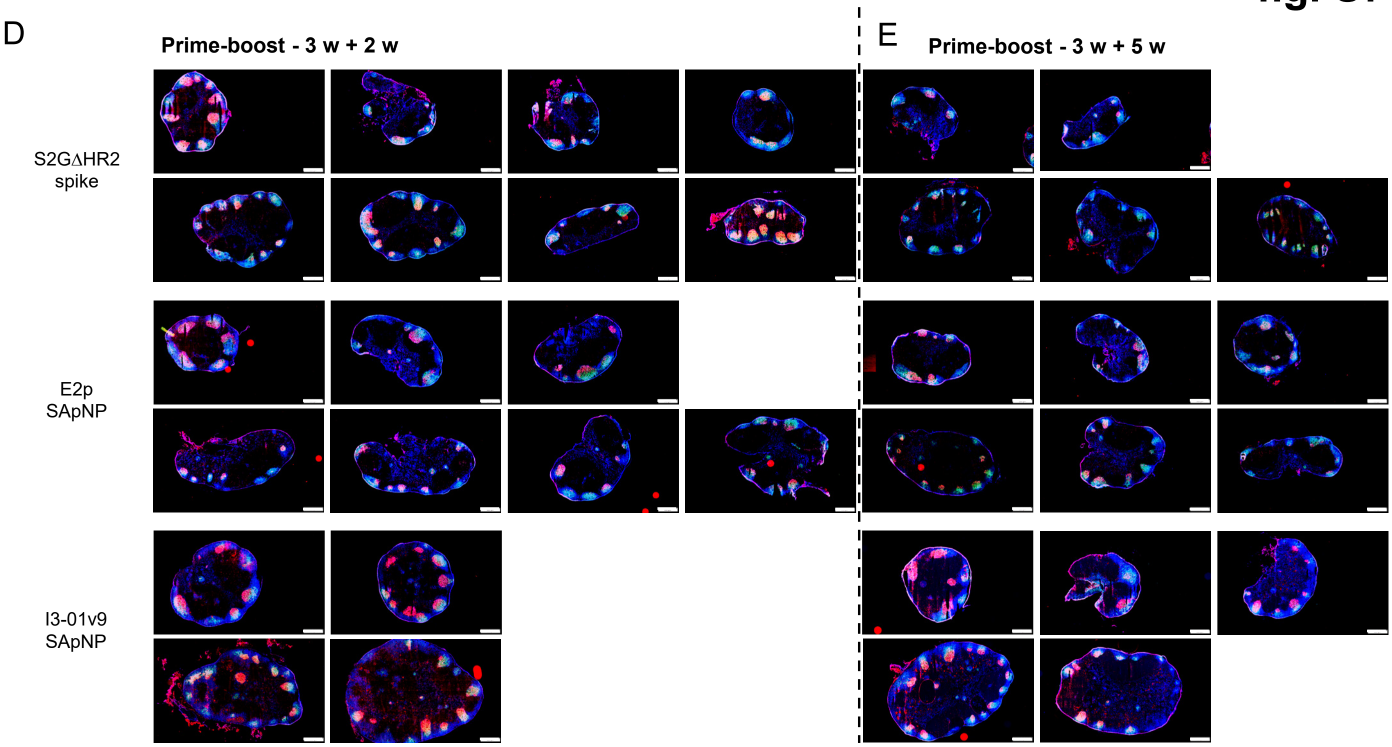


**fig. S7. Immunohistological analysis of SARS-CoV-2 spike/spike-presenting SApNP vaccine-induced GCs.** Images of germinal centers at (**A**) week 2, (**B**) week 5, and (**C**) week 8 after a single-dose injection of S2GΔHR2 spike and S2GΔHR2-presenting E2p and I3-01v9 SApNP vaccines (10 μg per injection, totaling 40 μg per mouse), with a scale bar of 500 μm for each image. Images of germinal centers at (**D**) week 2 and (**E**) week 5 after prime-boost injections.

A

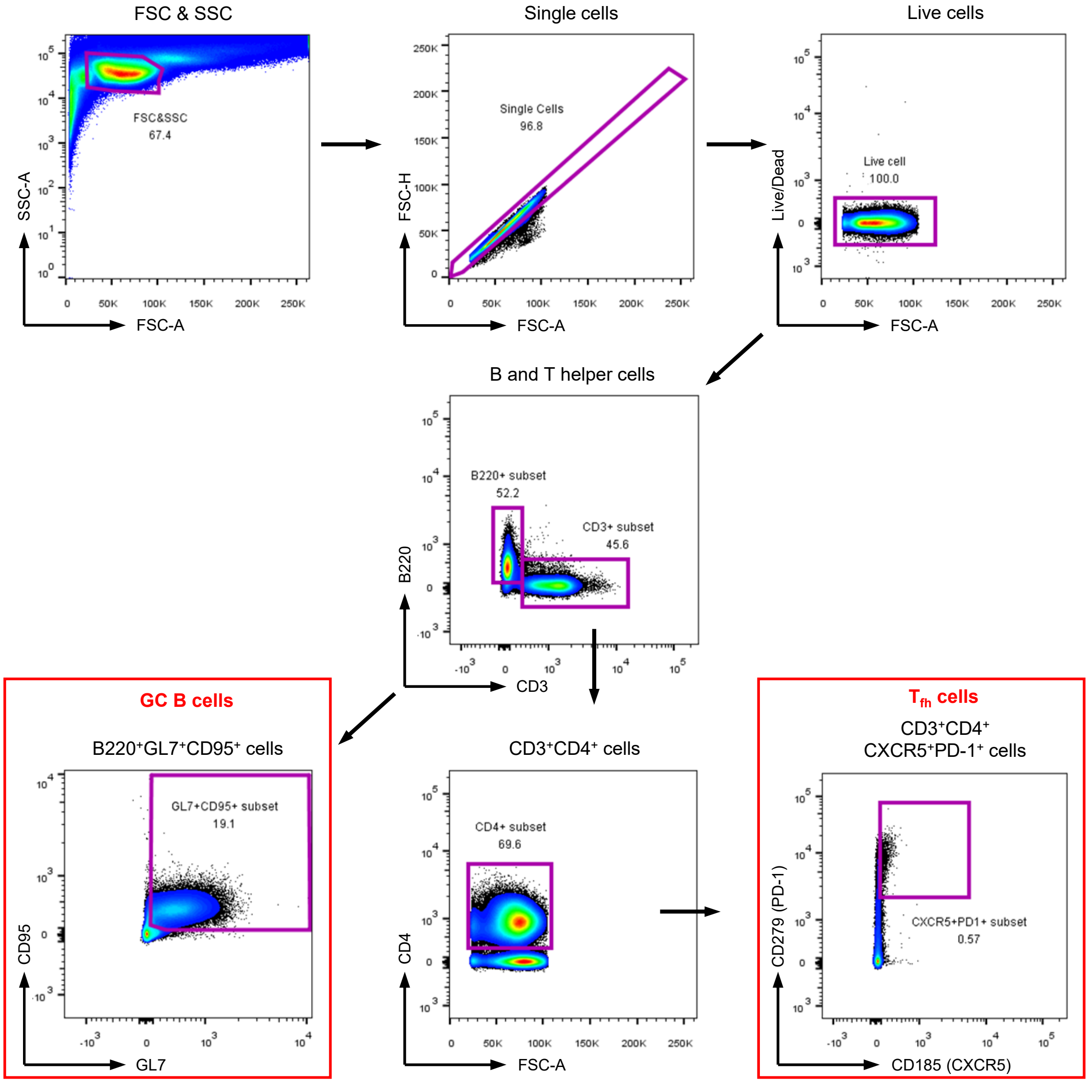
FSC & SSC
FSC&SSC
67.4
SSC-A
FSC-A
Single cells
Single Cells
96.8
FSC-H
FSC-A
Live cells
Live cell
100.0
Live/Dead
FSC-A
B and T helper cells
B220+ subset
52.2
CD3+ subset
45.6
B220
CD3
GC B cells
B220⁺GL7⁺CD95⁺ cells
GL7+CD95+ subset
19.1
CD95
GL7
CD3⁺CD4⁺ cells
CD4+ subset
69.6
CD4
FSC-A
Tfh cells
CD3⁺CD4⁺
CXCR5⁺PD-1⁺ cells
CXCR5+PD1+ subset
0.57
CD279 (PD-1)
CD185 (CXCR5)

fig. S8

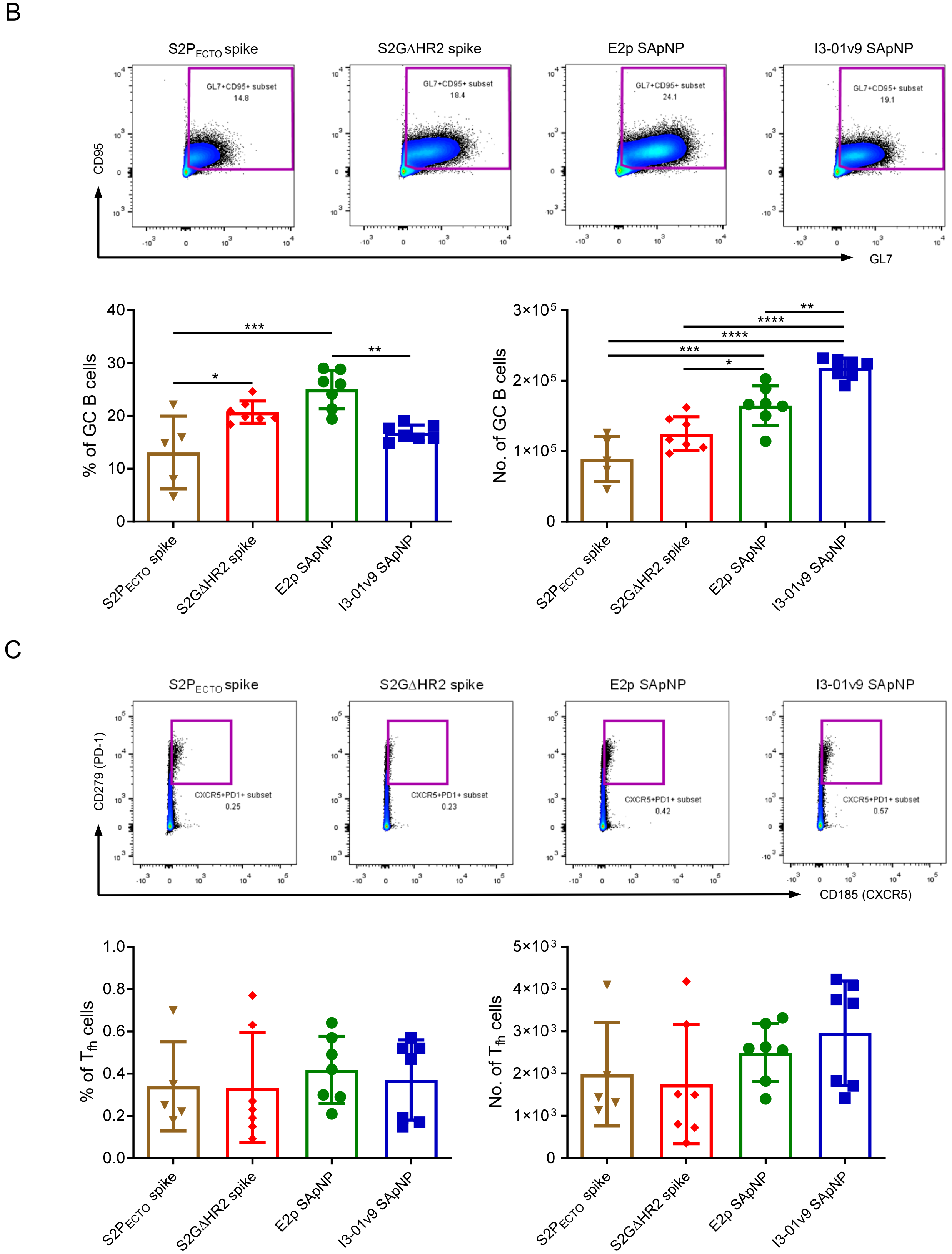


**fig. S8. Flow cytometry analysis of SARS-CoV-2 spike/spike-presenting SApNP vaccine-induced GCs.** (**A**) Gating strategy for analyzing germinal center reactions using flow cytometry. Quantification of germinal center reactions, including (**B**) GC B cells and (**C**) T follicular helper cells at week 2 after a single-dose injection of $S2P_{ECTO}$, S2GΔHR2, and S2GΔHR2-presenting E2p and I3-01v9 SApNP vaccines (10 μg per injection, totaling 40 μg per mouse). Data points are presented as mean ± SD. The *P* values were determined by one-way ANOVA followed by Tukey's multiple comparisons *post hoc* test for each timepoint. **p* < 0.05, ***p* < 0.01, ****p* < 0.001, *****p* < 0.0001.

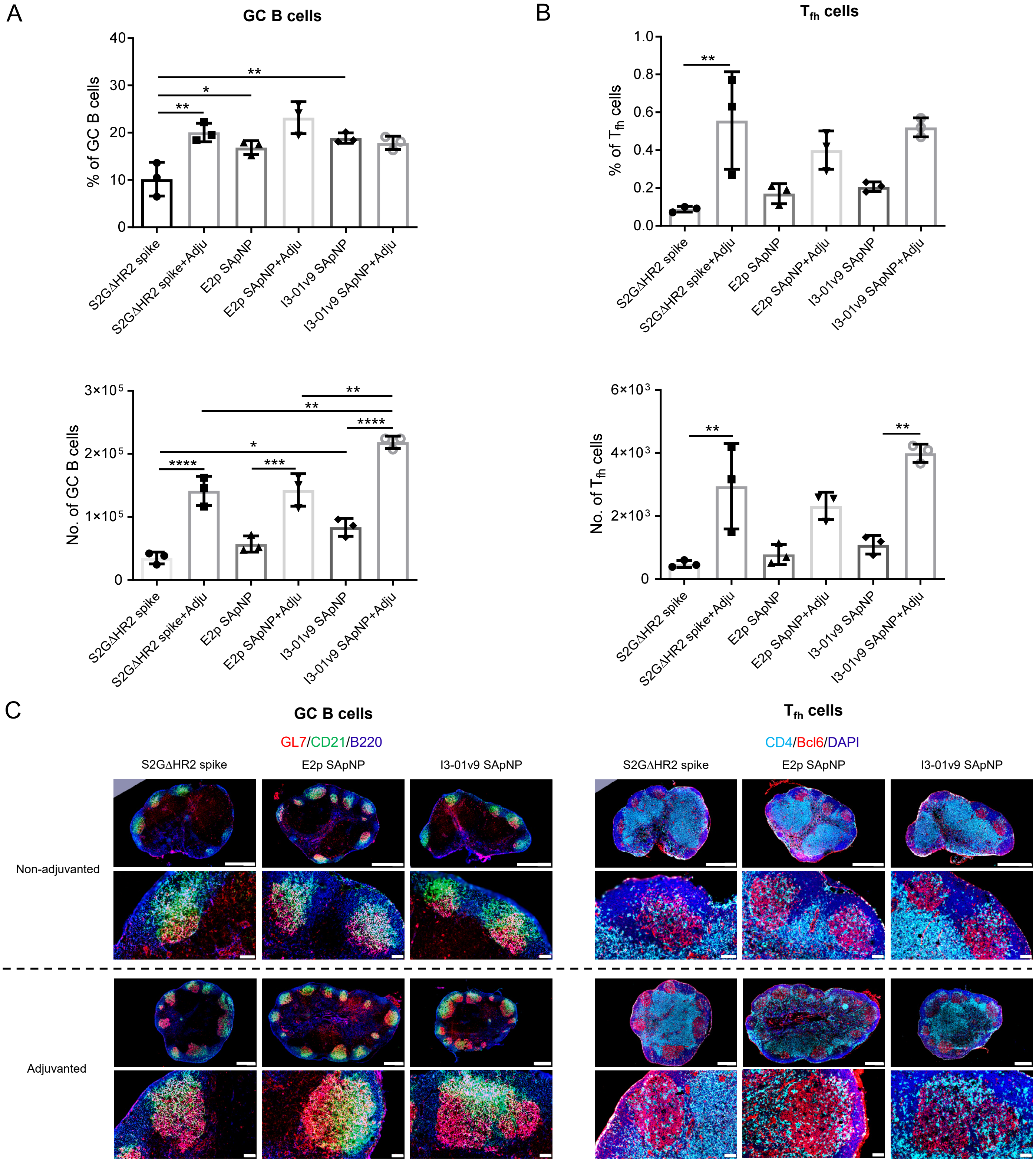


**fig. S9. Adjuvant effect on SARS-CoV-2 spike/spike-presenting SApNP vaccine-induced GCs.** Quantification of germinal center reactions, including (**A**) GC B cells and (**B**) T follicular helper cells at week 2 after a single-dose injection of S2P$_{ECTO}$, S2GΔHR2, and S2GΔHR2-presenting E2p and I3-01v9 SApNP vaccines with/without adjuvants using flow cytometry (10 μg per injection, totaling 40 μg per mouse). (**C**) Immunohistology of germinal centers at week 2 after a single-dose injection with/without adjuvants. Scale bars of 500 μm and 50 μm are shown for a complete lymph node and an enlarged image of a follicle, respectively. Data points are presented as mean ± SD. The *P* values were determined by one-way ANOVA followed by Tukey's multiple comparisons *post hoc* test for each timepoint. *$p < 0.05$, **$p < 0.01$, ***$p < 0.001$, ****$p < 0.0001$.

### fig. S10

A

**Gating strategies in bulk B cell sorting**

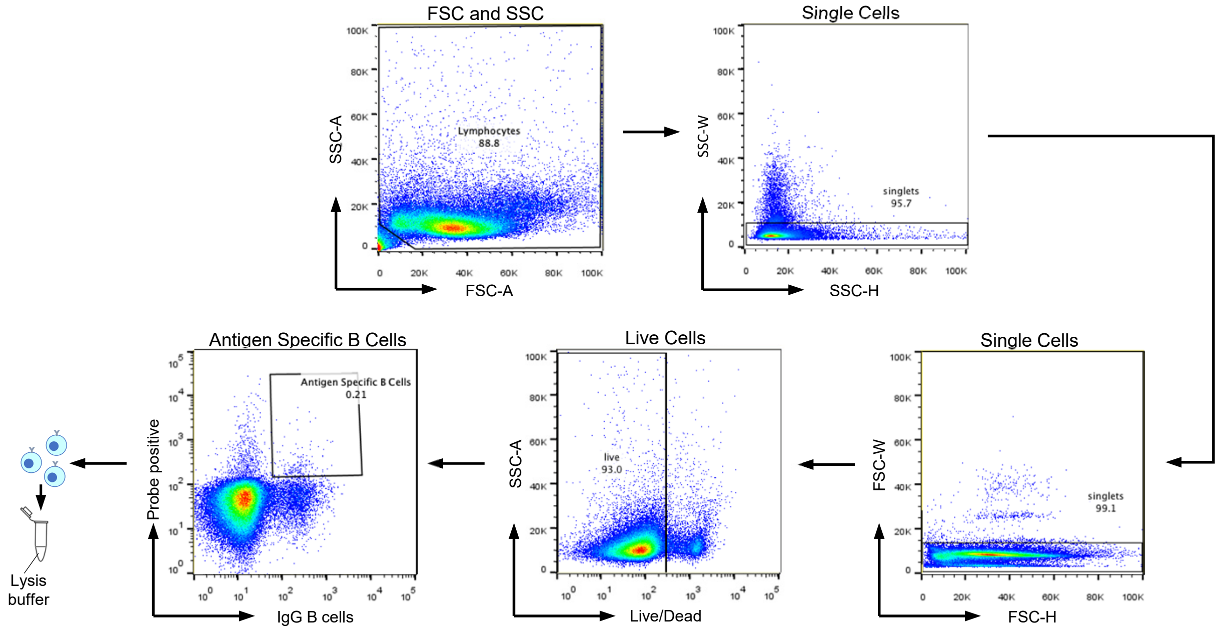


**SARS-CoV-2 S2GΔHR2-specific sorting of mouse lymph node (LN) B cells from 3 low-dosage vaccine groups [a]**

| S2GΔHR2-5GS-1TD0 | | | S2GΔHR2-5GS-E2p-LD4-PADRE | | | S2GΔHR2-5GS-I3-01v9-LD7-PADRE | | |
|---|---|---|---|---|---|---|---|---|
| Mouse | Sorted B cells | %LN B cells | Mouse | Sorted B cells | %LN B cells | Mouse | Sorted B cells | %LN B cells |
| G5-1 | 487 | 0.07656 | G6-1 | 670 | 0.08735 | G7-1 | 1425 | 0.23039 |
| G5-2 | 567 | 0.10521 | G6-2 | 381 | 0.07638 | G7-2 | 1196 | 0.24353 |
| G5-3 | 358 | 0.07592 | G6-3 | 314 | 0.05040 | G7-3 | 1893 | 0.39877 |
| G5-4 | 552 | 0.08819 | G6-4 | 522 | 0.09089 | G7-4 | 1164 | 0.23013 |
| G5-5 | 442 | 0.07488 | G6-5 | 297 | 0.05094 | G7-5 | 1377 | 0.08676 |

[a] In this study, each mouse was immunized at w0 and w3 with a dosage of 3.3μg nanoparticle protein mixed with respective adjuvants. LN samples at w5 were processed for bulk sorting of S2GΔHR2-specific B cells to facilitate deep sequencing analysis.

B

**Next-generation sequencing (NGS) analysis of 3 low-dosage SARS-CoV-2 vaccine groups [a]**

| Vaccine antigen | Mouse | $N_{Raw}$ | $N_{V\text{-}assign}$ | $N_{V\text{-}align\ (250bp)}$ | Chain | $N_{Chain}$ | $N_{Step5}$ | <Length> | $N_{Usable}$ |
|---|---|---|---|---|---|---|---|---|---|
| S2GΔHR2-5GS-1TD0 (Spike) | G5-1 | 1,596,202 | 441,528 | 99,119 | H | 49,456 | 49,347 | 361.3 | 49,347 |
| | | | | | K | 49,663 | 49,480 | 337.5 | 49,480 |
| | G5-2 | 846,960 | 393,825 | 208,528 | H | 47,402 | 47,286 | 364.0 | 47,283 |
| | | | | | K | 161,126 | 160,748 | 330.8 | 160,747 |
| | G5-3 | 843,754 | 289,986 | 71,301 | H | 27,115 | 270,12 | 362.0 | 27,010 |
| | | | | | K | 44,186 | 44,038 | 333.0 | 44,038 |
| | G5-4 [b] | 1,030,253 | 270,732 | 34,713 | H | 34,674 | 34,584 | 355.9 | 34,583 |
| | | | | | K [c] | 39 | 39 | 331.0 | 39 |
| | G5-5 | 1,018,511 | 362,904 | 75,634 | H | 37,696 | 37,544 | 361.2 | 37,542 |
| | | | | | K | 37,938 | 37,851 | 336.3 | 37,851 |
| S2GΔHR2-5GS-E2p-LD4-PADRE (NP) | G6-1 | 844,432 | 442,045 | 150,263 | H | 118,683 | 118,154 | 359.6 | 118,152 |
| | | | | | K | 31,580 | 31,338 | 340.6 | 31,338 |
| | G6-2 | 743,730 | 391,866 | 118,081 | H | 31,763 | 31,681 | 359.5 | 31,677 |
| | | | | | K | 86,318 | 86,150 | 334.6 | 86,149 |
| | G6-3 | 847,416 | 360,187 | 100,405 | H | 34,702 | 34,588 | 361.8 | 34,586 |
| | | | | | K | 65,703 | 65,385 | 343.5 | 65,385 |
| | G6-4 | 804,742 | 325,463 | 148,519 | H | 57,260 | 55,248 | 358.1 | 55,245 |
| | | | | | K | 91,259 | 90,934 | 332.2 | 90,934 |
| | G6-5 | 1,226,643 | 388,694 | 148,556 | H | 45,000 | 44,835 | 360.3 | 44,833 |
| | | | | | K | 103,556 | 103,386 | 328.5 | 103,385 |
| S2GΔHR2-10GS-I3-01v9-LD7-PADRE (NP) | G7-1 | 698,174 | 561,918 | 196,631 | H | 84,704 | 84,373 | 359.5 | 84,371 |
| | | | | | K | 111,927 | 111,777 | 331.6 | 111,776 |
| | G7-2 | 685,753 | 587,525 | 226,293 | H | 84,558 | 84,259 | 359.0 | 84,258 |
| | | | | | K | 141,735 | 141,506 | 332.1 | 141,505 |
| | G7-3 | 772,549 | 645,323 | 233,227 | H | 85,301 | 84,944 | 359.1 | 84,944 |
| | | | | | K | 147,926 | 147,648 | 332.4 | 147,648 |
| | G7-4 | 887,414 | 388,120 | 126,844 | H | 59,480 | 59,271 | 358.8 | 59,268 |
| | | | | | K | 67,364 | 67,232 | 330.4 | 67,232 |
| | G7-5 | 537,471 | 283,727 | 103,400 | H | 37,814 | 37,329 | 358.8 | 37,329 |
| | | | | | K | 65,586 | 65,438 | 345.3 | 65,437 |

[a] Listed items include the vaccine antigen, mouse sample ID, number of raw reads from Ion S5 sequencing, number of reads that can be assigned to a VH/VK gene with an E-value of $10^{-3}$ or lower, number of reads that can be aligned to a VH/VK gene with 250bp or longer, number of VH/VK chains, number of VH/VK chains at the last step (5) of pipeline processing, average read length, and number of usable chains. Of note, to determine usable chains, the 250bp V-gene alignment filter was applied again to remove any problematic sequences detected during the full pipeline processing.

[b] Due to the difficulty in κ-light chain library preparation, only 39 reads were obtained for M4 sample in the S2GΔHR2-5GS-1TD0 group. The results from pipeline processing are provided here only for the reason of completeness.

### fig. S10

C **S2GΔHR2-5GS-1TD0 spike group**

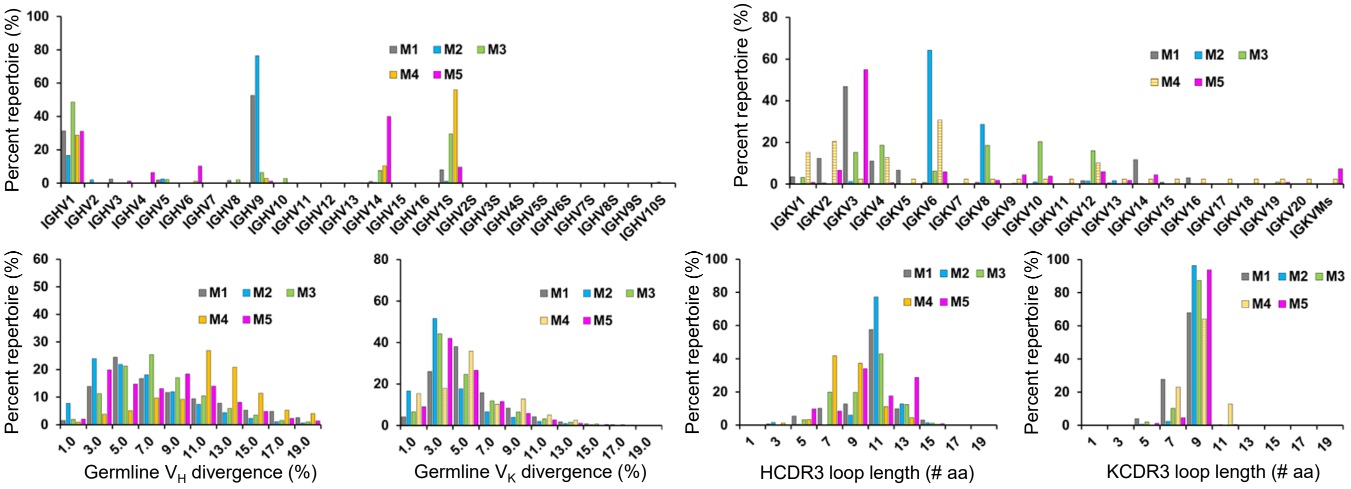


D **S2GΔHR2-5GS-E2p-LD4-PADRE NP group**

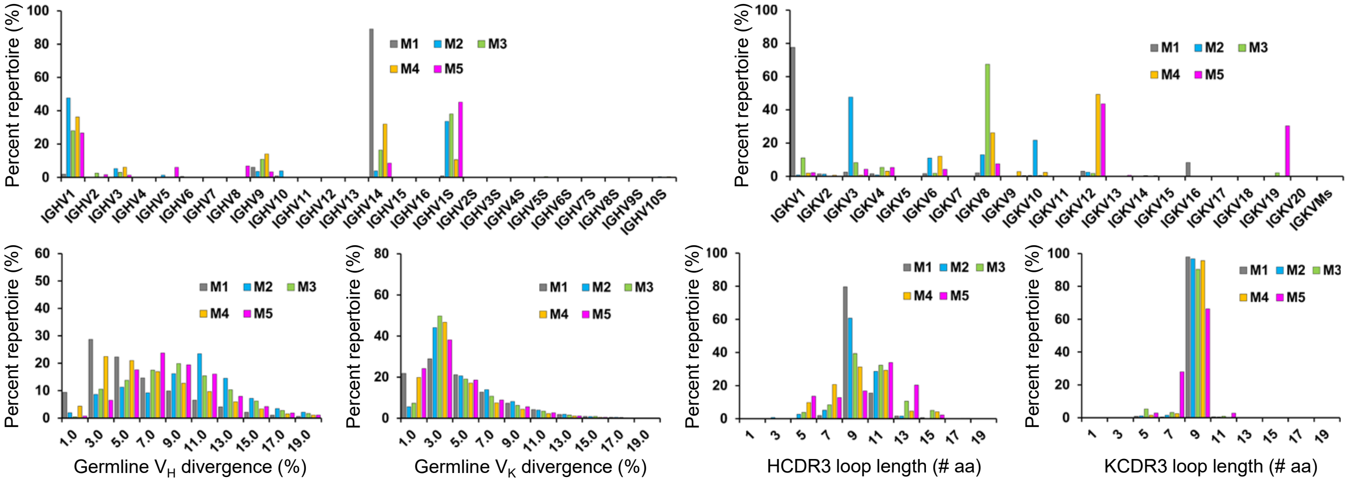


E **S2GΔHR2-10GS-I3-01v9-LD7-PADRE NP group**

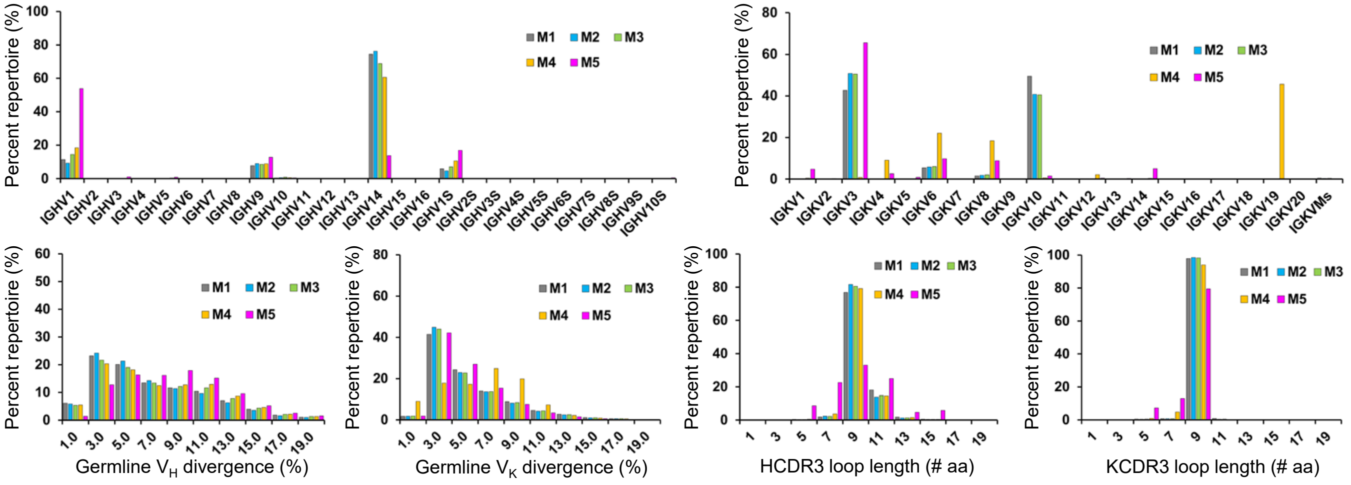

### fig. S10

F

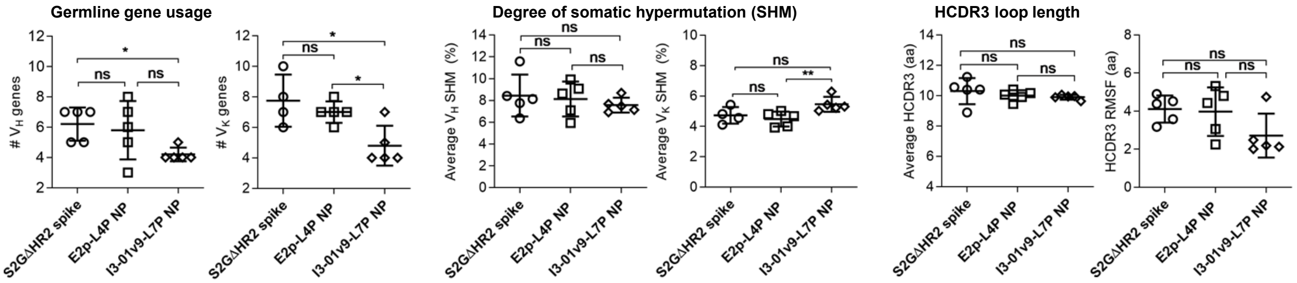


**fig. S10. NGS analysis of SARS-CoV-2 spike-specific lymph node (LN) B cells from mice immunized with the spike and SApNP vaccines.** In this study, three groups of mice were immunized with S2GΔHR2-5GS-1TD0/AddaVax, S2GΔHR2-5GS-E2p-L4P/AddaVax, S2GΔHR2-10GS-I3-01v9-L7P/aluminum phosphate (AP) at w0 and w3 via intradermal (i.d.) footpad injections (0.8 μg per injection site, totaling 3.3 μg per mouse). The spike probe in fig. S3A (left), S2GΔHR2-5GS-foldon-Avi-Biot, was used to sort mouse LN B cells. (**A**) Bulk B-cell sorting experiment. Top: gating strategies used in the antigen-specific sorting of mouse LB B cells (Step 1: remove cell debris; Steps 2 and 3: exclude clumped or sticky cells to ensure that only single cells remain; Step 4: remove dead cells; Step 5: identify antigen-specific B cells). Bottom: Summary of SARS-CoV-2 spike-specific bulk sorting of mouse LN B cells from three vaccine groups. Bottom: Summary of SARS-CoV-2 spike-specific bulk sorting of mouse LN B cells from three vaccine groups. (**B**) Antibodyomics analysis of repertoire NGS data obtained for spike-specific mouse LN B cells from three vaccine groups. B-cell repertoire profiles are shown for three groups of mice immunized with (**C**) S2GΔHR2-5GS-1TD0, (**D**) S2GΔHR2-5GS-E2p-L4P, and (**E**) S2GΔHR2-10GS-I3-01v9-L7P. Top: VH gene usage (left) and VK gene usage (right); Bottom: germline VH/VK divergence (left) and CDRH/K3 loop length (right). (**F**) Cross-group comparison of key repertoire properties. Left: statistical comparison of the number of activated VH/VK genes (≥ 1% of the total population); middle: statistical comparison of the average VH/VK SHM rate; right: statistical comparison of HCDR3 loop length and root-mean-square fluctuation (RMSF) of HCDR3 loop length. The RMSF value is used as an indicator of how much HCDR3 loop length varies within the spike-specific antibodies from each mouse. In statistical comparison, mean value and standard deviation (SD) are shown as black lines. *P*-values were determined by an unpaired two-tailed *t* test in GraphPad Prism 9.1.2. The asterisk symbol (*) indicates the level of statistical significance: ns (not significant), *$p < 0.05$, **$p < 0.01$. Of note, for M4 of the S2GΔHR2-5GS-1TD0 group, only 39 reads were obtained from Ion S5 sequencing due to the difficulty in light chain library preparation, as shown in the summary in (B). Therefore, the light chain profiles for M4 are only shown for the sake of completeness in (C) and will not be included in the comparison between different vaccine groups in (F).
